# Predicting Affinity Through Homology (PATH): Interpretable Binding Affinity Prediction with Persistent Homology

**DOI:** 10.1101/2023.11.16.567384

**Authors:** Yuxi Long, Bruce R. Donald

## Abstract

Accurate binding affinity prediction is crucial to structure-based drug design. Recent work used computational topology to obtain an effective representation of protein-ligand interactions. While algorithms using algebraic topology have proven useful in predicting properties of biomolecules, previous algorithms employed uninterpretable machine learning models which failed to explain the underlying geometric and topological features that drive accurate binding affinity prediction. Moreover, they had high computational complexity which made them intractable for large proteins.

We present the fastest known algorithm to compute persistent homology features for protein-ligand complexes using opposition distance, with a runtime that is independent of the protein size. Then, we exploit these features in a novel, interpretable algorithm to predict protein-ligand binding affinity. Our algorithm achieves interpretability through an effective embedding of distances across bipartite matchings of the protein and ligand atoms into real-valued functions by summing Gaussians centered at features constructed by persistent homology. We name these functions *internuclear persistent contours (IPCs)*. Next, we introduce *persistence fingerprints*, a vector with 10 components that sketches the distances of different bipartite matching between protein and ligand atoms, refined from IPCs. Let the number of protein atoms in the protein-ligand complex be *n*, number of ligand atoms be *m*, and *ω* ≈ 2.4 be the matrix multiplication exponent. We show that for any 0 *< ε <* 1, after an 𝒪 (*mn* log(*mn*)) preprocessing procedure, we can compute an *ε*-accurate approximation to the persistence fingerprint in 𝒪 (*m* log^6*ω*^(*m/ε*)) time, independent of protein size. This is an improvement in time complexity by a factor of 𝒪 ((*m*+ *n*)^3^) over any previous binding affinity prediction that uses persistent homology. We show that the representational power of persistence fingerprint generalizes to protein-ligand binding datasets beyond the training dataset. Then, we introduce *PATH*, Predicting Affinity Through Homology, a two-part algorithm consisting of PATH^+^ and PATH^−^. PATH^+^ is an interpretable, small ensemble of shallow regression trees for binding affinity prediction from persistence fingerprints. We show that despite using 1,400-fold fewer features, PATH^+^ has comparable performance to a previous state-of-the-art binding affinity prediction algorithm that uses persistent homology. Moreover, PATH^+^ has the advantage of being interpretable. We visualize the features captured by persistence fingerprint for variant HIV-1 protease complexes and show that persistence fingerprint captures binding-relevant structural mutations. PATH^−^, in turn, uses regression trees over IPCs to differentiate between binding and decoy complexes. Finally, we benchmarked PATH versus established binding affinity prediction algorithms spanning physics-based, knowledge-based, and deep learning methods, revealing that PATH has comparable or better performance with less overfitting, compared to these state-of-the-art methods. The source code for PATH is released open-source as part of the osprey protein design software package.

## 1 Introduction

Structure-based drug design (SBDD) is an invaluable tool for effective lead discovery [8]. An important step in SBDD is virtual screening, where large libraries of compounds are computationally screened against a protein target of known structure to predict inhibitors that bind to the target [98]. SBDD is enabled by docking, where a library of small compounds is computationally docked into the binding site of a target protein, and the most promising compounds are selected for further evaluation [50, 69]. A reliable ranking of docking poses, based on affinity, potency, or other biophysical properties, is essential for accurate SBDD [69].

Binding affinity characterizes the strength of the interaction between a protein and a ligand. Binding affinity is also a key factor in determining the efficacy of a drug, as tight-binding ligands are more likely to be potent drugs *in vivo* [96, 56, 92]. Since experimental determination of protein-ligand binding affinity is time-consuming and costly in many cases [121], accurate methods for protein-ligand binding affinity prediction *in silico* are crucial in the structure-based drug design process [8, 6]. One approach to *in silico* binding affinity prediction is through molecular dynamics simulations [7, 4]. Unfortunately, while molecular dynamics is rigorous and accurate in predictions, it is computationally intensive [77] and is not suitable for virtual screening, where a large number of compounds must be screened. Therefore, many scoring functions [57, 74] have been developed, including physics-based, regression-based, and knowledge-based scoring functions [74]. Recent years also saw the application of many deep learning (DL) techniques for binding affinity prediction, including convolutional neural networks [45, 105, 100], attention mechanism [96, 44], and graph neural networks [115, 55]. These deep learning methods have produced more accurate predictions than handcrafted scoring functions.

Beside accuracy, *interpretability* is an essential quality and a key factor of trust for an algorithm. A non-interpretable model, also called a *black box* model, can “predict the right answer for the wrong reason” [94]. As a result, it is questionable whether a black box model could generalize beyond the training dataset [94]. A prime example of this caveat is AlphaFold2 [46], a black box model that achieved near-experimental accuracy on the CASP14 protein folding challenge [52] but is unable to predict the impact of structure-disrupting mutations, which are frequently associated with protein aggregation, misfolding, and dysfunction [11, 83]. The inability to understand the underlying workings of a black box model means that such solecisms are hard to discover in advance and their causes can only be speculated. On the other hand, an interpretable model is more robust, since the model can be calibrated to adapt to scenarios or considerations outside of its training dataset [93], such as mutant proteins, which were not abundantly present in the PDB dataset. As a result, when a discrepancy between an interpretable algorithm and empirical measurements is discovered, its cause can be precisely identified and fixed. Furthermore, interpretations from a model can produce insights, contribute to the understanding of the underlying system [80], and facilitate development for further efficient algorithms. While physics-based scoring functions for binding affinity usually have some interpretability [74], the adoption of deep learning techniques poses an inherent challenge in interpretability in DL based algorithms [93].

We propose a novel, interpretable algorithm for protein-ligand binding affinity prediction. Our algorithm achieves interpretability by an effective embedding of protein-ligand interactions in a low-dimensional vector through *persistent homology*. Persistent homology quantifies the shapes of protein-ligand complexes by computing the *persistence* of topological invariants like holes and voids in the biomolecular structures at different spatial resolutions [31]. We show that the features encoded by persistent homology can be both highly descriptive and interpretable (Section 2.1). The information captured by persistent homology from a subset of atoms can be faithfully represented in a *persistence diagram*, which is a collection of points (*x, y*) in ℝ^2^ where the *x*- and *y*-coordinates respectively represent the birth and death radii of a homology group in a filtration. Algorithms using persistent homology have been applied to neuronal morphologies [47] and protein cavity detection [28, 58]. The TNet-BP algorithm in the TopologyNet family of algorithms [13] captures features with persistent homology and uses a convolutional neural network to predict binding affinity of protein-ligand complexes without auxiliary features constructed from other sources. Persistent homology has been used in several algorithms for binding affinity prediction, where persistent homology features are combined with chemical features, and the prediction is made using a neural network [14, 12, 112, 106, 110, 62]. Persistent homology shows promise as an approach for accurate binding affinity prediction, and an algorithm using persistent homology and deep learning won many challenges in the D3R Grand Challenge 3 [82, 36].

For representing molecular features, persistent homology provides two advantages that correlate to known properties of biomolecules:

### 1. Stability with respect to noise

It has been proven that small changes in the input to persistent homology (measured by *bottleneck distance*) cause only small changes in the persistence diagrams (measured by *Gromov-Hausdorff distance*) [19, 31], which are representations of persistent homology. Thus, in embedding biochemical structure, small differences in the protein structure that arise often due to the structural heterogeneity of proteins [25] will have little effect on the resulting representation.

### 2. Invariance under translation and rotation

The persistence diagram representation is invariant under translation and rotation of the biomolecule. This corresponds to the physical fact that the structure of a protein is not affected by the rigid-body translation or rotation of the protein in space.

However, while the features encoded by persistent homology have geometric relevance, the TNet-BP algorithm [13] uses a convolutional neural network to predict binding affinity from these features. This makes it difficult to interpret the model or results. Additionally, the authors of TNet-BP introduce a novel distance function, the *opposition distance* (Section 2.2). While the prediction accuracy was high, there has been little interpretation of the features captured through persistent homology and the prediction algorithms have been black box algorithms, even among later works [12, 106]. Overall, black box predictions of high stakes targets have been difficult to deploy and reduce to practice [93]. To overcome these limitations and develop a machine learning algorithm which is both interpretable and comparably accurate, we describe a one-dimensional representation of the features captured by persistent homology constructed with opposition distance, which we term *internuclear persistence contour* (IPC). From IPCs, we introduce a new representation for protein-ligand interactions, *persistence fingerprints*, which are low dimensional feature subsets that were iteratively refined from a set of persistent homology features inspired by the TNet-BP algorithm [13] and encoded using IPCs. Let the number of protein atoms be *n*, number of ligand atoms be *m*, and *ω* ≈ 2.4 be the matrix multiplication exponent. Previous binding affinity prediction algorithms that use persistent homology all have computational complexity 𝒪 ((*m* + *n*)^4.8^) or worse [106, 13, 12, 110, 62, 63, 61, 64]. However, in many biologically relevant protein-ligand complexes [17, 27, 42], *n* can be very large, resulting in unwieldy runtime and space consumption by these algorithms. We will show that for any 0 *< ε <* 1, after an 𝒪 (*mn* log(*mn*)) preprocessing procedure, we can compute an approximation to the persistence fingerprint in 𝒪 (*m* log^6*ω*^(*m/ε*)) time, independent of protein size, such that the maximum difference between each component in this approximation and that of the true persistence fingerprint is less than *ε*. Our ability to create a provably accurate approximation algorithm is primarily due to the small number of features that are employed in the persistence fingerprint.

PATH^+^ uses a small ensemble of shallow regression trees to predict binding affinity from persistence fingerprints. A *decision tree* is a predictive model represented by a rooted tree, where each internal node represents a test on an attribute, each branch represents the outcome of the test, and each leaf node represents a discrete target class label [89]. Decision trees are easily interpretable [67], and as a result have been broadly useful in many applications [117, 113]. Predictions of individual trees can be easily made by evaluating the tree manually, but the interpretability decreases with the depth of each tree and the number of trees in the ensemble. A *regression tree* is a decision tree whose target values are continuous variables. PATH^+^ and PATH^−^ are both *gradient boosting regressors (GBRs)*, which are ensembles of regression trees iteratively built up based on the error of the previous iteration. Gradient boosting regressors have been used for protein solvent accessibility [32], protein interactions [120], and predicting protein–RNA binding hot spots [24].

By presenting PATH, our paper makes the following contributions:

1. *Internuclear persistence contours (IPCs)*, real-valued functions that represent the number of different bipartite matchings between the protein and ligand atoms at different resolutions captured by persistent homology. An interpretation of IPCs reveals that PATH^+^ captures the bipartite matching between the protein and ligand at different scales to predict binding affinity (Section 3.1). This interpretation also provides insights on the features that other persistent homology-based binding affinity predictions algorithms [13, 12, 106] potentially capture.
2. A feature subset refined from IPCs which we term *persistence fingerprint* (Section 3.2). Validation of the generalization ability of persistence fingerprints on two large protein-ligand binding databases (Binding MOAD [43, 3, 99, 103] and BindingDB [37, 60]), which are disjoint from the database whence we curated persistence fingerprint (PDBBind [107, 66]), shows that persistence fingerprints can accurately predict binding affinity (Section 4).
3. A fast approximation algorithm for persistence fingerprint. Let number of protein atoms be *n*, number of ligand atoms be *m*, and *ω* ≈ 2.4 be the matrix multiplication exponent [53]. We show that for any 0 *< ε <* 1, after an 𝒪 (*mn* log(*mn*)) preprocessing procedure, one can compute an approximation to the persistence fingerprint in *𝒪* (*m* log^6*ω*^(*m/ε*)) time, independent of protein size, such that the maximum difference between each component in this approximation and that of the true persistence fingerprint is less than *ε*. This is an improvement in time complexity by a factor of 𝒪 ((*m* + *n*)^3^) over any previous binding affinity prediction that uses persistent homology (Theorem 1).
4. Two algorithms, PATH^+^ and PATH^−^, which together make up *PATH*, Predicting Affinity Through Homology. PATH^+^ uses persistence fingerprints to interpretably predict binding affinity. To our knowledge, PATH^+^ is the first *interpretable* algorithm that uses persistent homology to predict binding affinity [106, 13, 12, 110, 62, 63, 61, 64] (Section 3.3).
5. Validation of PATH^+^ on a held-out subset of the PDBBind v2020 refined set (519 protein-ligand complexes) showing that PATH^+^ achieves comparable performance with TNet-BP [13], a previous uninterpretable deep learning-based binding affinity prediction algorithm that uses persistent homology (Section 4). Compared to TNet-BP, PATH^+^ uses 1,400-fold fewer features, employing a fully interpretable model, achieving an accuracy better than an open-source implementation of TNet-BP, and an accuracy only 7% less (in RMSE) than a closed-source version – only 8% the energy of a hydrogen bond.
6. Comparison of PATH^+^ versus established scoring functions spanning physics-based, knowledge-based, and deep learning-based methods on Binding MOAD and BindingDB demonstrating that PATH^+^ shows comparable or better performance with less overfitting. Comparison of PATH^−^ on a held-out subset of the DUD-E [81] dataset versus established scoring functions showing that PATH^−^ has state-of-the-art performance on differentiating between decoys and active compounds.
7. Visualization of the features captured by persistence fingerprint on mutant HIV-1 proteases bound to the inhibitor darunavir (Section 4).
8. A free, open-source implementation of PATH in the computational protein software suite osprey [40] at https://github.com/donaldlab/OSPREY3.

## 2 Background

### 2.1 Introduction to Persistent Homology

The *persistent homology transform* of a geometric complex such as a protein-ligand structure is a cosheaf of combinatorial persistence diagrams. Specifically, the persistent homology of a point cloud is obtained as the composition of the birth-death functor and the Möbius inversion functor [33]. In operational terms, persistent homology takes a point cloud with a distance function and computes the *persistence* of “holes” of different dimensions at different spatial resolutions. In the case of protein-ligand complexes, the point cloud consists of the centers of all the protein and ligand atoms (e.g., from the Protein Data Bank [20]), and the pairwise distance is usually the Euclidean distance or in our case, the opposition distance (Eq. (2)). There are different *filtration functions* from which persistence fingerprints can be constructed. We describe persistent homology using the Vietoris-Rips filtration, which is employed to construct persistence fingerprints. (We invite the interested referee to find a formal definition of persistent homology in Appendix Section A.)

A simplex is the generalization of a filled triangle to higher dimensions. A 0-simplex is a point, a 1-simplex is a line segment, a 2-simplex is a triangle, and so on. Given a point cloud *S* and a pairwise distance matrix, we can define the Vietoris-Rips complex built on this point cloud with radius *r* ∈ [0, ∞) by constructing a simplex for any set of points *σ* whose pairwise distance is at most *r* [122]. Let **VR**_*r*_(*S*) denote the Vietoris-Rips complex built on *S* with radius *r*, then

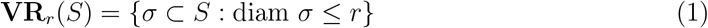

where diam *σ* denotes the the supremum over the distances between the points in *σ*.

Geometrically, the rank of the *n*^th^ homology group measures a topological invariant: the number of *n*-dimensional holes in the simplicial complex. For example, the 0^th^ homology group measures the number of connected components, the 1^st^ homology group measures the number of loops, and the 2^nd^ homology group measures the number of voids. In persistent homology, a sequence of simplicial complexes is built up with respect to an increasing *filtration parameter*, which is the radius *r* in the case of Vietoris-Rips complex described above, and change in rank of the homology group (i.e., the appearance or disappearance of copies of ℤ in the direct sum generating the homology group) is measured [13, 5]. Geometrically, persistent homology measures the appearance and disappearance of these topological invariants (Appendix Figure 6).

One way to represent the persistence of these topological invariants is by *persistence diagrams*. Persistence diagrams represent the birth and death of each invariant as a point (*x, y*), where *x* is the filtration parameter at which the invariant appears and *y* is the filtration parameter at which the invariant disappears [1]. Since there can be varying numbers of topological invariants, a vectorization technique is used to convert persistence diagrams to fixed size vectors to employ machine learning techniques, such as support vector machine, decision tree, and neural networks [1]. All of these require a fixed size input. In the construction of persistence fingerprints, we constructed *internuclear persistence contours (IPCs)*, which is a special case of *persistence images* (see Section 3.1 for a detailed explanation of IPCs. For the interested referee, Appendix Section A.2 has the definition of persistence images.) to vectorize the persistence diagrams. Persistence images are provably stable with respect to input noise [1].

**Table 1.** Comparison of PATH^+^ with TNet-BP [13]. PATH^+^ uses a much lower dimensional input vector than TNet-BP, which allowed the use of an interpretable regression model.

|  | **TNet-BP [13]** | **PATH<sup>+</sup> (this paper)** |
| --- | --- | --- |
| **Dimensionality reduction method** | Persistence diagrams constructed using opposition distance and Euclidean distance on 40 subsets of atoms | Internuclear persistence contours (IPCs) |
| **Input vector to regressor** | Binned persistence diagrams | Persistence fingerprints refined from IPCs |
| **Input vector dimension** | 14,400 | 10 |
| **Regressor Algorithm** | Convolutional neural network | Sparse ensemble of regression trees |
| **Interpretation?** | No | Yes |

**Table 2.** The 10 features captured by persistence fingerprint. The source of each feature is represented by a 4-tuple, consisting of ^**a**^the element of protein atoms used in the IPC, ^**b**^the element of ligand atoms used in the IPC, ^**c**^the dimension of the homology group where the IPC is derived from, and ^**d**^the interval (or bin) where the IPC is integrated over to yield the value of this feature. For example, the first row describes that the first component of the vector is calculated by integrating an IPC over the interval [9.5, 10.0], where the IPC is constructed using the carbon atoms of the protein and the carbon atoms of the ligand, and the persistence of homology groups of dimension 1 is measured.

| Protein Atom<sup>a</sup> | Ligand Atom<sup>b</sup> | IPC Dimension<sup>c</sup> | IPC Density Bin<sup>d</sup> (Å) | Protein Atom<sup>a</sup> | Ligand Atom<sup>b</sup> | IPC Dimension<sup>c</sup> | IPC Density Bin<sup>d</sup> (Å) |
| --- | --- | --- | --- | --- | --- | --- | --- |
| C | C | 1 | [9.5, 10.0] | N | C | 1 | [8.0, 8.5] |
| C | C | 1 | [9.0, 9.5] | C | N | 0 | [7.5, 8.0] |
| C | C | 1 | [7.0, 7.5] | C | N | 0 | [8.5, 9.0] |
| C | C | 1 | [4.0, 4.5] | C | O | 0 | [6.5, 7.0] |
| N | C | 1 | [10.0, 10.5] | C | S | 0 | [5.0, 5.5] |

### 2.2 The TNet-BP algorithm [13]

TNet-BP from the TopologyNet family of algorithms [13] is a previous persistent homology-based algorithm for protein-ligand binding affinity prediction. TNet-BP [13] introduced the *opposition distance* (*d*_*op*_) between two atoms *a*_*i*_ and *a*_*j*_ as follows:

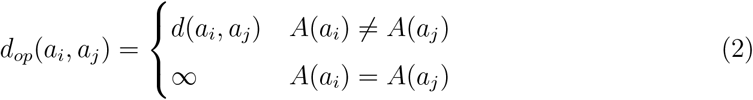

where *d*(·, ·) is the Euclidean distance between two atoms and *A*(·) denotes the *affiliation* of an atom, which is either a protein or a ligand. Note that opposition distance does not satisfy triangle inequality and does not have a clear interpretation by itself. Rather, opposition distance works together with the construction of Vietoris-Rips complexes and persistent homology to capture bipartite matching of protein and ligand atoms at different scales. TNet-BP employed 36 persistent homology diagrams with 36 different subsets of atoms using opposition distance and 4 additional persistent homology diagrams using Euclidean distance. The center of each atom (in 3D) is used as its position in the point cloud.

To vectorize each persistence diagram, TNet-BP [13] created a 200 *×* 72 array where every row represents the births, deaths and persistences of features in one dimension of a persistence diagram. Each row consists of 200 bins, and the value of each bin is the number of features in the persistence diagram that fall into that bin. Finally, to predict binding affinity, TNet-BP used a convolutional neural network on this vector representation. This algorithm was trained and tested on the PDBBind v2020 refined set, a dataset curated from the Protein Data Bank [20] with protein-ligand complexes with their experimental binding affinities [107, 57, 65, 66].

## 3 Design of PATH

### 3.1 Internuclear Persistence Contours (IPCs)

Through feature selection (detailed in Appendix Section B for the interested referee), we found that only the 0- and 1-dimensional persistence diagrams constructed with the opposition distance (*d*_*op*_) are necessary for accurate binding affinity prediction. We make the following observations about the Vietoris-Rips filtration constructed with the opposition distance:

1. The 0D homology groups all have birth radius 0. This can be seen by the interpretation of 0D homology groups as the number of connected components: as the filtration radius increases, the number of connected components can only decrease. Given a protein atom and a ligand atom, the appearance of a 0D hole happens when the filtration radius *r* equals 0 and disappearance of a 0D hole happens when *r* equals the distance between these two atoms.
2. The 1D holes all have death radius ∞. The birth radius of a 1-dimensional hole corresponds to distances of bipartite matchings of the protein and ligand atoms (For the interested reviewer, Lemmas 1 and 2 in the appendix prove this).

Note that each feature captured by persistent homology with opposition distance involves the distance between a protein and a ligand atom, hence the similarity with bipartite graphs. Also note that the persistence of each 0D and 1D homology group constructed with opposition distance can be captured using only a single scalar value (death time for 0D, birth time for 1D), rather than a two-dimensional vector. We term this scalar value the *critical value* of persistence.

This allows us to introduce a new representation of the persistence diagrams constructed with opposition distance. An IPC is a function *γ* : ℝ → [0, ∞). Given a 0- or 1-dimensional persistent homology constructed with the opposition distance, we can construct its corresponding IPC by summing Gaussians of a given standard deviation centered at each of its critical values. A precise definition is given in Appendix Section A.3 for the interested referee. In our paper, the standard deviation of the Gaussian is chosen to be 0.1 Å.

IPCs can be discretized by taking the integral of IPC over bins of a fixed width. We call the value of the integral over each bin *IPC density* and the resulting collection of IPC densities the *discretized IPC*. The discretized IPC is a nonnegative real-valued function with respect to the bins. Discretized IPCs are closely related to persistence images [1], but discretized IPCs are derived only from persistent homology whose information can be encoded in one dimension (such as when constructed with opposition distance), as opposed to persistence images which can be derived from any persistent homology construction.

### 3.2 Persistence Fingerprint

For a given protein-ligand complex, we first constructed 36 pairs of IPCs (0D and 1D) from the 36 subsets of atoms used to construct opposition distance-based persistence diagrams in TNet-BP [13]. Then, we selected the most important components in the IPCs for binding affinity prediction by constructing gradient boosting regressors (GBRs) on the IPCs, identifying the most important features measured by their mean decrease in impurity [86, 75] and through an iterative feature ablation procedure [76] (A detailed account of the iterative ablation procedure is given in Appendix Section B for the interested referee). We found that 10 specific components from the discretized IPCs suffice to produce a binding affinity prediction model with comparable performance to TNet-BP. We call the vector made up of these 10 components the *persistence fingerprint*.

#### Theorem 1

**(Complexity of approximating persistence fingerprint).** *Assume there exists a fixed lower bound on interatomic distances in a protein-ligand complex. Let the number of protein atoms be n, the number of ligand atoms be m, and ω* ≈ 2.4 *be the matrix multiplication exponent [53]. For any* 0 *< ε <* 1, *after an O*(*mn* log(*mn*)) *preprocessing procedure, we can compute an approximation to the persistence fingerprint in* 𝒪 (*m* log^6*ω*^(*m/ε*)) *time, independent of protein size, such that the maximum difference between each component in this approximation and that of the corresponding element in the true persistence fingerprint is less than ε*.

Proof of Theorem 1 relies on the choice of a weight function for IPCs that decays exponentially, such as the Gaussian. This leads to convergence of any persistence fingerprint component on a ligand atom *l* for a protein of any size. Then for any *ε*, there exists a radius *r*_*ε*_ such that removing all atoms further than *r*_*ε*_ from *l* yields an approximation that is at least *ε*-accurate. The interested reviewer can find a full proof in Appendix Section C. Additionally, an empirical evaluation on a subset of the BioLiP dataset [114] with 45,199 protein-ligand complexes corroborated our asymptotic runtime analysis and achieved an average runtime of 41.4 seconds to calculate the persistence fingerprint of a protein-ligand complex and a maximum approximation error of *ε* = 4.8 *×* 10^−7^, where *ε* is defined as in Theorem 1 (The interested referee can find details in Appendix Section C.4).

### 3.3 PATH^+^

While GBRs have been used as regressors in previous persistent homology based binding affinity prediction algorithms, previous models used 20,000 trees [13, 12, 14, 106] and the large number of trees makes these models impossible to interpret. In comparison, PATH^+^ has only 13 regression trees, which is three orders of magnitude fewer than previous algorithms, all while maintaining a comparable performance (Figure 3). The number of trees, tree depth, and learning rate of the GBRs in PATH^+^ were selected after we measured the performance of GBRs with respect to these three parameters (For the interested referee, Figure 11 in the appendix shows ablation results with respect to number of trees) and balanced performance (Figure 3) with interpretability (Table 3 shows a comparison between PATH^+^ and TNet-BP, and for the interested referee, Table 5 in the appendix shows the performance of GBRs with larger inputs and more trees. Additional experiments on alternative regression methods in Appendix Section B.4 confirms that trees-based regressors are optimal on persistence fingerprint). The simplicity of regression trees highlights the representational power of persistence fingerprints. (For the interested referee, the precise set of decision trees in PATH^+^ can be found in Appendix Section D.1.)

**Table 3.**
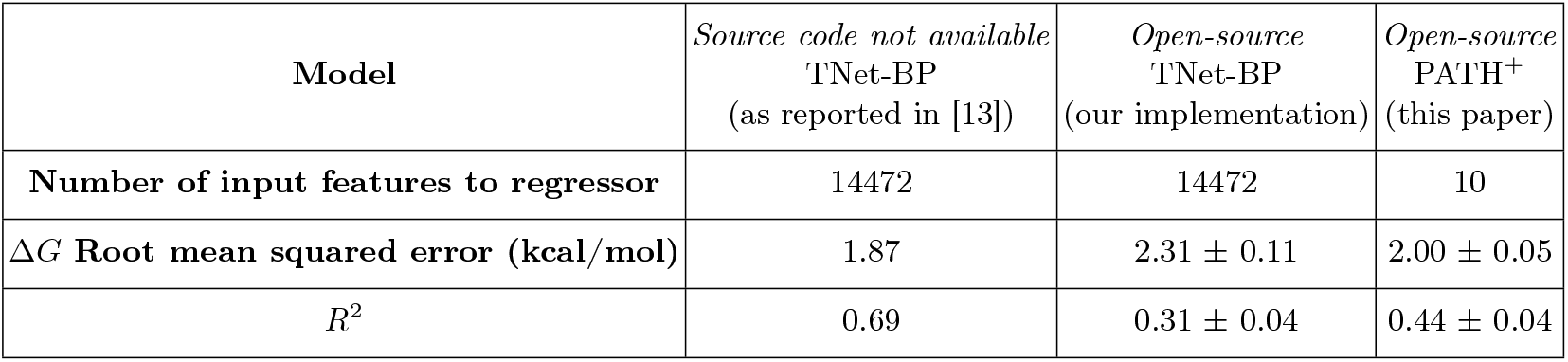
Performance of TNet-BP and PATH^+^ on the PDBBind v2020 refined set shows that PATH^+^ achieves similar performance with TNet-BP while having three orders fewer features. For each of the algorithms we have implemented, mean *±* standard deviation is reported for root mean squared error (RMSE) and *R*^2^ between predicted Δ*G* and experimental Δ*G* over 100 random restarts. The RMSE is comparable in magnitude to a hydrogen bond with the atoms N–H … O, which has a molar energy of about 1.9 kcal/mol [22]. We believe that the modest *R*^2^ for PATH^+^ is due to the fact that the small number of trees means that PATH^+^ predicts a relatively small number of discrete values, which hurts the *R*^2^ metric of PATH^+^ because *R*^2^ tracks small differences in predictions.

**Fig. 1.**
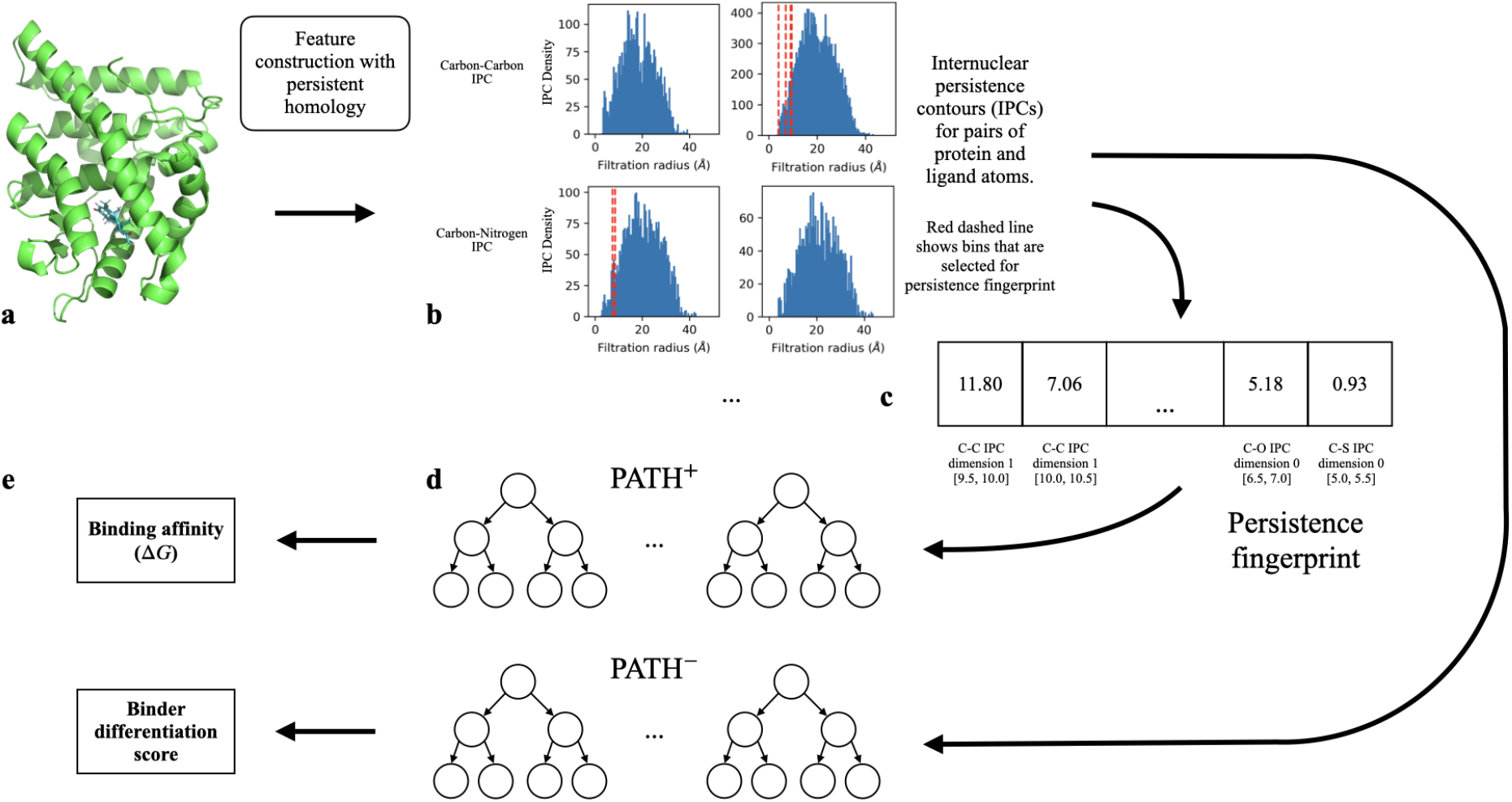
An overview of the PATH algorithm. ^**a**^The input is a 3D structure of the protein-ligand complex. Shown as example: humanised monomeric RadA in complex with indazole (PDB ID: 4b2i [95]). ^**b**^*Internuclear persistence contours (IPCs) are* constructed using persistent homology. Two sets of IPCs are shown as examples. 10 specific IPC densities (shown in red dotted lines in ^**b**^) are collected into ^**c**^persistence fingerprint. The features that each component of persistence fingerprint encodes are detailed in Table 2 and the procedure by which these features are selected is detailed in Appendix B for the interested referee. ^**d**, **e**^Binding affinity is predicted using a small ensemble (10) of regression trees from the persistence fingerprint in PATH^+^, and a score for differentiating between binders and non-binders is predicted using an ensemble of trees from the IPCs in PATH^−^.

**Fig. 2.**
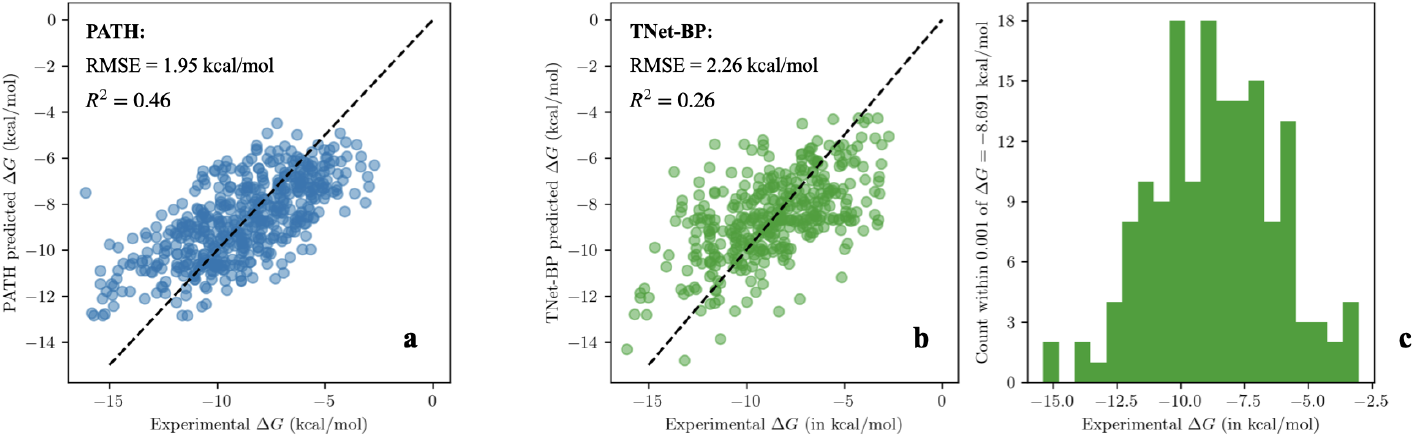
Scatter plots of PATH^+^ and our implementation of TNet-BP’s predictions on a held-out, test subset of PDBBind v2020 refined set for one run (90:10 train:test split ratio, *n*_test_=519) shows that PATH^+^ produces better predictions, especially on protein-ligand complexes whose binding affinity that deviate significantly from the mean. This highlights PATH^+^’s generalizability. ^**a**^Predictions of PATH^+^: *R*^2^=0.46, *RMSE*=1.95 kcal/mol ^**b**^Predictions of TNet-BP: *R*^2^=0.26, *RMSE*=2.26 kcal/mol. ^**c**^To declutter the TNet-BP scatter plot in ^**b**^, we removed 142 data points that are all predicted to have Δ*G* within 0.001 kcal/mol of −8.691 kcal/mol by TNet-BP, and instead show the distribution of these points on a separate histogram. The 1-run performances of each algorithm in ^**a,b**^ are very close to their average performances over 100 runs in Table 3.

**Fig. 3.**
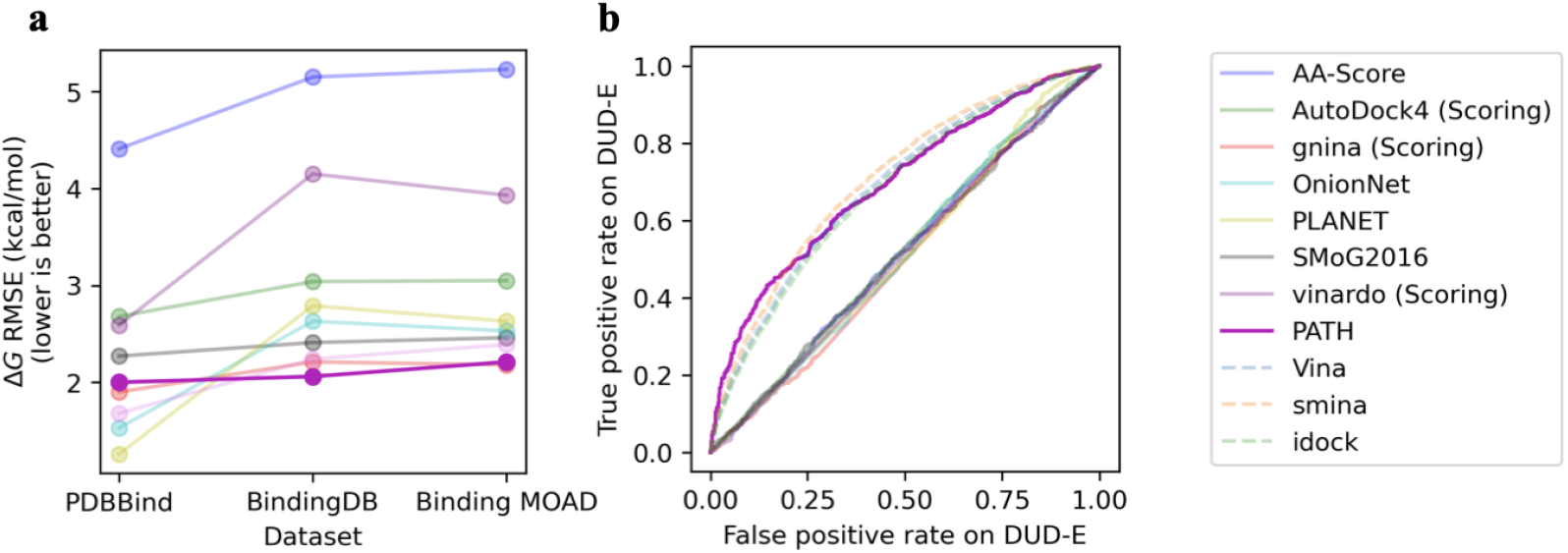
PATH has state-of-the-art performance versus previous binding affinity prediction algorithms. ^**a**^PATH^+^ shows comparable or better performance with less overfitting, as evidenced by a smaller slope, with much less increase in Δ*G* RMSEs beyond the training dataset, compared to established binding affinity prediction algorithms spanning a variety of methods. The benchmarked algorithms include physics-based and deep learning algorithms from the famous AutoDock framework (scoring function of AutoDock4 implemented in the AutoDockFR package [79, 91], Vinardo [90], GNINA [73]), empirical (AA-Score [84]), knowledge-based (SMoG2016 [23]), and deep learning-based scoring functions (OnionNet [119], PLANET [118]). We believe that PATH^+^ overfit far less to training dataset than other methods due to the small number of parameters in the sprase regression trees of PATH^+. **b**^ROC curves of scoring functions benchmarked on the DUD-E dataset show PATH^−^ has state-of-the-art performance in discriminating decoys in the DUD-E dataset. AutoDock4, gnina, and vinardo are all benchmarked as scoring functions. We also plot interpolated ROC curves (dashed) based on AUCs from [71] which benchmarked Vina [29], smina [49], and idock [54] using the full AutoDock framework. The only algorithms with non-diagonal ROCs are PATH^−^ (AUC=0.696), and the three scoring functions tested with the full AutoDock framework: Vina (AUC=0.69), Smina (AUC=0.71), and Idock (AUC=0.68). (The interested reviewer can find the full results in numeral tables in Appendix E.)

### 3.4 PATH^−^

Based on the intuition that binding is a mostly local interaction hence mostly driven by local structures, and the observation that persistence fingerprints from PATH^+^ (Table 2) all have IPC radius less than 11 Å, we hypothesize that the components of the discretized IPCs that are important to distinguishing between active and decoy compounds are also within a certain radius of the ligand. Therefore, to construct PATH^−^, we trained a gradient boosting regressor on the 36 pairs of discretized IPCs, the same set that was used in persistence fingerprint of PATH^+^ (Section 3.2), constructed from 551 active and 20227 decoy compounds with 70 proteins from the DUD-E dataset, where only protein atoms within 15 Å from the ligand were used to construct the IPCs. Removal of atoms beyond 15 Å from computing IPCs yields extremely low error in computing the persistence fingerprint for PATH^+^; hence, we expect the error for important components in discretized IPC due to this approximation to be low as well. Using this approximation, PATH^−^ achieves the same fast 𝒪 (*mn* log(*mn*)) + 𝒪 (*m* log ^6*ω*^ (*m/ε*)) runtime as PATH^+^.

## 4 Computational Experiments and Results

First, we measured the performance of PATH^+^ versus TNet-BP [13] on the PDBBind v2020 refined set [107]. Due to the lack of a published source code for TNet-BP, we reimplemented TNet-BP’s persistent homology feature generation and neural network exactly as described in [13] and benchmarked it on the PDBBind v2020 refined set. Despite following [13] metic-ulously, we found that our implementation of TNet-BP performs worse than described in the original paper. We report both the performance of TNet-BP from our implementation and the results from the original paper in Table 3. Even when compared to the results of TNet-BP reported in the original paper [13], PATH^+^ sacrifices only a slight amount of accuracy in exchange for interpretability, an important characteristic that TNet-BP does not possess. Overall, these results show that PATH^+^ is an interpretable model using 1000-fold fewer features with only a 7% drop in accuracy compared to TNet-BP.

Next, We benchmarked PATH^+^ and PATH^−^ against established scoring functions for docking on the PDBBind, Binding MOAD, BindingDB (for binding affinity prediction of binders) and a subset of the DUD-E dataset (for decoy prediction). To avoid data leakage, this subset of the DUD-E dataset was chosen to be disjoint from the training dataset of PATH^−^. The scoring functions we tested include physics- and deep learning-based algorithms from the famous AutoDock framework (scoring function of AutoDock4 implemented in the AutoDockFR package [91], Vinardo [90], GNINA [73]), empirical (AA-Score [84]), knowledge-based (SMoG2016 [23]), and deep learning-based scoring functions (OnionNet [119], PLANET [118]). We note that the AutoDock framework has an advanced pose generation algorithm, which may filter out non-binding conformations before they reach the scoring functions. Therefore, for the decoy prediction task on the DUD-E dataset, we report not only the performance of AutoDock4, gnina, and Vinardo as standalone scoring functions, but also the performance of Vina, smina, and idock benchmarked on DUD-E using the entire AutoDock framework from [71]. Each scoring function was tested with 1 CPU core, 8GB of memory, and 1 hour of compute time. The RMSEs reported on the positive datasets for each algorithm were computed using the subset of protein-ligand complexes where that algorithm returned a prediction. The AUC of negative datasets were computed by considering the protein-ligand complexes where predictions were not returned as either all binders or all nonbinders, whichever yields a better AUC.

PATH^+^ achieved RMSEs of 2.00, 2.06, 2.21 in Δ*G* (kcal/mol) on PDBBind, BindingDB, and Binding MOAD respectively, which is a comparable or better performance with less over-fitting compared to the established binding affinity prediction algorithms we benchmarked (Figure 3). The error is comparable in magnitude to a hydrogen bond with the atoms N–H … O, which has a molar energy of about 1.9 kcal/mol [22]. PATH^−^ has an AUC of 0.696 on predicting decoys from the DUD-E subset disjoint from the training set of PATH^−^, outperforms the 7 binding affinity prediction algorithms, and performs similarly to the AutoDock algorithms when they are run with the entire AutoDock framework as reported in [71] (Figure 3).

Finally, to demonstrate the interpretability of PATH^+^, we inspected the persistence fingerprint and the atoms contributing to persistence fingerprint of two mutant HIV-1 proteases bound to the small molecule inhibitor darunavir. Figures 4 and 5 show the persistence fingerprint and atoms that contribute to a persistence fingerprint component of two mutant HIV-1 protease variants bound to a small molecule inhibitor darunavir (G48V: PDB ID 3cyw [59] & L90M: PDB ID 2f81 [51]). The structural changes induced by the mutation were captured by the persistence fingerprint (Figure 4). Figure 5 highlights a specific region of the protein-ligand complex, showing how a persistence fingerprint component changed from the structural difference. [51] observed a strong hydrogen bond (2.5 Å) to the carboxylate moiety of Asp30 shown in Figure 5, which correlates with the stronger contributions of backbone carbon atoms to persistence fingerprint in the L90M mutant than in the G48V mutant.

**Fig. 4.**
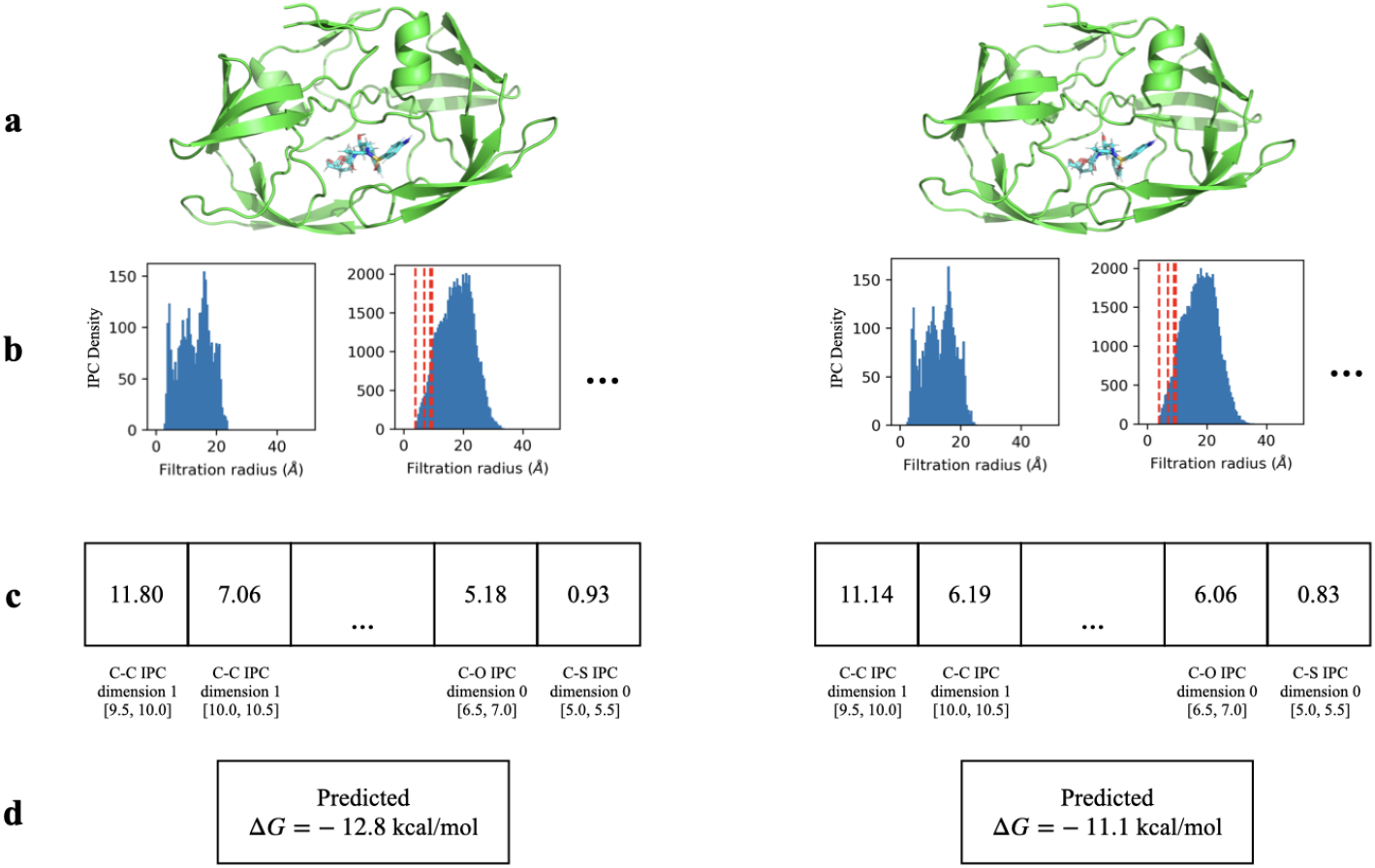
PATH^+^ correctly predicted a weaker binding affinity for HIV-1 protease with the drug-resistant G48V mutation (right, experimental Δ*G* = −10.6 kcal/mol, PDB ID: 3cyw [59]) bound to darunavir, compared to L90M HIV-1 protease (left, experimental Δ*G* = −14.35 kcal/mol, PDB ID: 2f81 [51]) complexed with the same inhibitor. ^**a**^The structure of each complex. ^**b**^The discretized internuclear persistence contour (IPC) of each complex. ^**c**^The persistence fingerprint of each complex. ^**d**^PATH^+^ correctly predicted a weaker binding affinity for the HIV-1 protease with G48V mutation.

**Fig. 5.**
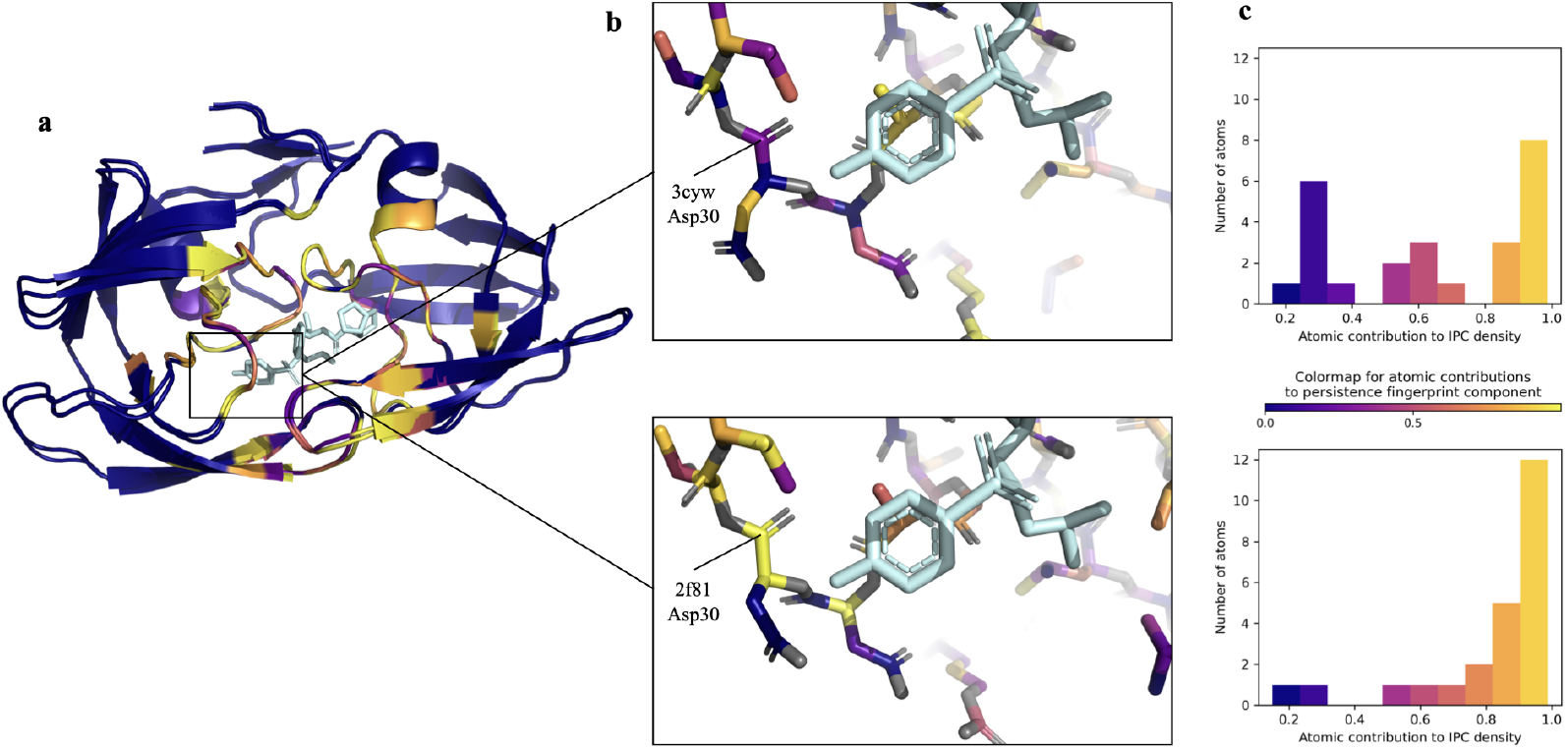
^**a**^Two HIV-1 protease mutants bound to inhibitor darunavir (G48V: PDB ID 3cyw [59] & L90M: PDB ID 2f81 [51]). Light blue: darunavir. The carbon atoms are colored by their individual contributions (blue through yellow, see legend) to the 2^nd^ component of persistence fingerprint (carbon-carbon IPC density at dimension 1 and bin [9.0, 9.5]). Grey: other protein heavy atoms. ^**b**^Detail of residues 27-32 for each protease with darunavir. Note change in conformation (and IPC densities) of Asp30. [51] observed a strong hydrogen bond (2.5 Å) to the carboxylate moiety of Asp30. This correlates to Asp30 of the L90M variant contributing highly to the persistence fingerprint component, which obtained a prediction of tighter binding affinity for L90M via the decision trees. ^**c**^Histograms of atomic contributions of residues 27-32 to the persistence fingerprint shows the carbon atoms of 2f81 in these residues had generally higher contributions to persistence fingerprint.

**Fig. 6.**
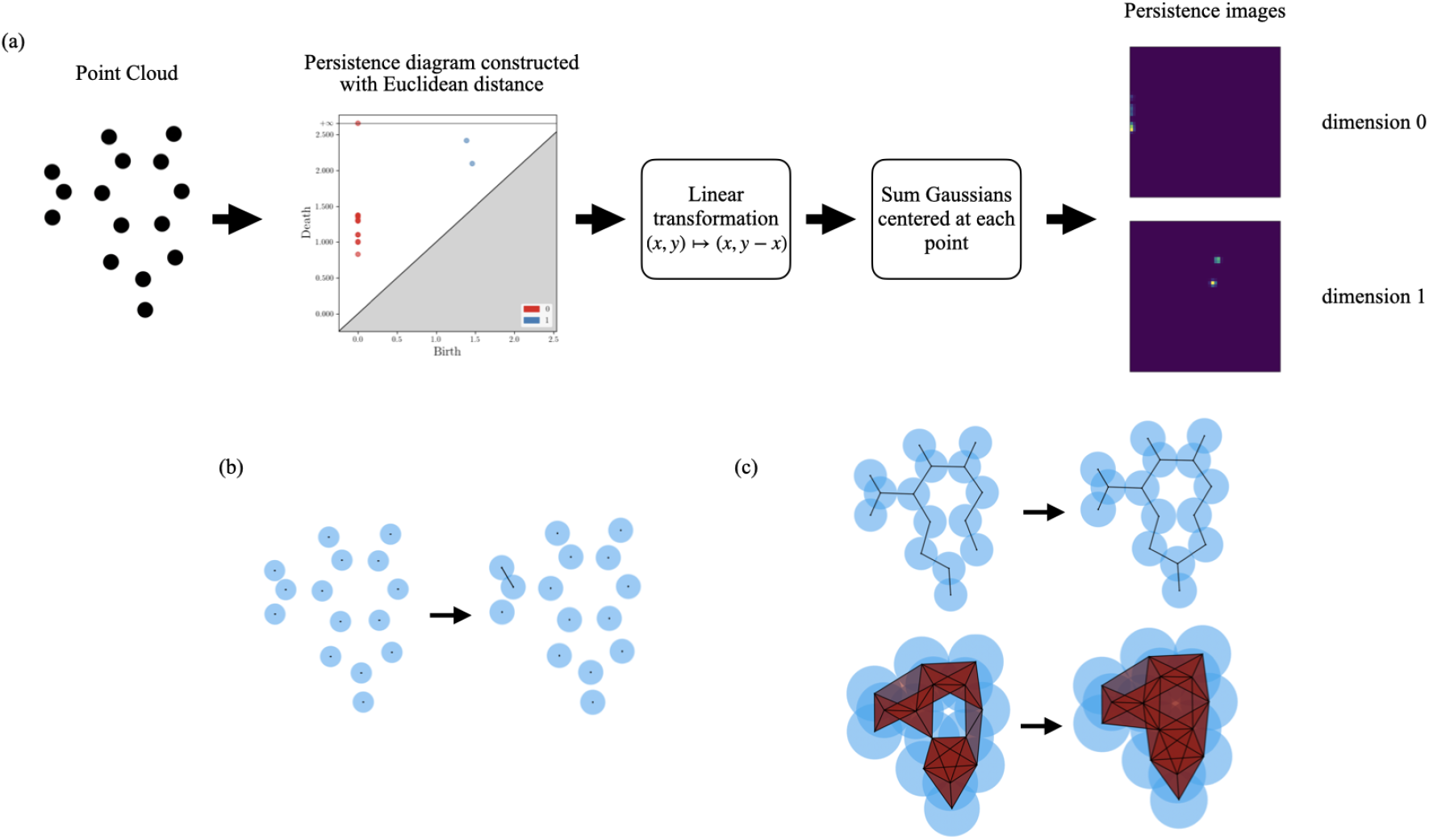
Construction of persistence images with the Euclidean distance. **(a)** Example construction of a persistence image using an adenine molecule and Euclidean distance as an example. Note two differences between this illustration and construction of persistence fingerprints (detailed in Appendix Section A.2). One, while this figure only shows persistence image constructed with the Euclidean distance, persistence fingerprint includes features constructed with *d*_*op*_, which is a different distance function (Table 2).Two, the adenine molecule is planar, hence can be shown in 2D, but the actual construction of persistence fingerprints are done in 3D. Details are left to Appendix Section A.2 **(b)** Disappearance of a connected component (i.e., zero-dimensional hole) at *r* = 0.8. Note that the disconnected components had existed since *r* = 0. These correspond to the point on the persistence diagram with coordinates (0, 0.8). **(c)** Appearance of a one-dimensional hole at *r* = 1.4 and disappearance at *r* = 2.5. This corresponds to the point on the persistence diagram with coordinates (1.4, 2.5).

## 5 Discussion

Persistent homology is a promising way to capture the structural information of a protein-ligand complex. The persistence of “holes” can capture bipartite matching of protein and ligand atoms at different spatial scales. Invariance to translation and rotation of its input and stability under small perturbation also make persistent homology advantageous for embedding biomolecules.

Our work highlights interpretability, a previously overlooked aspect in machine learning-based drug design, that persistent homology can bring to embedding biomolecules. The Vietoris-Rips filtration is such a simple construction that a person can effectively compute the persistent homology of a small point cloud by hand. As a result, the features captured by persistent homology can be accurately traced back to the precise atoms that constructed them. Despite the simplicity of their construction, features constructed by persistent homology are sufficient to produce a competent binding affinity prediction model. We also showed that in fact, only a tiny subset of features is needed to achieve comparable performance with previous works.

As opposed to the 14,472 features in TNet-BP, a previous work using persistent homology, our persistence fingerprint representation has only 10 features. Having three orders of magnitude fewer features not only helps interpretability, but also mitigates overfitting for the downstream algorithm, which is prominent in deep learning models in the biochemical field due to the curse of dimensionality that arises from data that are naturally high-dimensional. We propose that the generalizability of the features captured by persistence fingerprint means that persistence fingerprint can serve as a feature set for future machine learning models for molecular interaction.

Due to the good generalizability of persistence fingerprints and the high performance of PATH^−^ in discerning decoy compounds (Figure 3), we believe that the features in IPCs and persistence fingerprints (Figure 2) indirectly capture protein-ligand interactions. For example, a distance of 9.5 Å between a C*α* in the protein backbone and a carbon atom in the ligand, as captured by persistence fingerprint components, is likely to correlate with interactions between its side chain and the ligand. Other possibilities include allosteric interactions [38], domain reorientation [88], and solvent mediated interactions [108]. Remarkably, through a subsequent literature review, we discovered the features in persistence fingerprint, which were completely automatically derived, are similar to the “interaction fingerprints” manually constructed in previous works on binding affinity prediction [104, 111]. Interpretability of PATH^+^ provides verification of the robustness beyond simply benchmarking on datasets and provides insights on the geometric features important to predicting binding affinity with persistent homology. The ability to pinpoint the precise atoms that contribute to a feature in persistence fingerprint also enables us to visualize the structural changes that drive a change in binding affinity, and justify a highly efficient approximation of persistence fingerprint (Theorem 1). This foregrounds the benefits of an interpretable algorithm. PATH can be further accelerated by using approximation algorithms for persistent homology [97, 18, 21].

## 6 Conclusion

Describing the shapes of biomolecules via topological invariants is a promising direction. We filled an important gap in the field of computational structural biology by designing an interpretable vector space with only 10 dimensions for describing protein-ligand interactions, which we call persistence fingerprint. This reduces the dimensionality of the machine learning problem by over three orders of magnitude, opening the door to efficient interpretable algorithms. We showed that the discriminating power of persistence fingerprint generalizes beyond the training dataset. We provided an interpretable algorithm (PATH^+^) effective at predicting protein-ligand binding using persistence fingerprints. To our knowledge, PATH^+^ is the first interpretable algorithm for binding affinity prediction using persistent homology, while previous algorithms all resorted to black box models for regression. Despite using three orders of magnitude fewer features, PATH performed with comparable accuracy (only an approximately 7% larger RMSE on the PDBBind v2020 refined set) to TNet-BP, a previous state-of-the-art algorithm that uses persistent homology information for protein-ligand binding affinity prediction. Because the persistence fingerprint we constructed has very few dimensions, we could visually demonstrate that the features measured by our model correspond to biochemically relevant features in the HIV-1 protease-darunavir complex. We also provide the algorithm PATH^−^, which uses internuclear persistence contours to effectively discriminate between binders and non-binders. We believe PATH will improve existing structure-based drug design pipelines, provide insights in future ones, and enable a novel representation of protein-ligand interactions for future algorithms.

## Acknowledgments

We thank Graham Holt for his suggestions on initial results, and curation of protein-ligand structures. We thank Henry Childs, Kiran Kanekal, Tomás Lozano-Pérez, Carlo Tomasi and Ron Parr for detailed feedback. We thank Cynthia Rudin and Eric Chen, and all members of the Donald research group for proofreading drafts.

## Optional Appendix

The following is an optional appendix providing additional information to substantiate the claims made in the main paper. It is provided for the interested referee.

In this appendix, all distances are expressed in Angstroms; hence, the unit is omitted in the text. All logarithms are natural logarithms. To avoid confusion with vector norms, which are denoted by || · ||, the cardinality of a set *X* is denoted by 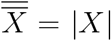. The figures in this appendix are numbered differently from the figures in the main paper. Intermediate lemmas toward Theorem 1 and their proofs are nested within the proof of this theorem. For the sake of clarity, proofs of lemmas are ended with a □ symbol, while proofs of theorems are ended with a ◼ symbol.

Appendix Section A details the precise definition of persistent homology and the construction of IPCs, which are the inputs to the regression trees of PATH^+^ and PATH^−^. In particular, Appendix Section A.1 formally defines the construction of persistent homology and persistence diagrams; Appendix Section A.2 formally defines the construction of persistence images, which is similar to IPCs; And finally, Appendix Section A.3 formally defines the construction of IPCs.

Appendix Section B details the process in which we curated the features used to construct persistence fingerprint from persistence images and justifies our choice of hyperparameters, such as the number of features and the number of regression trees. In particular, Appendix Section B.1 describes the process of selecting important features measured by their mean decrease in purity; Appendix Section B.2 describes selecting the most important features among the features resulting from Appendix Section B.1 using an *ablate-and-test* procedure; Appendix Section B.3 shows the intermediate results of the previous two subsections and supports our choice of 10 persistence fingerprint components and 13 regression trees in PATH^+^; Finally, Appendix Section B.4 confirms that decision trees are still the optimal choice of model, given these 10 selected features as input. Finding that the top 10 important features all belong to persistence images from the persistence of 0D and 1D homology groups constructed using the opposition distance (*d*_*op*_), we interpreted their geometric relevance and developed a one-dimensional analog of persistence images, which we called internuclear persistence contours (IPCs) (Main MS Section 3.1). Such a one-dimensional representation is only possible with the persistent homology of 0D and 1D Vietoris-Rips filtration constructed with the opposition distance, where all of the persistence fingerprint components come from. Appendix Section B.5 shows the generalizability of persistence fingerprints using boxplots and PaCMAP [109].

Appendix Section C explains the high complexity that is obtained by naively computing IPCs and the high complexity of previous binding affinity prediction algorithms based on persistent homology (Appendix Section C.1) and elucidates a fast and provably *ε*-accurate approximation for persistence fingerprint, which would naively have inherited the computational complexity of IPCs. Derivation of the asymptotic complexity of this approximation and proof of its accuracy are done in Appendix Section C.2 and Appendix Section C.3. Experimental runtime and accuracy for this approximation is reported in Appendix Section C.4 and validate the complexity claims and proofs in Appendix Sections C.2 and C.3.

Appendix Section D shows the decision trees of PATH^+^. In particular, Appendix Section D.1 explains how to interpret each decision tree; Appendix Section D.2 shows the full set of decision trees.

Appendix Section E shows the results of benchmarking PATH^+^ and PATH^−^ against previous binding affinity prediction algorithms in numeral tabular form, to complement the plots made in main MS Figure 3.

### A Definition of Persistent Homology

#### A.1 Construction of Persistent Homology

We describe persistent homology using notation from [30, 31, 35, 41]. Chapter 50 of [28] gives an in-depth summary of computational topology on protein structure. Visualizations of growing filtration parameters in persistent homology are made using a Mathematica notebook by Adams and Sergert [2]. Computation of persistent homology and persistence images are done using GUDHI [70], giotto-tda [101], giotto-ph [87], and Ripser [9].

##### Simplicial Homology

A *simplex* is a generalization of a triangle to arbitrary dimensions. An *n*-simplex is the convex hull of *n* + 1 points and is denoted by a square bracket around its vertices. As an example of this notation, [*v*_0_, *v*_1_, *v*_2_, …, *v*_*n*_] denotes an *n*-simplex made up of the vertices {*v*_0_, *v*_1_, …, *v*_*n*_}. Note that the order of the vertices in a simplex is important. The *face* of a simplex is a simplex with vertices any nonempty subset of the *v*_*i*_’s. For example, the two-dimensional faces of a 3-simplex (a solid tetrahedron) are four 2-simplices (filled triangles), and the one-dimensional faces of a 2-simplex (a filled triangle) are three 1-simplices (line segments).

A finite *simplicial complex K* is a finite collection of simplices such that for any three simplices *σ, τ, σ*_0_

1. If *σ* ∈ *K* and *τ* ≤ *σ*, then *τ* ∈ *K*.
2. If *σ, σ*_0_ ∈ *K*, then *σ* ∩ *σ*_0_ is either empty or a face of both.

In this manuscript, we assume finite simplicial complexes when referring to simplicial complexes. Given a simplicial complex *K*, we can define *C*_*n*_(*K*) as the vector space generated by the *n*-simplices in *K* as basis over a ring. Herein, the ring is chosen to be ℤ. Define an *n-chain* to be the formal sum 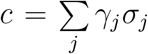, where *γ*_*j* ∈_ ℤ are the ring elements and *σ*_*j*_ are the *n*-simplices in *K*. Then *C*_*n*_(*K*) consists of all the *n*-chains.

We define the *boundary operator* ∂_*n*_ on a *n*-simplex to be an alternating sum of its (*n*− 1) dimensional faces. Formally, 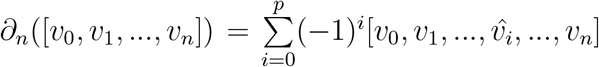 where the hat in 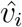 means that 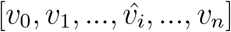 is the simplex formed by removing the *i*-th vertex in [*v*_0_, *v*_1_, …, *v*_*n*_]. For example, ∂_1_([*v*_0_, *v*_1_]) = [*v*_1_] − [*v*_0_] and ∂_2_([*v*_0_, *v*_1_, *v*_2_]) = [*v*_1_, *v*_2_] − [*v*_0_, *v*_2_] + [*v*_0_, *v*_1_]. The boundary of an *n*-chain is obtained by extending ∂_*n*_ linearly, 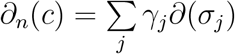 where *σ*_*j*_ are the *n*-simplices in *c*. Then ∂_*n*_ defined on the *n*-chains is a homomorphism: *C*_*n*_(*K*) → *C*_*n*−1_(*K*). This gives us a sequence of chains connected by boundary homomorphisms, called *chain complexes*:

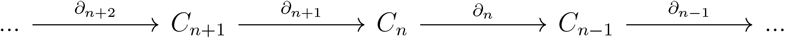

Note that ∂_*n*−1_ ○ ∂_*n*_(*c*) = 0 for any integer *n* and any *c* ∈ *C*_*n*_(*K*). The *n*-chains that have boundary 0 (i.e., in the kernel of ∂_*n*_) are called *n-cycles* and denoted *Z*_*n*_(*K*). The *n*-chains that are the boundary of (*n* + 1)-chains (i.e., in the image of ∂_*n*+1_) are called *n-boundaries* and denoted *B*_*n*_(*K*). The *n*^*th*^ *homology group H*_*k*_(*K*) is the quotient group *Z*_*n*_(*K*)*/B*_*n*_(*K*) = ker(∂_*n*_)*/*im(∂_*n*+1_). The fact that ∂_*n*_ ○ ∂_*n*+1_(*c*) = 0 tells us that *B*_*n*_(*K*) ⊂ *Z*_*n*_(*K*). Note that *B*_*n*_(*K*) is a normal subgroup because *Z*_*n*_(*K*) is abelian, so *H*_*n*_(*K*) is well-defined. By the structure theorem for finitely generated abelian groups, *H*_*n*_(*K*) ≅ ℤ^*r*^ ⊕ (ℤ */d*_1_ ℤ) ⊕ (ℤ */d*_2_ ℤ) ⊕ …⊕ (ℤ */d*_*m*_ ℤ), where *d*_1_|*d*_2_| …|*d*_*m*_ and the symbol ≅ denotes group isomorphism. Respectively, ℤ ^*r*^ and 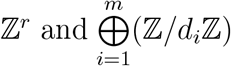 are called the *free part* and the *torsion part* of this homology group [28]. Geometric objects that are embeddable in ℝ ^3^ such as the simplicial complex for a protein-ligand complex will generally have zero torsion [28].

The *n*^*th*^ *Betti number* of *H*_*n*_(*K*), denoted *β*_*n*_, is equal to *r*, the rank of the free part of *H*_*n*_(*K*). Geometrically, the *n*^th^ Betti number can be understood as the number of *n*-dimensional holes in the complex [35]. These terms will be used interchangebly for this manuscript.

**Persistence**. Given a simplicial complex *K*, a *filtration* of *K* is a sequence of subcomplexes such that

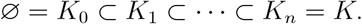

Given a set *X* with a distance function *d* : *X × X* → ℝ ∪{+∞, −∞}, we can construct the *Vietoris-Rips complex* ^3^ *of X with parameter r* ∈ [0, ∞), denoted **VR**_*r*_(*X*), by forming a simplex for every finite set of points in which every pair of points has distance at most *r*. The Vietoris-Rips filtration of *X* is then defined as {**VR**_*r*_(*X*)}_*r*∈ℝ_. Note that for any *r* ≤ *s*, we have **VR**_*r*_(*X*) ⊂ **VR**_*s*_(*X*). Therefore, the Vietoris-Rips filtration is a filtration of **VR**_∞_(*X*).

The Vietoris-Rips filtration is indexed by R, an uncountably infinite set. However, if *X* is finite, then **VR**_*r*_(*X*) can be represented by a filtration with a finite number of subcomplexes without losing information [9]. Intuitively, the information contained in {**VR**_*r*_(*X*)}_*r*∈R_ can be entirely captured by subcomplexes before and after the formation of each simplex, and there is a finite number of simplices in **VR**_∞_(*X*) for finite *X*.

While the Vietoris-Rips filtration is usually defined for metric functions, it can be easily seen that the Vietoris-Rips filtration is well-defined for the opposition distance (Eq. 2 in the Main MS) as well.

A *homology class* is an element of a homology group, termed a “class” because the group is expressed as a quotient. A homology class *α* is defined to be *born* at *K*_*i*_ if it is not in the image of the map induced by the inclusion *K*_*i*−1_ *↪K*_*i*_. Furthermore, if *α* is born at *K*_*i*_, then it *dies entering K*_*j*_ if the image of the map induced by *K*_*i*−1_ ↪ *K*_*j*−1_ does not contain the image of *α* but the image of the map induced by *K*_*i*−1_ ↪ *K*_*j*_ does.

We can precisely represent the birth and death of homology classes of a certain dimension using a *persistence diagram*, where each point (*x, y*) corresponds to an *n*-dimensional homology class (an *n*-dimensional hole) that appears (is born) at scale *x* and disappears (dies) at scale *y*. We denote the *n*^th^ persistence diagram as *D*_*n*_(*X*) and define 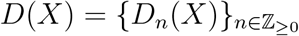 Persistence diagrams has been shown to be stable with respect to small perturbations in input:

###### Theorem 2

**(Stability of persistence diagrams [15, 16])**. *For finite metric spaces* (*X, d*_*X*_) *and* (*Y, d*_*Y*_), *we have d*_*b*_(*D*(*X*), *D*(*Y*)) ≤ 2*d*_*GH*_(*X, Y*). *Here d*_*b*_ *and d*_*GH*_ *are the bottle-neck distance and the Gromov-Hausdorff distance respectively between persistence diagrams and metric spaces [15, 16]*.

Proof of Theorem 2 can be found in [15, 16].

Thus, in embedding biochemical structure, small differences in the protein structure which arise often due to the structural heterogeneity of proteins [25] will have little effect on the resulting representation.

#### A.2 Persistence Images

The concept of *persistence image* was developed by Adams et al. [1] as a stable vectorization method for persistence diagrams. Let *T* : ℝ^2^ → ℝ^2^ be the linear transformation *T* (*x, y*) = (*x, y* − *x*). Let *ϕ*_*u*_ : ℝ^2^ → ℝ be a differentiable probability distribution with mean *u* ∈ ℝ^2^. Additionally, fix a nonnegative weighting function *f* : ℝ^2^ → ℝ that is zero along the horizontal axis, continuous, and piecewise differentiable. Then the *persistence surface* of a persistence diagram *D*_*n*_(*X*) is defined as the continuous function 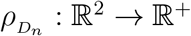 given by

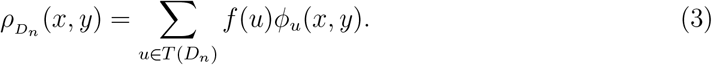

Hence, the persistence surface of a diagram is the sum of translated and weighted Gaussian kernels centered at persistence of each homology class, represented by its birth and death radii as coordinates (*x, y*) in the 2D plane. Given a positive pixel width *w* ∈ ℝ^+^, we can construct a partition of ℝ^2^ into square *pixels* with side length *w*, which we denote by 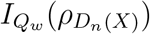 is the collection of pixels 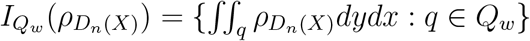.

##### Theorem 3

**(Stability of persistence images [1])**. *The persistence image with Gaussian distributions is stable with respect to the 1-Wasserstein distance between diagrams*.

Proof of Theorem 3 can be found in [1].

Persistence images are provably stable with respect to persistence diagrams, and thus stable with respect to perturbations by Theorem 2.

While the persistence image is defined over all of ℝ^2^, in this manuscript, only the subset of pixels in the range [0, 50] *×* [0, 50] (in Angstroms) are used. In this manuscript, each persistence image is created with standard deviation of Gaussian kernel *σ* = 0.1, and the width of each pixel is 0.5 Å. The persistence images are then flattened into a 10,000-dimensional vector as inputs to regression algorithms (Appendix Section B). PATH^+^ and PATH^−^ uses persistence images from dimensions *n* = {0, 1}, and TNet-BP uses persistence images from dimensions *n* = {0, 1, 2}.

#### A.3 Definition of Internuclear Persistence Contours (IPCs)

As discussed in the beginning of Section 3.1, the persistence of each 0D or 1D homology group constructed with the opposition distance can be captured with a single scalar value (death radius for 0D, birth radius for 1D). We call each of these scalar values a *critical value* [28] of the filtration. Then the 0D or 1D persistent homology information can be captured by a sequence of critical values {*r*_1_, *r*_2_, …, *r*_*n*_} without losing information. In the following subsection, we formalize this intuition.

Fix a protein-ligand complex and fix a persistence fingerprint component. Fixing the persistence fingerprint component fixes a subset of protein atoms *P* and a subset of ligand atoms *L* matching certain element types^4^. The protein atom subset *P*, ligand atom subset *L* for each persistence fingerprint component are given by Table 2 in the Main MS. For example, if we fix the 5^th^ persistence fingerprint component, we read from Table 2 the information then *P* is the subset of all protein nitrogen atoms, *L* is the subset of all ligand carbon atoms^5^. Then the protein-ligand complex, which we notate as *S* = *P* ⊔ *L*, is a disjoint union of the atoms in *P* and *L*, while each atom additionally has information of its affiliation (whether it came from the protein or the ligand). The filtration in concern in the remaining proof will be the Vietoris-Rips filtration of *S* of filtration radius *r* with *d*_*op*_, which is **VR**_*r*_(*S*).

| Protein Atom | Ligand Atom | IPC Dimension | IPC Density Bin ($\text{\AA}$) |
| --- | --- | --- | --- |
| N | C | 1 | [10.0, 10.5] |

##### Lemma 1

(Persistence of 1D holes). *A hole in the 1D Vietoris-Rips filtration with d*_*op*_ *as distance function is generated by a 1-cycle and never dies. The number of 1D holes in this construction monotonically increases with increasing filtration radius r*.

**Proof of Lemma 1:** Because *B*_2_ = im(∂_2_) is generated by the faces of 2-simplices, each element in its basis must be of the form [*v*_*y*_, *v*_*z*_] − [*v*_*x*_, *v*_*z*_] + [*v*_*x*_, *v*_*y*_] = ∂([*v*_*x*_, *v*_*y*_, *v*_*z*_]) resulting from some 2-simplex [*v*_*x*_, *v*_*y*_, *v*_*z*_]. However, for any three atoms {*x, y, z}* ∈ *S*, two of them must have the same affiliation (either *P* or *L*), and the opposition distance between these two atoms with the same affiliation (i.e., both atoms are from the protein or both atoms are from the ligand) is ∞. This means that [*y, z*] − [*x, z*] + [*x, y*] cannot be a generator for *B*_2_. Therefore, for any filtration radius *r*, we have *B*_2_ ≅ 0, where the symbol ≅ denotes group isomorphism.

Then for any positive *r*, we have *H*_1_ ≅ ker(∂_1_) = *Z*_1_, which are the 1-cycles. Obviously, no 1-cycle perishes with increasing *r*. As a result, the rank of the 1^st^ homology group (i.e., number of 1D holes) monotonically increases with increasing filtration radius *r*. □

##### Lemma 2

(Critical values of filtration with *d*_*op*_). *The persistence of each 0D and 1D hole in the Vietoris-Rips complex constructed with d*_*op*_ *can be faithfully characterized by a single scalar value. Namely, death radius for a 0D hole and birth radius for a 1D hole. Name this scalar value the* critical value *of the hole, then the set of critical values R for a filtration satisfy*

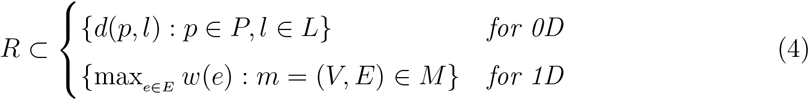

*Where M is defined as the set of all cycles m generated by a sequence of alternating protein atoms and ligand atoms, and each cycle is viewed as a graph in which each edge e has a weight w*(*e*) *equal to the Euclidean distance between the two atoms it connects*.

**Proof of Lemma 2:** An intuition of this proof can be found in Figure 7. Because the rank of the 0^th^ homology group *H*_0_ of a simplicial complex is just number of connected components in this complex, each 0D hole is born at *r* = 0. Hence the persistence of each 0D hole can be characterized only by the death radius of the hole, which is distance between a protein and a ligand atom, which is in the set of all distances between protein and ligand atoms. Thus the 0D case in Lemma 2 holds.

**Fig. 7.**
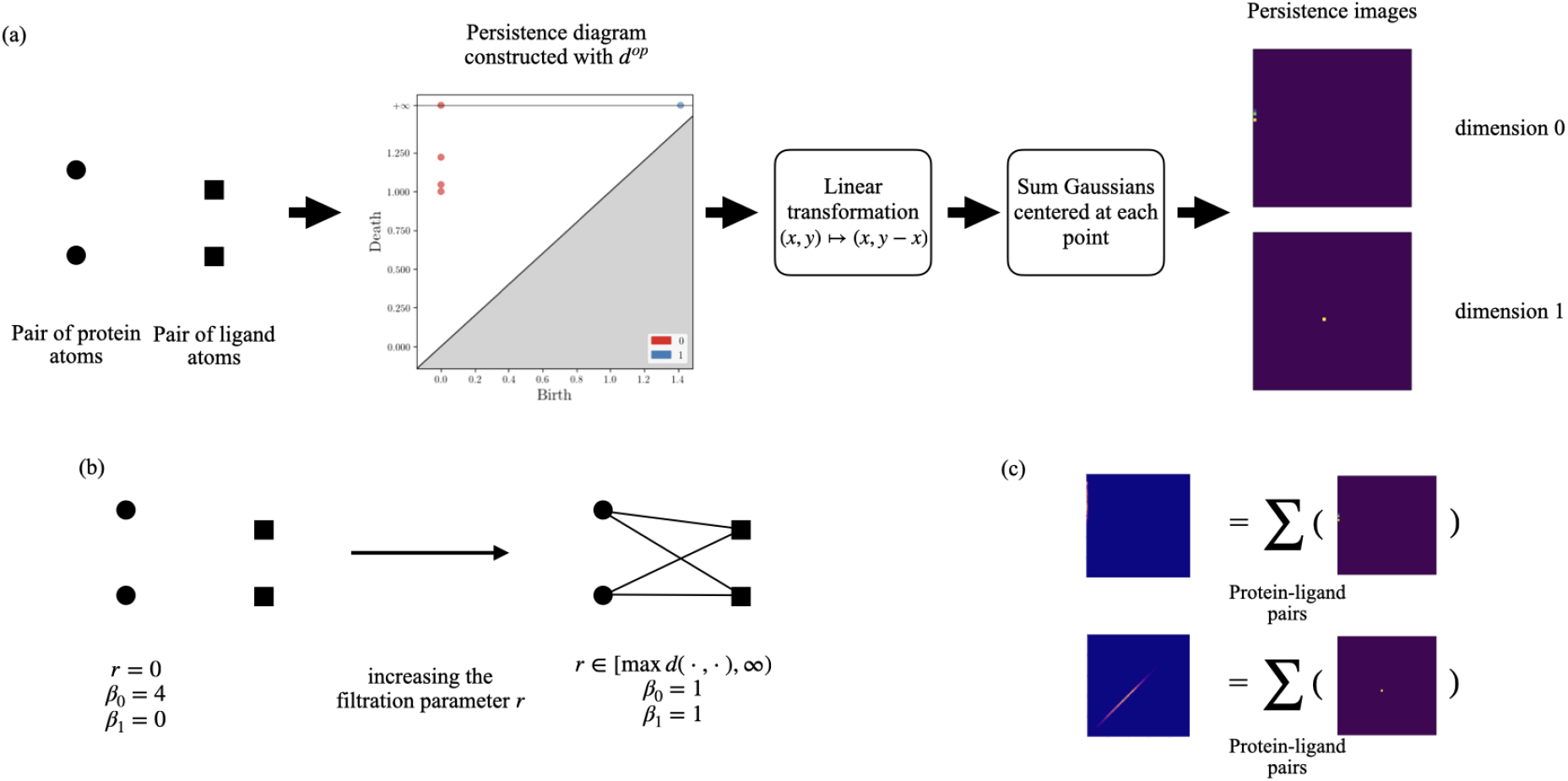
Construction of persistence images with *d*_*op*_. **(a)** In the persistence diagram, a pair of protein atom and a pair of ligand atoms will create 4 points representing 0-dimensional holes, and 1 points represent a 1-dimensional hole. This results in a few bright areas on the left edge of the persistence image for dimension 0 and a bright spot on the diagonal of the persistence image for dimension 1. **(b)** Illustration of how *d*_*op*_ creates a 1-dimensional hole in the persistence diagram representation for the two pairs of atoms: For protein atoms *p*_1_, *p*_2_ and ligand atoms *l*_1_, *l*_2_, when the filtration parameter *r* increases beyond max{*d*(*p*_*i*_, *l*_*j*_) : *i, j* ∈ {1, 2}} distance among the atoms, there is always a 1-dimensional hole (Lemma 1). Intuitively (see Lemma for proof), there never exists a filled triangle because the distance between two atoms with the same affiliation (protein or ligand) is ∞. **(c)** The persistence image representation of a protein-ligand complex is the sum of the individual interactions at different scales, taking multiplicity into consideration.

For 1D holes, the death radius is ∞ by Lemma 1, so the persistence of each 1D hole can be characterized only by the birth radius of the hole, which is the birth radius of a 1-cycle by Lemma 1. Note that because *d*_*op*_(*p*_*i*_, *p*_*j*_) = ∞ and *d*_*op*_(*l*_*i*_, *l*_*j*_) = ∞, a 1-cycle in *d*_*op*_ must consists of a sequence of alternating protein atoms and ligand atoms and looks like *Z* = [*p*_1_, *l*_1_] + [*l*_1_, *p*_2_] +[*p*_2_, *l*_3_] +[*l*_3_, *p*_4_] + … +[*l*_*n*−1_, *p*_*n*_] +[*p*_*n*_, *l*_*n*_] +[*l*_*n*_, *p*_*n*−1_] +[*p*_*n*−1_, *l*_*n*−2_] + … +[*p*_3_, *l*_2_] +[*l*_2_, *p*_1_] (Figure 9). Let the elements in this sum be labeled ζ = {(*p*_*i*_, *l*_*j*_) : [*p*_*i*_, *l*_*j*_] ∈ *Z* ∨ [*l*_*j*_, *p*_*i*_] ∈ *Z*}. Then this 1-cycle has birth time 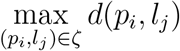

**Fig. 8.**
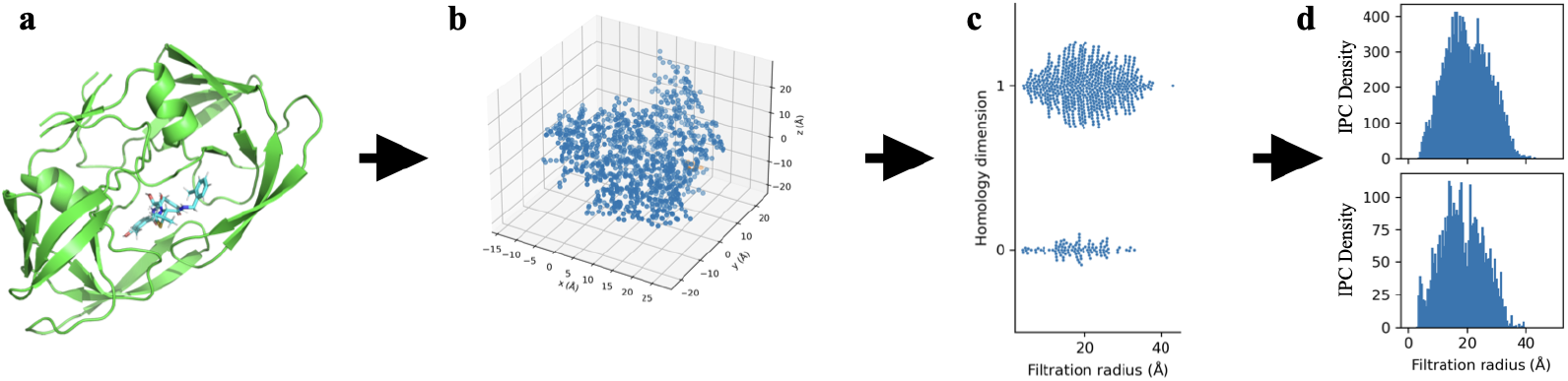
Construction of internuclear persistence contours (IPCs). IPCs are constructed for each pair of protein and ligand heavy atoms in the training dataset, and the integrals of IPCs in certain bins are selected into persistence fingerprint. ^**a**^Protein-ligand complex shown as example: the HIV protease (mutant Q7K/L33I/L63I) complexed with KNI-764 (an inhibitor), PDB ID: 1msm [102]. ^**b**^A point cloud is created from subsets of atoms with certain element types in protein and ligand (detailed in Table 4). Shown as example: carbon atoms from the protein and carbon atoms from the ligand. ^**c**^Persistent homology is calculated on this point cloud using opposition distance, and the birth filtration radii for 1D homology groups and death filtration radii for the 0D homology groups are collected (see Section 3.1 for why these suffice). ^**d**^Internuclear persistence contours (IPCs) are constructed by summing Gaussians centered at each of the birth or death radius. The IPCs in PATH are constructed with a standard deviation of 0.1. Two IPCs are shown. Top: carbon-carbon IPC dimension 1. Bottom: carbon-carbon IPC dimension 0.

**Fig. 9.**
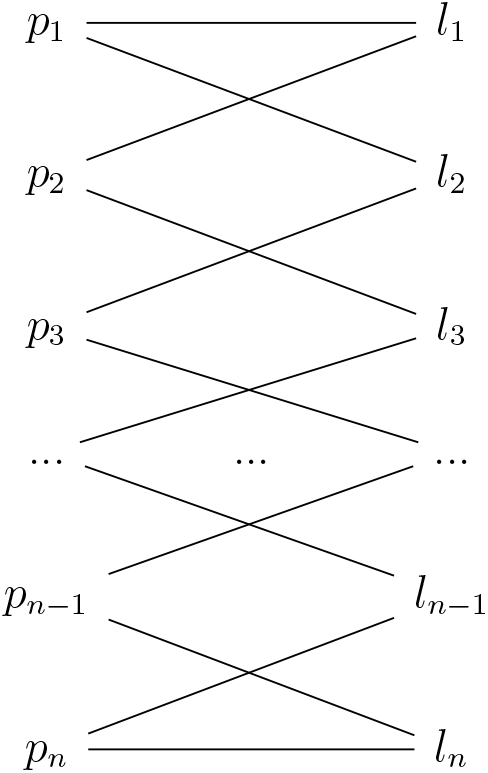
Example of a 1-cycle in the Vietoris-Rips filtration of a protein-ligand complex constructed with opposition distance. Notice that this 1-cycle is generated by a sequence of alternating protein atoms and ligand atoms.

*Z* can be viewed as a weighted graph *m* = (*V, E*) with vertices *V* = {*p*_*i*_} ∪ {*l*_*i*_}, edges *E* = ζ and let each edge be endowed the Euclidean distance between its vertices as its weight. Since the critical value of every hole in *H*_1_ is generated by such a graph, the 1D case in Lemma 2 holds. Note that *m* is similar to a bipartite matching between the protein and ligand atoms (Figure 9), hence we refer to the construction of 1D homologies as *bipartite matchings of the protein and ligand atoms*. □

With Lemma 2, we know that the persistent homology constructed on a protein-ligand complex in dimensions 0 and 1 can be faithfully represented by a sequence of critical values *R*. Thus, we can define the *Internuclear Persistence Contour (IPC)*, which is a non-negative real valued function *γ* : ℝ → [0, ∞). To construct the (continuous) IPC, we sum Gaussian kernels centered at each of the critical values (*r*_*i*_’s). Let us denote a Gaussian kernel with mean *µ* and standard deviation *σ* by *g*_*µ*_(*x*). The IPC is defined as follows:

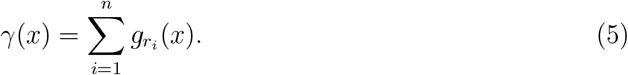

Note the similarity of *γ* with the definition of persistence image *ρ* (from Appendix Section A.2), because *γ* is a special case of *ρ* for Vietoris-Rips filtration constructed with the opposition distance *d*_*op*_ (Section 2.2) in dimensions 0 and 1, which only has one degree of freedom for the persistence of each homology group.

In our current implementation, the zero-dimensional IPCs are approximated by taking the left-most column of pixels of the 0D persistence image and the one-dimensional IPCs are approximated by taking the diagonal of pixels of the 1D persistence image. To put precisely, in PATH, 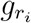 is defined as follows:

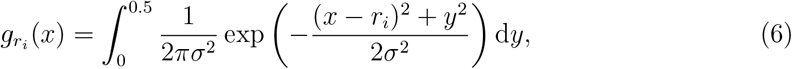

where *x* and *y* can be regarded as coordinates of the horizontal and vertical axes, respectively, of the two dimensional persistence image. This can be easily computed by existing packages for topological data analysis [26, 101], and behaves sufficiently similar to the Gaussian kernel so that the computational complexity and approximation error guarantees (Appendix Section C) still hold. In a future implementation, we plan to use the exact Gaussian kernel instead of this approximation.

Given a point cloud, we can compute the 0D and 1D persistent homologies with *d*_*op*_ and construct two IPCs from it: we call IPCs constructed using 0D death times the *0D IPC*, and IPCs constructed using 1D birth times the *1D IPC*. Definition of *IPC density* follows by integrating its underlying IPC over equally spaced bins. For this IPCs in this paper, the bins are constructed with a width of 0.5 starting from 0 (i.e., bins = {[0, 0.5]f, [0.5, 1.0], [1.0, 1.5]…}1), and the IPC densities of a certain IPC *γ* will be the ordered set 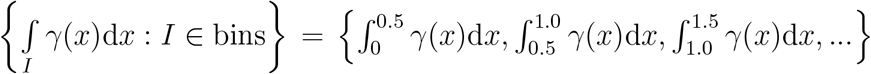 IPC density provides a fixed-size vectorization of IPCs. Stability of IPC density follows from the stability of persistence images (Theorem 3).

### B. Feature Selection

#### B.1 Feature Selection by Mean Decrease in Impurity (MDI)

**Initial images**. This section describes the process in which we selected the features used to construct persistence fingerprint from IPCs.

We created 40 subsets of atoms of from the protein and ligand (Table 4). From each subset of atoms, a persistence diagram is constructed for each of the homology dimensions {0, 1, 2}. Subsequently, three persistence images are created, one for each dimension. The resulting 120 persistence images constitute a superset of the features constructed in TNet-BP [13]. We call this set of persistence images the *initial images*.

**Table 4.** Atom subsets and distance functions used to construct persistent homology for the *initial images*. The set of protein atoms is denoted *P* and the set of ligand atoms is denoted *L*. The set of heavy atom types in the protein is defined as *P*_*e*_ = {C, N, O, S}, and the set of heavy atom types in the protein is defined as *L*_*e*_ = {C, N, O, S, P, F, Cl, Br, I}. These are exactly the atom subsets and distance functions used to construct the features of TNet-BP [13]. We construct a persistence image for each of the homology dimensions {0, 1, 2} in each of these atom subsets.

| Set | Atoms Used | Distance Function |
| --- | --- | --- |
| 0-35 | $\{a \in P : T(a) = e_P\} \cup \{a \in L : T(a) = e_L\} : e_P \in P_e, e_L \in L_e$ | $d_{op}$ |
| 36 | $\{a \in P : T(a) \in P_e\}$ | Euclidean |
| 37 | $\{a \in P : T(a) \in P_e\} \cup \{a \in L : T(a) \in L_e\}$ | Euclidean |
| 38 | $\{a \in P : T(a) = C\}$ | Euclidean |
| 39 | $\{a \in P : T(a) = C\} \cup \{a \in L : T(a) = C\}$ | Euclidean |

**Table 5.** Performance of intermediate GBR models across 100 random restarts (except TNet-BP as reported in [13]) on PDBBind v2020 refined set shows that intermediate ablated models did not lose much performance. The values are reported in *pK*_*i*_ = − log_10_ *K*_*i*_ or *pK*_*d*_ = − log_10_ *K*_*d*_, as is done in PDBBind [107]. All values measured by us are reported in mean *±* standard deviation across 100 random restarts. Train:Test ratio is 9-to-1.

| Model architecture | TNet-BP (as reported in [13]) | TNet-BP (our implementation) | GBR | GBR | GBR |
| --- | --- | --- | --- | --- | --- |
| Number of features | 14472 | 14472 | 1.2M (flattened initial images) | 77 (after ablation by MDI) | 10 (after ablate-and-test i.e., persistence fingerprint) |
| RMSE | 1.37 | $1.69 \pm 0.08$ | $1.36 \pm 0.04$ | $1.44 \pm 0.04$ | $1.45 \pm 0.04$ |
| Pearson | 0.83 | $0.56 \pm 0.04$ | $0.72 \pm 0.02$ | $0.67 \pm 0.02$ | $0.67 \pm 0.02$ |

##### Finding the important features for prediction

We trained gradient boosting regressors (GBRs) [34] using the set of flattened initial images of each complex the PDBBind v2020 refined set as inputs and their binding affinities as labels. Then we measured each feature’s importance using the data points’ mean decrease in impurity (MDI) at that feature, where the impurity is measured by Gini index [67].

We identified the features that were present at least once among the top 10 most important features across 100 random splits of the data. We found 77 features that satisfy this criterion. We also find that performance of a GBR trained on these 77 features is comparable to the performance of a GBR trained on the original 1.2 million features of the flattened initial images (Table 5 in appendix). Because most of these features will be pruned regardless during the next step (iterative ablation), we decided that it is sufficient to proceed into the next step using only these features.

#### B.2 Iterative ablation

Starting with these 77 features, we performed an iterative procedure to identify and ablate the least important features: At each iteration, we ablated the feature that when deleted, GBRs trained on the remaining features had the lowest root mean squared error (RMSE) across 50 random data splits. We term this procedure the *ablate-and-test* procedure. Code for ablate-and-test follows this paragraph, and the accuracies of optimal GBRs trained on each number of features from 77 to 1 are shown in Appendix Section B.3. The resulting set of 10 features is termed the *persistence fingerprint*.

##### Algorithm for Iteratively Ablating Features

Following is the algorithm that takes in the 77-dimensional vector of features (`highly_selected_observations`) resulted from Appendix Section B.1, and iteratively deletes the feature that when deleted, results in models with the lowest RMSE across 10 runs.

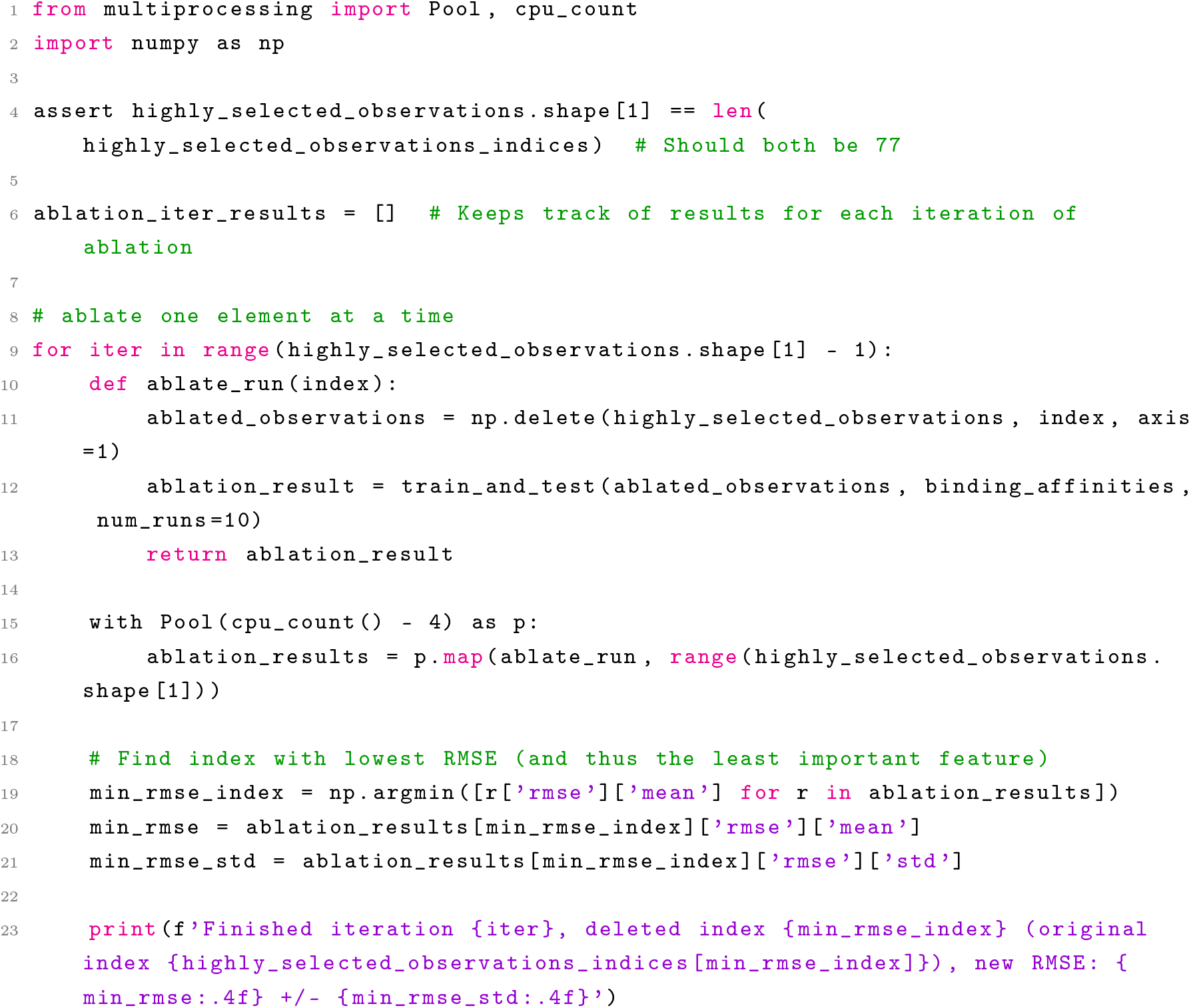
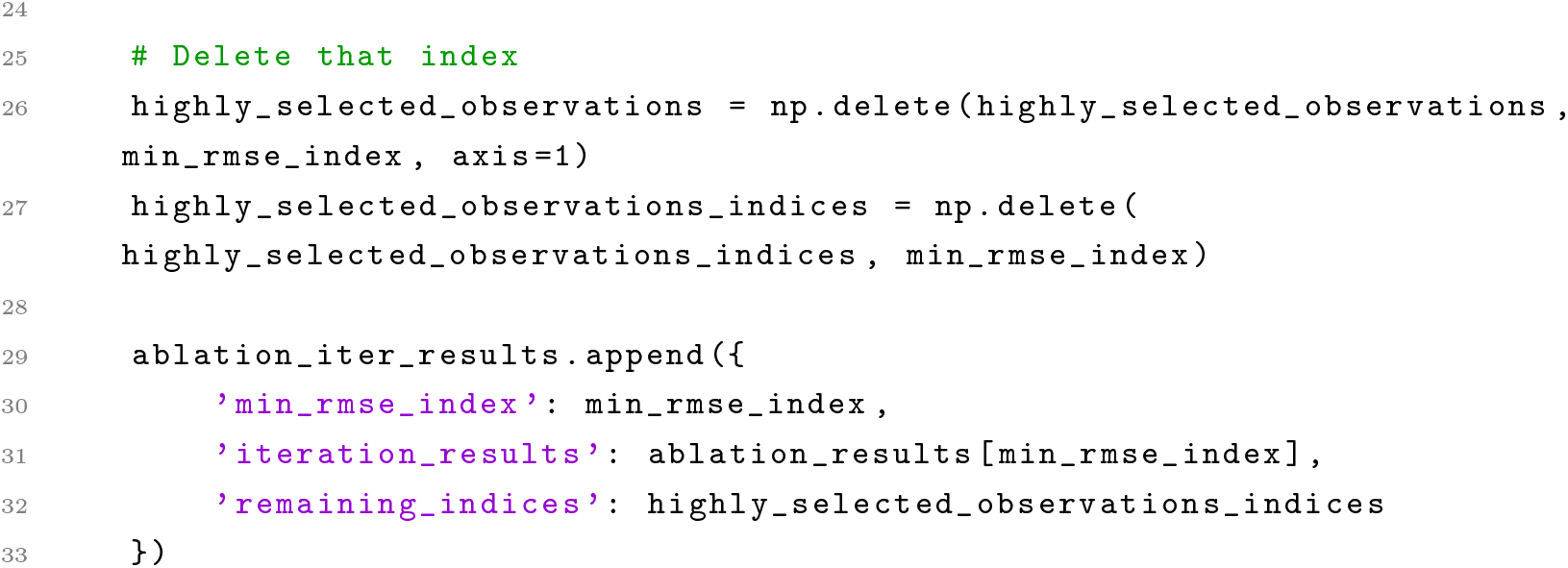

#### B.3 Final and Intermediate Results from Iterative Ablation

See Table 5 for performances of gradient boosting regressors with 1.2M, 77 and 10 features respectively, all with 100 trees. See Figures 10 and 11 for the accuracy of optimal GBRs with respect to the number of input features and number of trees, respectively. After balancing interpretability with performance, we selected the vector with 10 features to be the *persistence fingerprint* and a regressor with 13 regression trees.

**Fig. 10.**
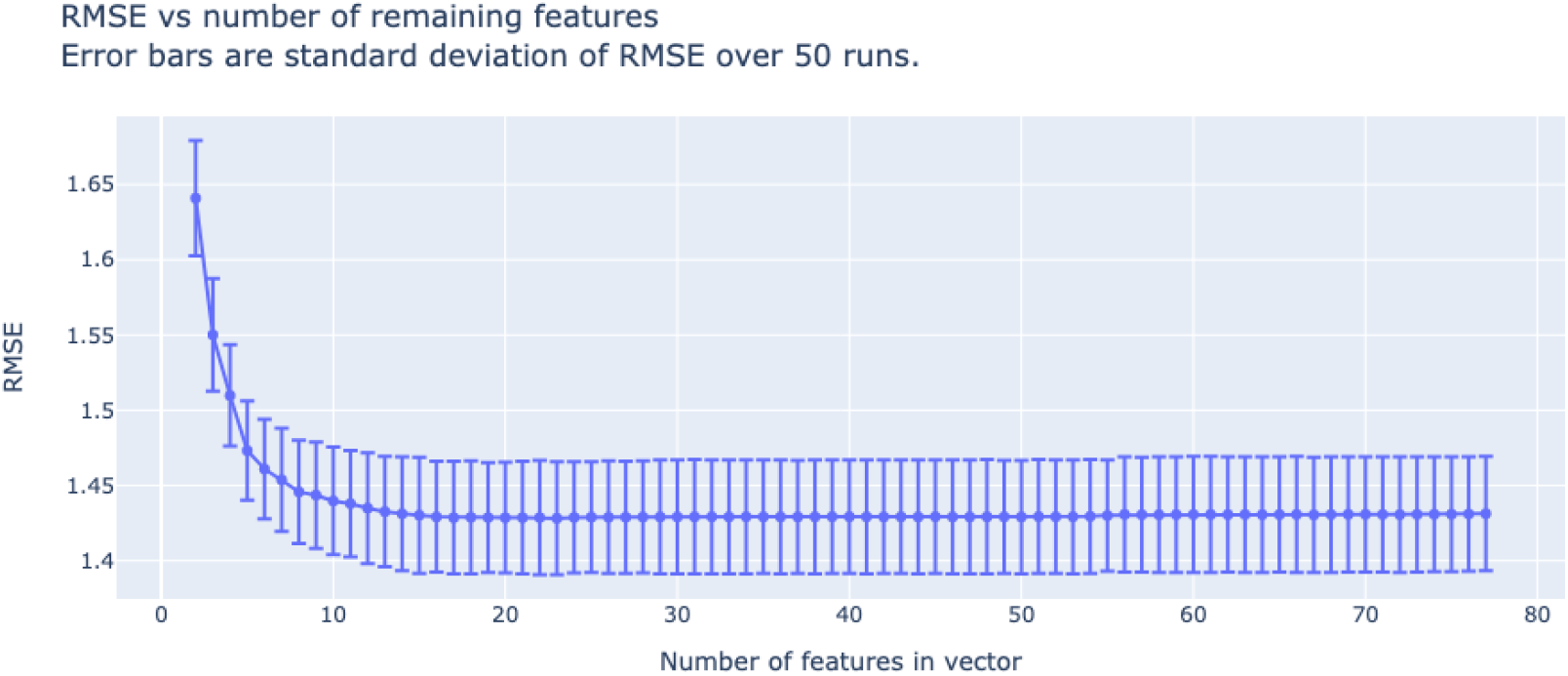
RMSE of trained GBRs with respect to number of features (from 77 to 1) in the vector through the iterative refinement process. We have chosen the number of features to be 10, as we observed that 10 is the number at which the performance of trained GBRs start to degrade quickly. As a result, this vector with 10 features (components) is termed *persistence fingerprint*. In this plot, the tree depth is set to be 3, learning rate is set to be 0.3, and number of estimators (trees) is set to be 100.

**Fig. 11.**
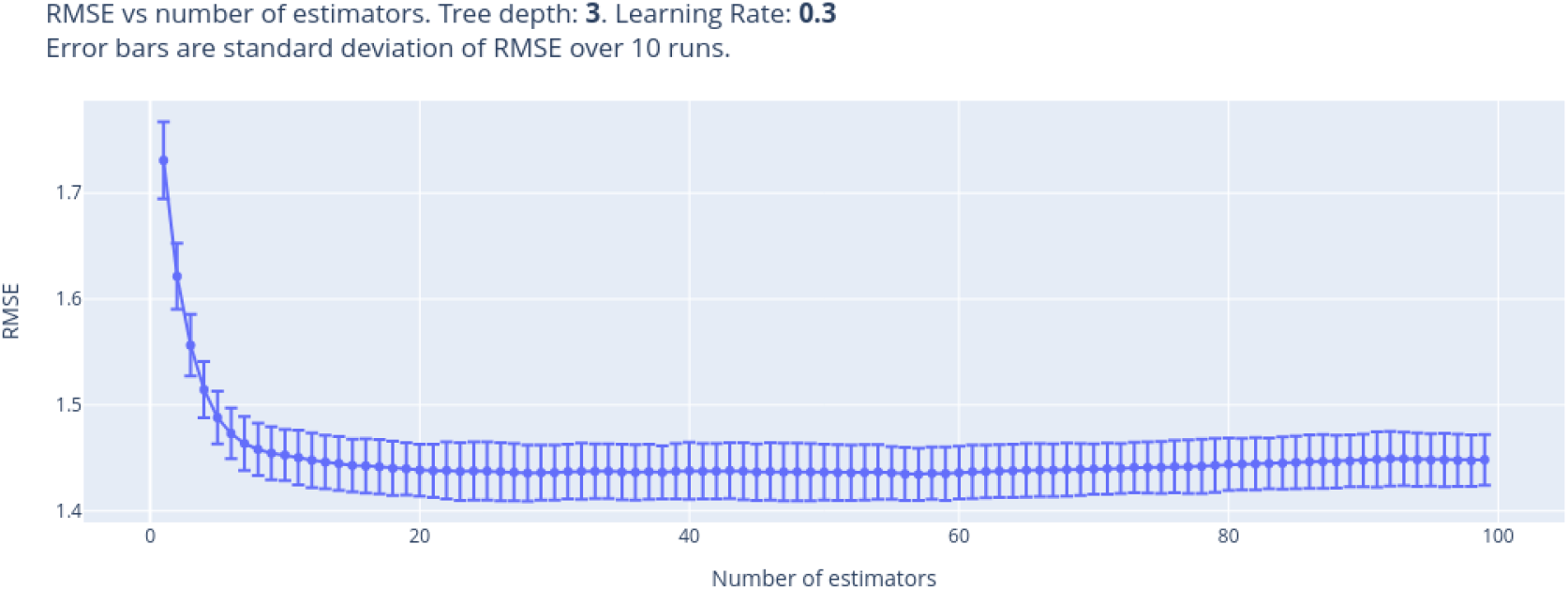
RMSE of trained GBRs with respect to the number of trees, using persistence fingerprint as input, with tree depth 3 and learning rate 0.3. These tree depth and learning rate were determined using a hyperparameter search for interpretability and performance. We selected 13 as the number of trees after balancing interpretability and performance.

#### B.4 Results for Alternative Regression Methods on the 10 Selected Features

To confirm that decision trees are the optimal model choice on the final curated features, we used LazyPredict [85] and trained the selected 10 features (i.e., persistence fingerprint) on 41 other regressors. We found that gradient boosting regressor still performs reasonably well among other regressors. The results are shown in Table 6.

**Table 6.** Results for training and testing the selected 10 features (i.e. persistence fingerprint) on other regression methods across 100 random restarts shows that regression trees are still the most performant models on the persistence fingerprint representation. Each entry of the table shows mean *±* standard deviation. All results are reported in *pK*_*i*_ = − log_10_ *K*_*i*_ or *pK*_*d*_ = − log_10_ *K*_*d*_. As in Table 5, the dataset is PDBBind v2020 refined set. Train:Test ratio is 9-to-1. All models are run in their default parameters in the LazyPredict [85] package.

| | RMSE across 100 restarts | $R^2$ across 100 restarts | Adjusted $R^2$ across 100 restarts |
| --- | --- | --- | --- |
| ExtraTreesRegressor | $1.38 \pm 0.04$ | $0.49 \pm 0.03$ | $0.48 \pm 0.03$ |
| RandomForestRegressor | $1.40 \pm 0.04$ | $0.47 \pm 0.04$ | $0.46 \pm 0.04$ |
| LGBMRegressor | $1.42 \pm 0.04$ | $0.46 \pm 0.03$ | $0.45 \pm 0.04$ |
| HistGradientBoostingRegressor | $1.43 \pm 0.05$ | $0.46 \pm 0.03$ | $0.45 \pm 0.04$ |
| NuSVR | $1.43 \pm 0.05$ | $0.45 \pm 0.03$ | $0.44 \pm 0.04$ |
| SVR | $1.43 \pm 0.05$ | $0.45 \pm 0.04$ | $0.44 \pm 0.04$ |
| MLPRegressor | $1.44 \pm 0.05$ | $0.45 \pm 0.04$ | $0.43 \pm 0.04$ |
| GradientBoostingRegressor | $1.44 \pm 0.04$ | $0.44 \pm 0.03$ | $0.43 \pm 0.03$ |
| KNeighborsRegressor | $1.47 \pm 0.05$ | $0.42 \pm 0.04$ | $0.41 \pm 0.04$ |
| BaggingRegressor | $1.47 \pm 0.05$ | $0.42 \pm 0.04$ | $0.41 \pm 0.04$ |
| XGBRegressor | $1.48 \pm 0.05$ | $0.41 \pm 0.04$ | $0.40 \pm 0.04$ |
| AdaBoostRegressor | $1.51 \pm 0.04$ | $0.39 \pm 0.03$ | $0.38 \pm 0.03$ |
| BayesianRidge | $1.56 \pm 0.05$ | $0.35 \pm 0.03$ | $0.34 \pm 0.04$ |
| ElasticNetCV | $1.56 \pm 0.05$ | $0.35 \pm 0.03$ | $0.34 \pm 0.03$ |
| RidgeCV | $1.56 \pm 0.05$ | $0.35 \pm 0.03$ | $0.34 \pm 0.04$ |
| LassoCV | $1.56 \pm 0.05$ | $0.35 \pm 0.03$ | $0.34 \pm 0.04$ |
| Ridge | $1.56 \pm 0.05$ | $0.35 \pm 0.03$ | $0.34 \pm 0.04$ |
| TransformedTargetRegressor | $1.56 \pm 0.05$ | $0.35 \pm 0.03$ | $0.34 \pm 0.04$ |
| LinearRegression | $1.56 \pm 0.05$ | $0.35 \pm 0.03$ | $0.34 \pm 0.04$ |
| LassoLarsCV | $1.56 \pm 0.05$ | $0.35 \pm 0.03$ | $0.34 \pm 0.04$ |
| LassoLarsIC | $1.56 \pm 0.05$ | $0.35 \pm 0.03$ | $0.34 \pm 0.04$ |
| HuberRegressor | $1.56 \pm 0.05$ | $0.35 \pm 0.04$ | $0.34 \pm 0.04$ |
| SGDRegressor | $1.56 \pm 0.05$ | $0.35 \pm 0.03$ | $0.34 \pm 0.04$ |
| LinearSVR | $1.56 \pm 0.05$ | $0.35 \pm 0.04$ | $0.34 \pm 0.04$ |
| OrthogonalMatchingPursuitCV | $1.56 \pm 0.05$ | $0.35 \pm 0.03$ | $0.33 \pm 0.04$ |
| TweedieRegressor | $1.58 \pm 0.05$ | $0.34 \pm 0.03$ | $0.32 \pm 0.03$ |
| PoissonRegressor | $1.58 \pm 0.05$ | $0.33 \pm 0.04$ | $0.32 \pm 0.04$ |
| GammaRegressor | $1.59 \pm 0.05$ | $0.32 \pm 0.03$ | $0.31 \pm 0.03$ |
| LarsCV | $1.65 \pm 0.23$ | $0.26 \pm 0.29$ | $0.25 \pm 0.29$ |
| OrthogonalMatchingPursuit | $1.65 \pm 0.05$ | $0.27 \pm 0.04$ | $0.25 \pm 0.04$ |
| ElasticNet | $1.73 \pm 0.05$ | $0.20 \pm 0.02$ | $0.18 \pm 0.02$ |
| Lasso | $1.93 \pm 0.05$ | $0.00 \pm 0.00$ | $-0.01 \pm 0.01$ |
| LassoLars | $1.93 \pm 0.05$ | $0.00 \pm 0.00$ | $-0.01 \pm 0.01$ |
| DummyRegressor | $1.94 \pm 0.05$ | $-0.00 \pm 0.00$ | $-0.02 \pm 0.00$ |
| DecisionTreeRegressor | $1.99 \pm 0.06$ | $-0.06 \pm 0.08$ | $-0.08 \pm 0.08$ |
| ExtraTreeRegressor | $2.01 \pm 0.06$ | $-0.08 \pm 0.09$ | $-0.10 \pm 0.09$ |
| RANSACRegressor | $2.08 \pm 0.21$ | $-0.17 \pm 0.25$ | $-0.19 \pm 0.25$ |
| Lars | $2.55 \pm 0.76$ | $-0.90 \pm 1.11$ | $-0.93 \pm 1.13$ |
| PassiveAggressiveRegressor | $2.59 \pm 0.60$ | $-0.87 \pm 0.92$ | $-0.91 \pm 0.94$ |
| GaussianProcessRegressor | $3.63 \pm 0.39$ | $-2.57 \pm 0.84$ | $-2.64 \pm 0.85$ |
| KernelRidge | $6.60 \pm 0.07$ | $-10.64 \pm 0.64$ | $-10.87 \pm 0.65$ |

#### B.5 Representability of persistence fingerprint

To supplement Figure 14 in the Main MS, which shows the representability of persistence fingerprints with PaCMAP [109], Figure 12 shows boxplots where each boxplot represents a different component of the persistence fingerprint, and each data point represents a protein-ligand complex in the PDBBind v2020 refined set. This figure shows the distribution of magnitude of each component of persistence fingerprint for protein-ligand complexes, grouped in quartiles by their experimentally-measured binding affinities. Figure 13 show boxplots for the Binding MOAD and BindingDB datasets produced using the same methodology.

**Fig. 12.**
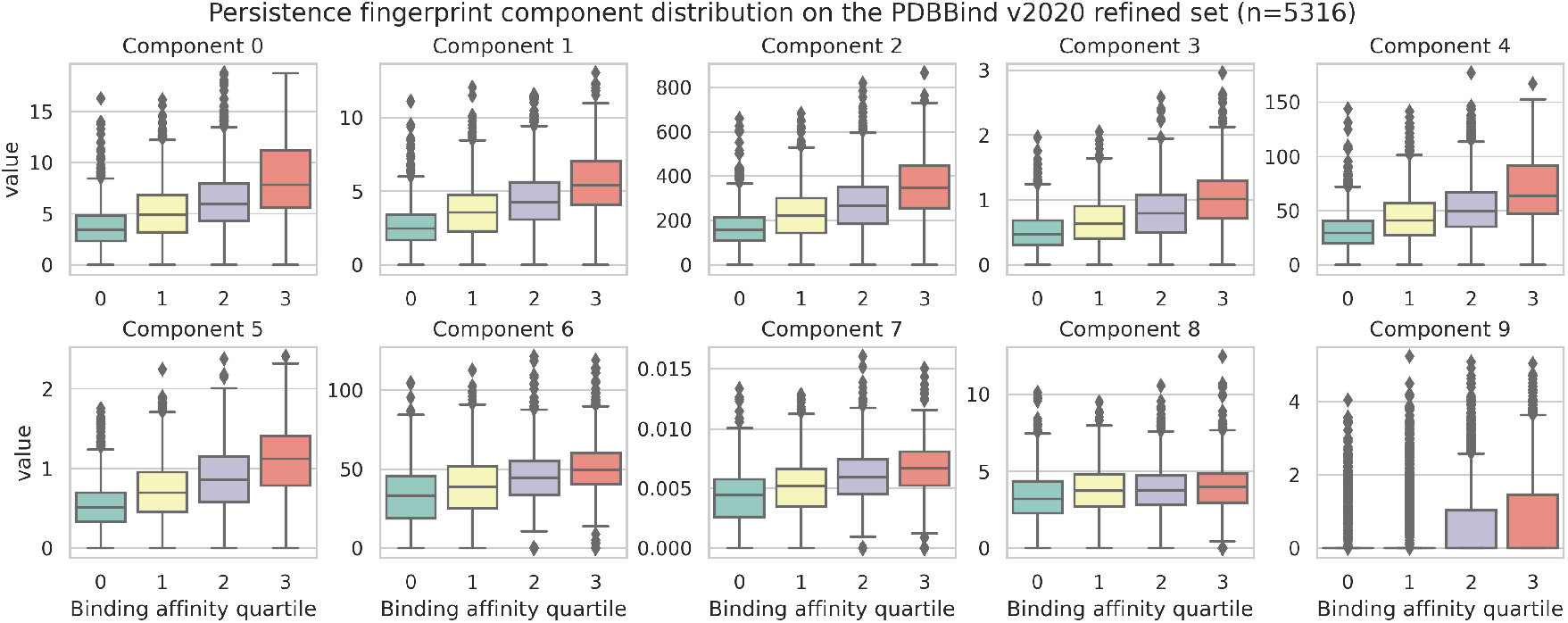
Persistence fingerprint has discriminating power on binding affinity. Boxplots of each persistence fingerprint component with respect to different quartiles of binding affinity on the PDBBind v2020 refined set show that binding affinity correlates with the values of persistence fingerprint components. The *x*-axis and color correspond to quartiles of data by their experimental binding affinity. The *y*-axis corresponds to the values of at that certain persistence fingerprint component.

**Fig. 13.**
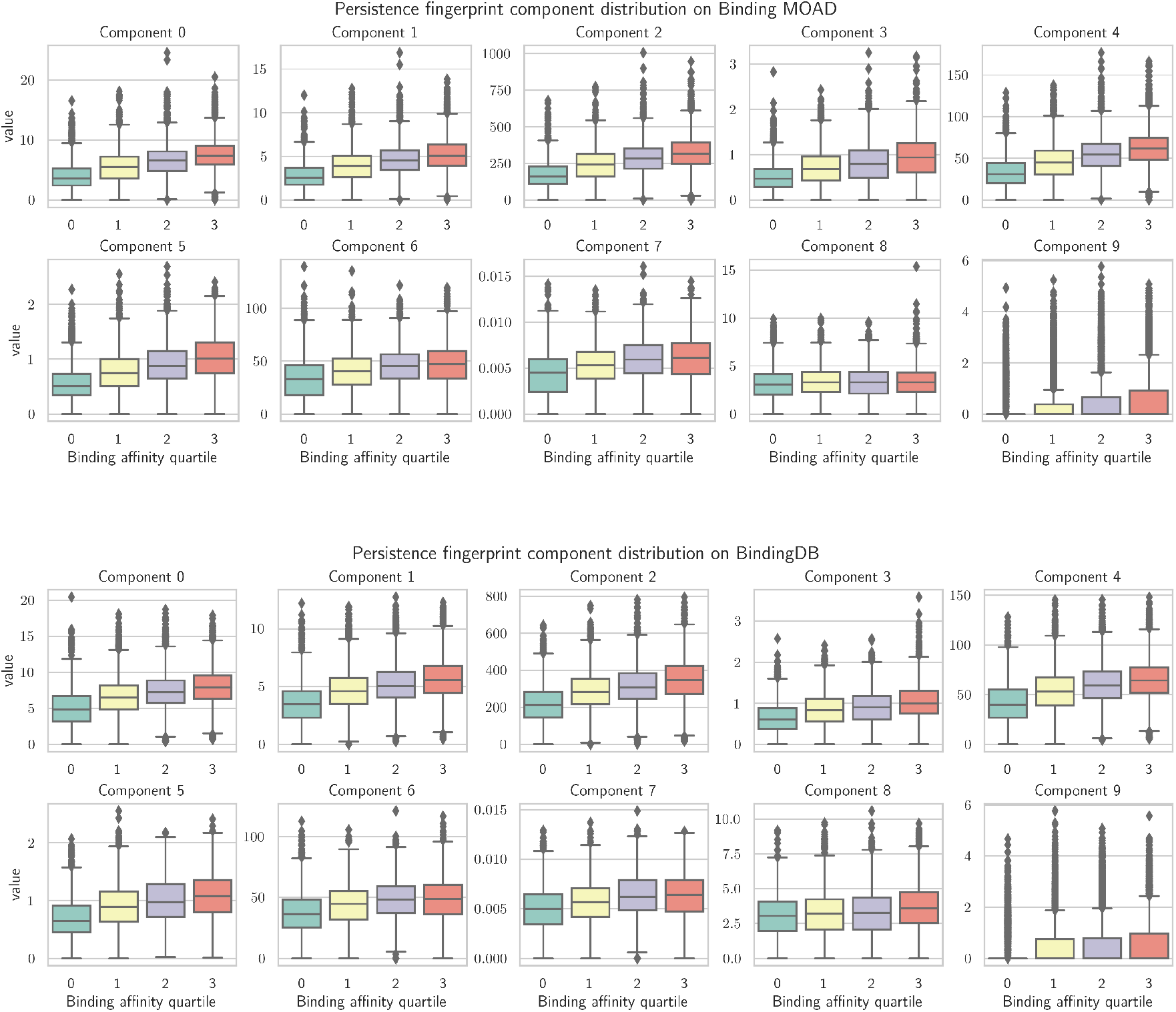
Boxplots of the persistence fingerprint values with respect to different quartiles of binding affinity on the Binding MOAD and BindingDB datasets show that the discriminating power of per-sistence fingerprint generalizes beyond its training dataset (which is PDBBind v2020 refined set, as seen in Figure 12), albeit to a lesser extent for components 8 and 9 of the persistence fingerprint. The *x*-axis and color correspond to quartiles of data by their experimental binding affinity. The *y*-axis corresponds to the values of at that certain persistence fingerprint component.

**Fig. 14.**
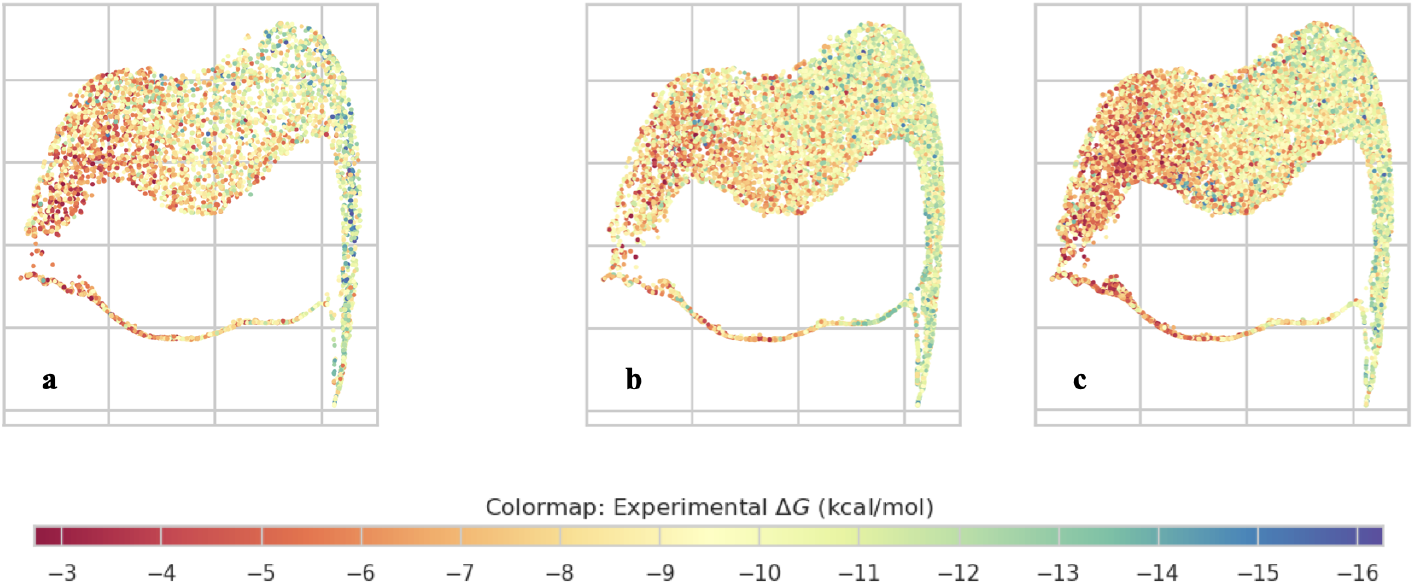
Visualization by PaCMAP [109] shows that persistence fingerprint clusters protein-ligand complexes with similar binding affinity reasonably well, even beyond the training dataset (PDBBind v2020 refined set). The *x*- and *y*-axes are the dimensionality reduced axes from PaCMAP. The color of each point is the experimental binding affinity of the protein-ligand complex. ^**a**^PaCMAP of the persistence fingerprints of the PDBBind v2020 refined set (training set), ^**b**^Binding MOAD dataset, and ^**c**^BindingDB dataset.

As expected, persistence fingerprint shows discriminating power on the PDBBind v2020 refined set (the training set). The remarkable observation is that the discriminating power of persistence fingerprint remains in the Binding MOAD and BindingDB datasets, which consist of more protein-ligand complexes than PDBBind v2020 refined set and the complexes are selected using different methodologies from PDBBind, albeit the discriminating power is to a lesser extent for components 8 and 9 of the fingerprint (Figures 12 and 13).

We also measured the generalizability of persistence fingerprint beyond the training dataset (Figure 14). We computed persistence fingerprints on two protein-ligand binding affinity datasets from the BioLiP database [116, 114], namely Binding MOAD [43, 3, 99, 103] and BindingDB [37, 60], which consist of protein-ligand complexes with binding affinity (Δ*G*) in the range −2.7 to −16.3 kcal/mol.

To visualize the distribution of persistence fingerprint, we used PaCMAP [109], a recent dimensionality reduction technique that down-projects higher dimensional data points onto a two-dimensional manifold, similar to UMAP [72] and t-SNE [68], which additionally preserves “both local and global structure of the data in original space”. We used PaCMAP [109] to down-project persistence fingerprints of the protein-ligand complexes in the PDBBind v2020 refined set [107] (training set), Binding MOAD, and BindingDB datasets. The results are shown in Figure 14. In these plots, each point represents a protein-ligand complex in the dataset, the position of each point is the projection of its persistence fingerprint, and points are colored according to its experimental binding affinity. In Figure 14, we see that binding affinity is separated by persistence fingerprint, as evidenced by a gradient of colors, meaning that complexes with similar binding affinities are similar to each other in persistence fingerprint. These plots show that the representational power of persistence fingerprints generalizes beyond its dataset.

### C. Computational complexity of persistence fingerprint

#### C.1 Computational complexity of IPCs

Persistent homology is computationally expensive to find. In particular, the *k*-skeleton of a Rips complex (a subcomplex with simplex dimension up to *k*) has 𝒪 (*n*^*k*+1^) simplices, where *n* is the number of points in the input point cloud [18]. The time complexity to compute persistent homology is then 𝒪 ((*n*^*k*+1) *ω*^) = 𝒪 (*n*^(*k*+1)*ω*^), where *ω* ≈ 2.4 is the matrix multiplication exponent [10, 53, 78]. For a protein-ligand complex, let *m* be the number of ligand atoms and *n* be the number of protein atoms. When computing the IPCs from whence persistence fingerprint is derived, both cases for *k* ∈ {0, 1} need to be computed and the computational complexity is 𝒪 ((*m* + *n*)^4.8^). Previous binding affinity prediction algorithms [106, 13, 12] that use persistent homology all calculated the the full persistence diagram for at least *k* ∈ {0, 1}, hence they all have computational complexity at least 𝒪 ((*m* + *n*)^4.8^).

Below in Appendix Sections C.2 and C.3, we show an effective algorithm to compute a provable approximation of persistence fingerprint.

#### C.2 Computational complexity of our provably *ε*-accurate approximation algorithm for persistence fingerprint is independent of protein size

By Appendix Section C.1, constructing the full IPC for identifying the components of persistence fingerprint naively takes 𝒪 (*n*^4.8^) time for each protein-ligand complex, where *n* is the number of atoms in the protein-ligand complex. However, in many biologically relevant protein-ligand complexes [17, 27, 42], *n* can be very large, resulting in unwieldy runtime and space consumption by these algorithms. In this subsection, we prove Theorem 1:

##### Theorem 1

**(Complexity of approximating persistence fingerprint)**. *Assume there exists a fixed lower bound on interatomic distances in a protein-ligand complex. Let the number of protein atoms be n, the number of ligand atoms be m, and ω* ≈ 2.4 *be the matrix multiplication exponent [53]. For any* 0 *< ε <* 1, *after an* 𝒪 (*mn* log(*mn*)) *preprocessing procedure, we can compute an approximation to the persistence fingerprint in* 𝒪 (*m* log^6*ω*^(*m/ε*)) *time, independent of protein size, such that the maximum difference between each component in this approximation and that of the corresponding element in the true persistence fingerprint is less than ε*.

First, recall that in the hard sphere model^6^, each atom is modeled by a hard sphere with the atom’s van der Waals radius as the sphere’s radius. Let *r*_*m*_ be the smallest van der Waals radius among all atoms. Let 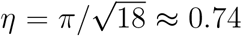 be the maximum packing density of congruent spheres in three dimensions [39]. Then if we denote *a* to be the number of atoms in a ball of radius *r*, we have 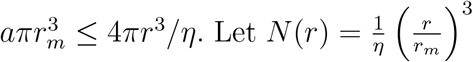 be the maximum number of atoms that can be packed within a ball of radius *r* such that each atom’s “hard sphere” is in this ball of radius *r*, we have *a* ≤ *N* (*r*).

We claim that we can remove protein atoms that are more than a certain distance *r*_*ε*_ from the ligand while erring from the true value of that component by less than *ε*, and *r*_*ε*_ can be set independent of the protein size or protein shape. This removal will reduce the computation time of persistence fingerprint to become independent of protein size. In the following proof, we only prove the case for the persistence fingerprint components that result from 1D IPCs, and the proof for removing atoms in optimizing 0D IPC is easily obtained *mutatis mutandis*.

Suppose we remove (i.e., prune) all protein atoms beyond a certain radius *R* of a certain ligand atom *l* ∈ *L*, we want to find the resulting difference in the contribution to a component of the persistence fingerprint, which is 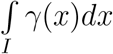 over a certain closed interval *I* = [*i*_*l*_, *i*_*r*_] ⊂ℝ. This interval *I* is given by the table of features of persistence fingerprint (Table 2) and has a fixed width Δ*i* = *i*_*r*_ − *i*_*l*_ = 0.5. Fix *l*. Define the protein atoms within the radius *r* be *P*_*r*_ = {*p* ∈ *P* : *d*(*p, l*) ≤ *r*} and denote the protein atoms to be pruned 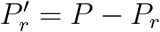. Additionally, provide the constraint that *r > i*_*r*_. Then for a given atom 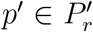, the contribution of *p*^*′*^ in this integral, denoted *c*(*p*^*′*^) with *c* : *P* → ℝ, satisfies

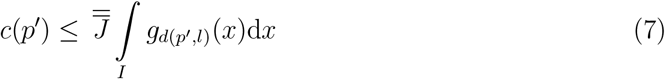

where

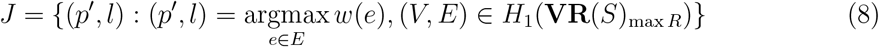

is the set of 1-cycles whose longest edge is the (*p*^*′*^, *l*), and where each generator of the first homology group, a 1-cycle, is viewed as a bipartite graph (Figure 9). Eq. (7) follows from Lemma 2. For convenience, let ||*p*^*′*^|| = *d*(*p*^*′*^, *l*) and 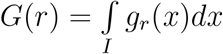.

##### Lemma 3

(Maximum number of 1D holes from opposition distance). *Consider the Vietoris-Rips filtration of a protein-ligand complex with the opposition distance d*_*op*_. *Let the set of protein atoms be P and the set of ligand atoms be L. Suppose* 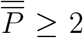 *and* 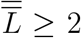 *Then at any filtration parameter* 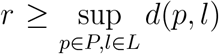 *the homology group in the simplicial complex of the filtration with parameter r has rank* 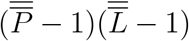.

**Proof of Lemma 3:** Let the simplicial complex at 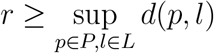 be *K* = **VR**(*S*)_*r*_. Then *K* as a vector space has basis {[*p*_*i*_, *l*_*j*_] : *p*_*i*_ ∈ *P, l*_*j*_ ∈ *L*} ∪ {[*p*_*i*_] : *p*_*i*_ ∈ *P*} ∪ {[*l*_*j*_] : *l*_*j*_ ∈ *L*}. Consider, for example, the case where 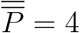 and 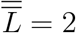 Then *K* looks like

By Lemma 1, *H*_1_(*K*) ≅ *Z*_1_(*K*) = ker(∂_1_(*K*)). And

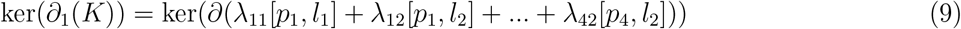

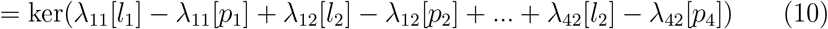

where *λ*_*ij*_ is the coefficient of the generator [*p*_*i*_, *l*_*j*_]. We are led to solve the following system of equations:

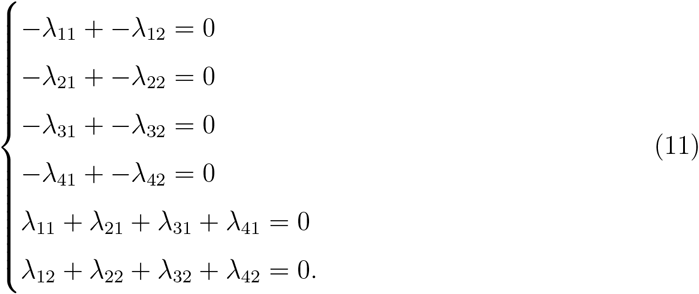

**Fig. 15.**
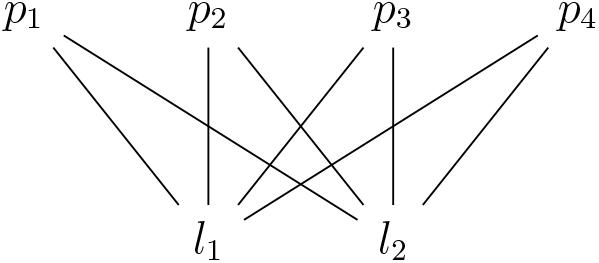
Example of *K* for *P* = {*p*_1_, *p*_2_, *p*_3_, *p*_4_} and *L* = {*l*_1_, *l*_2_}.

In this system of equations (Eq. (11)), the first four equations are generated by each of the protein atoms and the last two equations are generated by each of the ligand atoms. Note that exactly one equation is redundant, so this system of equations yields 5 linearly independent constraints. Since there are 8 variables, given 5 constraints then there are 8 − 5 = 3 free variables, which is the rank of *H*_1_(*K*).

This reasoning generalizes to any arbitrary number of protein atoms and ligand atoms. Suppose *K* has *n* protein atoms and *m* ligand atoms, then

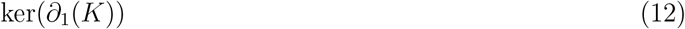

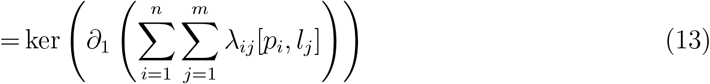

which generates the system of equations

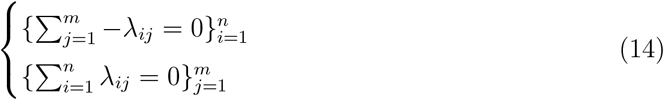

Similar to before, this system of *m* + *n* equations has exactly one redundant equation. Subtracting the number of constraints from the number of variables, we get that the number of free variables to be *mn* − (*m* + *n* − 1) = (*m* − 1)(*n* − 1), which is the rank of *H*_1_(*K*). □

Hence

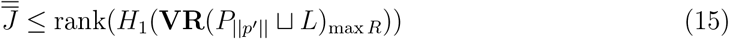

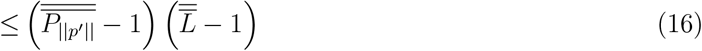

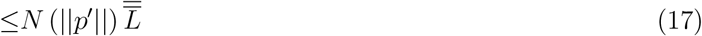

where the protein-ligand complex after pruning all protein atoms farther from *l* than *p*^*′*^ is denoted *P*_||*p*_*′*|| ⊔ *L*. The inequality of Eq. (15) follows from the fact that a 1-cycle formed with any protein atom 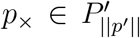 must have its longest edge longer than *d*(*p*^*′*^, *l*) because *d*(*p*_*×*_, *l*) *> d*(*p*^*′*^, *l*). The inequality of Eq. (16) follows from Lemma 3.

We can simplify Eq. (7):

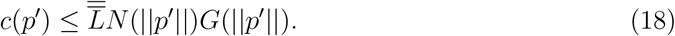

Now we have a bound on *c*(*p*^*′*^). Then, the total effect of all the atoms pruned out on the persistence fingerprint component of concern is given by

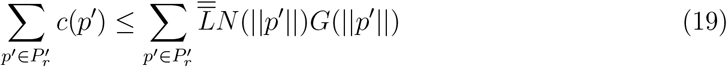

where 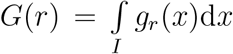 is the integral of *g*_*r*_ is the Gaussian function centered at *r* with standard deviation *σ*, integrated over the interval *I* = [*i*_*l*_, *i*_*r*_].

##### Lemma 4

(Monotonicity and bound of *G*(*r*)*N* ^2^(*r*)). *There exists r*_0_ *such that for all r > r*_0_,

1. *G*(*r*)*N* ^2^(*r*), *G*(*r*)*N* (*r*), *and G*(*r*) *monotonically decrease*.
2. *G*(*r*)*N* ^2^(*r*) *<* exp(−*r*).

*and r*_0_ *does not depend on ε*, 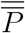 *or* 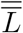.

**Proof of Lemma 4:** Recall that *N* (*r*) is a cubic polynomial in *r*. Intuitively, *g*_*r*_(*x*) decays exponentially and is integrated over an interval [*i*_*l*_, *i*_*r*_] with a fixed width Δ*i* to yield *G*, so asymptotically as *r* → ∞, *G*(*r*)*N* ^2^(*r*) behaves like 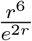, which tends to 0 and decays faster than exp(−*r*). To be precise, consider any *r > i*_*r*_, we have

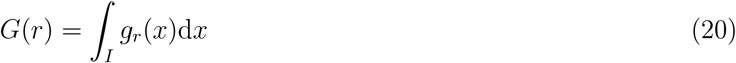

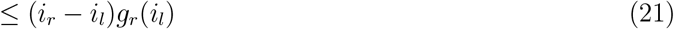

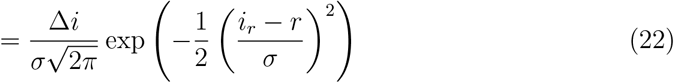

where the inequality in Eq. (21) follows from the assumption that *r > i*_*r*_ and hence *g*_*r*_(*x*) is monotonically decreasing in *I*.

Then to show that there exists *r*_0_ such that *G*(*r*)*N* ^2^(*r*) *<* 0 is strictly decreasing beyond *r*_0_:

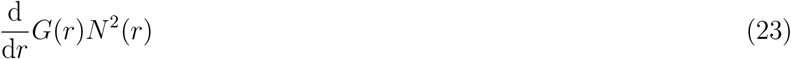

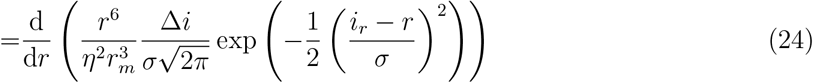

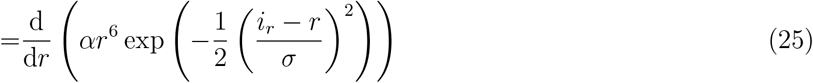

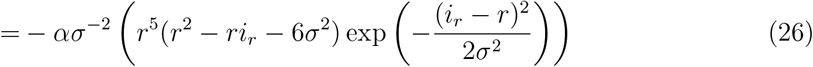

where 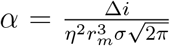 is a positive constant. Since *ασ*^−2^ and 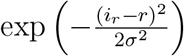 are positive and there exists *r*_1_ such that *r*^5^(*r*^2^ − *ri*_*r*_ − 6*σ*^2^) is positive for all *r > r*_1_, we have that Eq. (26) is negative for all *r > r*_1_. In other words, *G*(*r*)*N* ^2^(*r*) monotonically decreases beyond *r*_1_.

Similarly, there exists *r*_2_ such that *G*(*r*)*N* (*r*) monotonically decreases for all *r > r*_2_. Thus choosing *r >* max(*r*_1_, *r*_2_) satisfies the first claim of Lemma 4.

To show the second claim in Lemma 4, we first show that there exists *r*_3_ such that for all *r > r*_3_, we have

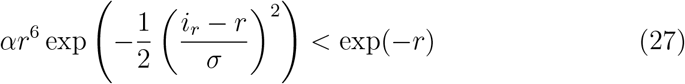

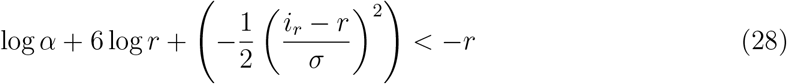

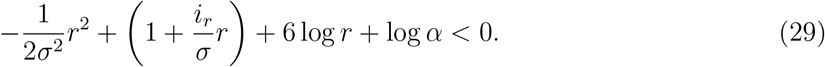

Since the left hand side of Eq. (29) is dominated by 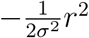, there exists *r*_3_ such that this inequality is true for any *r > r*_3_, which satisfies the second claim of Lemma 4.

Finally note that *i*_*r*_, *r*_1_, *r*_2_, *r*_3_ depend only on the integration interval *I* in *G*(*r*), and not on *ε*, 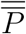, or 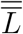 Therefore, for *r*_0_ = max(*i*_*r*_, *r*_1_, *r*_2_, *r*_3_), Lemma 4 is satisfied. □

Choose a pruning radius *r > r*_0_. We claim that for any number and any placement of protein atoms, the total effect of pruning out all atoms in 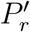, as described in 19, is finite.

Consider a partition of 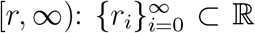 starting from a pruning radius *r* such that *r* = *r*_0_ ≤ *r*_1_ ≤ *r*_2_ ≤ … and ∀*i* ∈ ℤ^+^ : *r*_*i*+1_ − *r*_*i*_ = Δ*r* for some fixed value Δ*r*. Then note that in each interval [*r*_*i*_, *r*_*i*+1_], there can at most be *N* (*r*_*i*+1_) −*N* (*r*_*i*_) protein atoms. Then the total contribution of atoms in [*r*_*i*_, *r*_*i*+1_] to our persistence fingerprint can be bounded component:

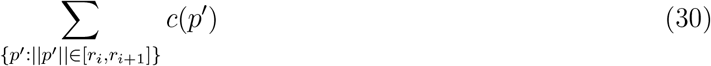

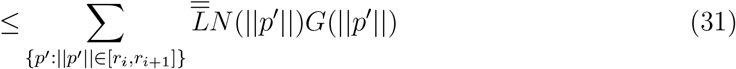

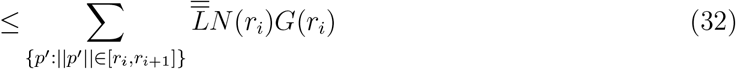

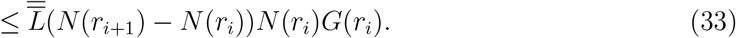

Then the total contribution of all pruned atoms (i.e.,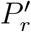) to our persistence fingerprint component is

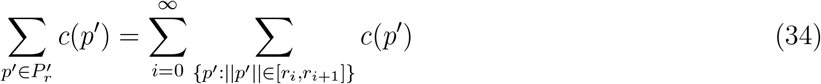

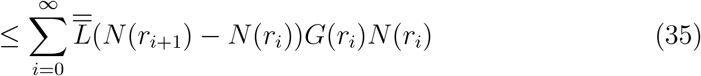

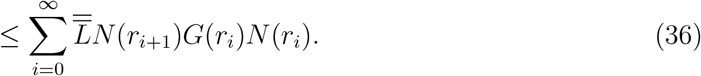

By assumption, 1*/G*(*r*) ∈ 𝒪 (*e*^*r*^) and recalling that *N* (*r*) ∈ 𝒪 (*r*^3^), the last sum (i.e., Eq. (36)) converges. In other words, for any pruning radius *r*, the error resulting from pruning all protein atoms beyond *r* is bounded. Since the pruning error monotonically decreases with respect to increasing *r*, we conclude that given any *ε*, there exists *r*_*ε*_ such that for any pruning radius *r > r*_*ε*_, the error in calculating a persistence fingerprint component of removing all atoms in 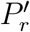 will be less than *ε*, independent of the protein size.

We just showed that for a given ligand atom *l* and a given persistence fingerprint component, there exists a pruning radius *r*_*ε*_ independent of protein size such that when all protein atoms farther than *r*_*ε*_ from *l* are removed, the error in the contribution of *l* in this persistence fingeprint component is less than *ε*. Since each persistence fingerprint component is a sum of contributions of each ligand atom, we can extend our result easily. Let 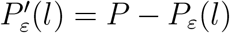 be the set of protein atoms that can be pruned for a given ligand atom *l* ∈ *L* with an error less than *ε*. Then for a ligand 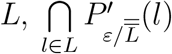 can be pruned while guaranteeing a total pruning error of less than *ε*. The remaining atoms after pruning are contained in a finite volume, thus there is an upper bound for the number of atoms. This bound on pruning radius is only dependent on the number of ligand atoms and *ε*, which guarantees an upper bound on the runtime of persistence fingerprint for a fixed ligand and an *ε*.

#### C.3 Computational complexity with respect to ligand size and *ε*

In Appendix Section C.2, we showed that using the approximation scheme we devised, complexity of computing the persistence fingerprint can be independent of protein size. In this subsection, we derive computational complexity of this approximation of persistence fingerprint in terms of the number of atoms in the ligand 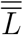 and the error bound *ε*. First, following Eq. (36), we have that for a certain ligand atom and a certain set of protein atoms 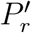 to be removed, the total pruning error is bounded by

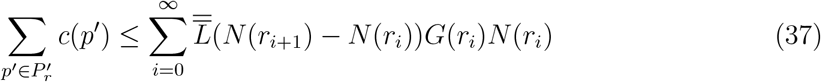

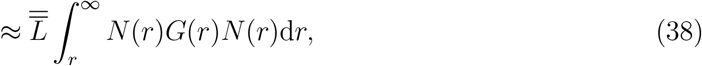

where the approximation in Eq. (38) follows when [*R*, ∞) is partitioned finely (i.e., (*r*_*i*+1_ − *r*_*i*_) → 0). Then given any *ε* bound on error, we want *r*_*ε*_ to satisfy

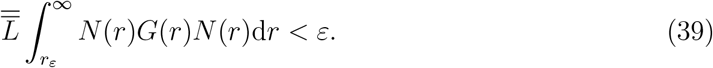

By Lemma 4, there is *r*_0_ independent of *L* or *ε* such that for all *r > r*_0_, we have *N* (*r*)*G*(*r*)*N* (*r*) *<* exp(−*r*). Then for any 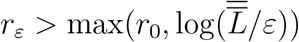, we have

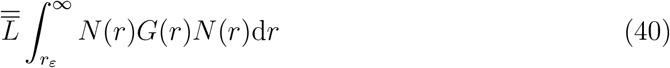

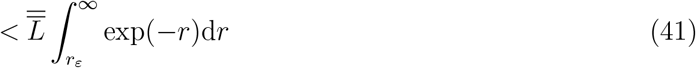

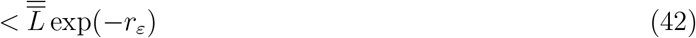

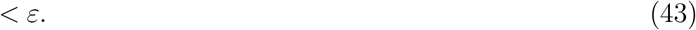

Therefore, any pruning radius 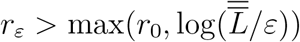 yields a pruning error of less than *ε*.

Then the number of atoms that are included to calculate the persistence fingerprint contribution of this particular ligand atom after pruning is at most 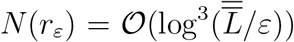. We must compute each ligand atom’s contribution to the persistence fingerprint, and there are 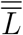 ligand atoms, so the runtime for computing the persistence fingerprint on this pruned complex is:

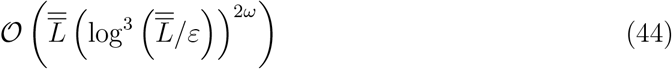

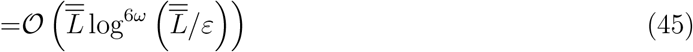

where *ω* ≈ 2.4 is the matrix multiplication exponent [53], 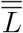 is the number of atoms in the ligand and *ε* is the error bound.

In summary, letting 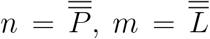, computing an *ε*-accurate approximation to each component of persistence fingerprint entails the following steps:

1. Preprocessing. First, compute and sort the pairwise distances between the protein and the ligand. Then, prune all protein atoms that are at least a distance *r*_*ε*_ away from any ligand atom. Preprocessing takes *𝒪* (*mn* log(*mn*)) time due to sorting.
2. Compute the persistent homology of this pruned complex and calculate the persistence fingerprint. We have shown this approximation to be *ε*-accurate (Eq. (43)). This step takes *𝒪* (*m* log^6*ω*^(*m/ε*)) time.

Since the number of components in the persistence fingerprint is constant (all 10 components are shown in Table 2 in the Main MS), the runtime of computing the entire persistence fingerprint is 𝒪 (*mn* log(*mn*)) for preprocessing and pruning, and 𝒪 (*m* log^6*ω*^(*m/ε*)) to compute the persistent homology of the pruned complex. ◼

#### C.4 Empirical running times

Computation of persistence fingerprint using the pruning method described in previous sections was tested by calculating persistence fingerprint of protein-ligand complexes in the BioLiP dataset [114], which consists of entries from PDBBind [107], Binding MOAD [103], and Binding DB [37]. For this test, the pruning radius *r* is arbitrarily set to be 15 Angstroms.

Each trial in the runtime measurements was allocated a single core of an Intel Xeon processor and 16GB of RAM. The runtime showed a positive correlation with respect to the size of the ligand and is independent of size of the protein (Figure 16. Pearson correlation coefficient (PCC) of protein size vs computation is −0.19. PCC of ligand size vs computation time is 1.00.), which corroborates Eq. (45). Overall, for the 48469 data points, the mean was 41.35 seconds, standard deviation was 71.20 seconds, with a minimum of 1 second, 25% percentile at 19 seconds, 50% percentile at 26 seconds, 75% percentile at 34 seconds, and a maximum of 785 seconds.

Results for the accuracy of pruning is shown in Figure 17, when compared to computing the full IPC and deriving persistence fingerprint from there. Note that the data points on this plot is a subset of the protein-ligand complexes in the BioLiP dataset can be computed by this approximation algorithm, since the full IPCs were intractable to compute for large protein-ligand complexes (Appendix Section C.1). Number of data points = 45199, mean = 3.16 *×* 10^−11^, standard deviation = 2.87 *×* 10^−9^, the values for the 25%, 50%, and 75% percentiles are all 0.0, with a max of 4.8 *×* 10^−7^.

**Fig. 16.**
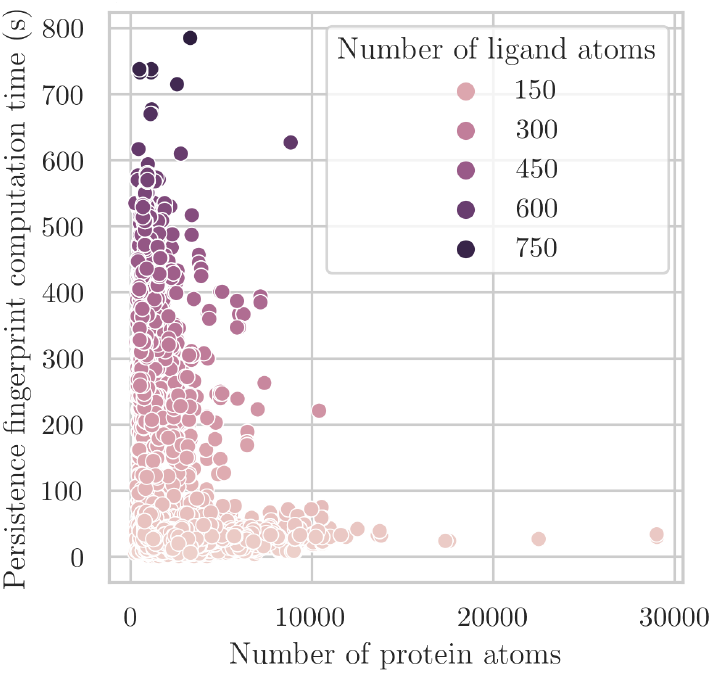
Runtime of computing persistence fingerprint on the BioLiP dataset corroborates Eq. (45) (*n* = 48469). Pearson correlation coefficient (PCC) of protein size vs computation is −0.19, PCC of ligand size vs computation time is 1.00.

### D The decision trees of PATH+

#### D.1 Interpreting the decision trees

Given the small number of decision trees in PATH^+^, we are able to plot and interpret each of the decision trees. The first decision tree is shown in Figure 19.

Note that the nodes on the decision trees index into persistence fingerprint components with zero-based numbering. Also note the prediction values of the decision trees are expressed in *pK*_*d*_ or *pK*_*i*_. For example, in this decision tree 1 (Figure 19), the top node corresponds to the 5^th^ component of the persistence fingerprint, which corresponds to the approximate number of 1-cycles formed by protein nitrogen atoms and ligand nitrogen atoms around 4 Angstroms. A protein-ligand complex chooses one of the two branches based on a the value of that component.

As an example, two complexes are shown in Figure 18: HIV-1 protease in complex with VX-478 (PDB ID: 1hpv [48]) and humanised monomeric RadA in complex with indazole (PDB ID: 4b2i [95]). Looking at Figure 19, we find that the 1hpv complex heads down the left branch because it has a higher value in the 5^th^ component of its persistence fingerprint, which corresponds to a higher number of carbon-nitrogen bipartite matching around 10 Angstroms, while the 4b2i complex heads down the right branch due to a lower value in the 5^th^ component of its persistence fingerprint.

A final prediction is done using all decision trees from persistence fingerprint. Remarkably, through a subsequent literature review, we discovered the features in persistence fingerprint, which were completely automatically derived, are similar to the “interaction fingerprints” manually constructed in previous works on binding affinity prediction [104, 111].

**Fig. 17.**
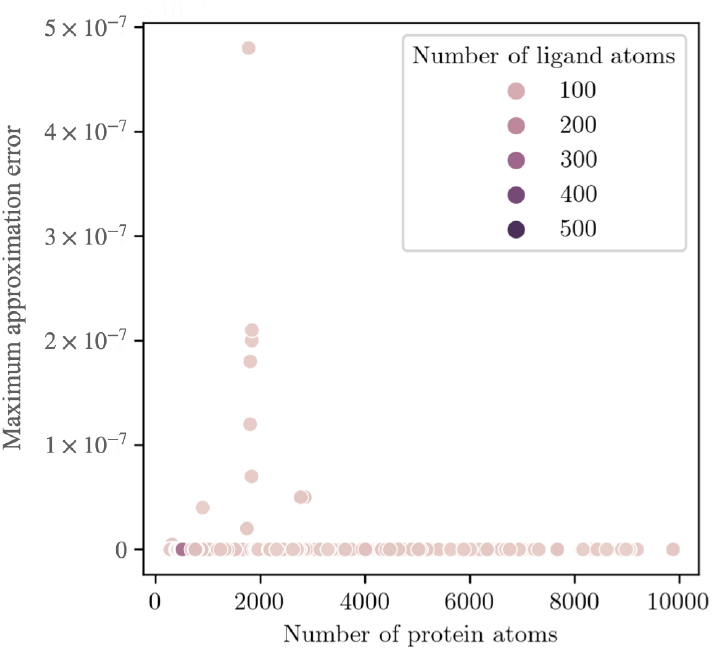
Error from pruning all atoms beyond radius *r* = 15 Å from all ligand atoms is extremely low. Shown: scatter plot of pruning error with respect to number of protein atoms and number of ligand atoms. Note that the data points on this plot is only a subset of the entire BioLiP dataset, since the full IPCs were intractable to compute for large protein-ligand complexes (Appendix Section C.1). Number of data points = 45199, mean = 3.16 *×* 10^−11^, standard deviation = 2.87 *×* 10^−9^, the values for the 25%, 50%, and 75% percentiles are all 0.0, with a max of 4.8 *×* 10^−7^.

Interpretability of PATH provides verification on the robustness beyond simply benchmarking on datasets and provides insights on the geometric features important to predicting binding affinity with persistent homology.

See the full set of decision trees in Appendix Section D.2.

#### D.2 Full Set of 13 Decision Trees

See Figures 20, 21, and 22.

### E. Full benchmarking results

This section shows the performance of PATH and other benchmarked algorithms on predicting binding affinity on PDBBind, BindingDB, and Binding MOAD (Table 7), as well as differentiating between active and decoy ligands on the DUD-E dataset (Table 8) in numeral form. Table 8 shows the AUCs for the protein-ligand complexes for which the various softwares successfully returned a prediction under the following experimental conditions: 1 CPU core, 8GB of memory, and 1 hour of compute time.

**Fig. 18.**
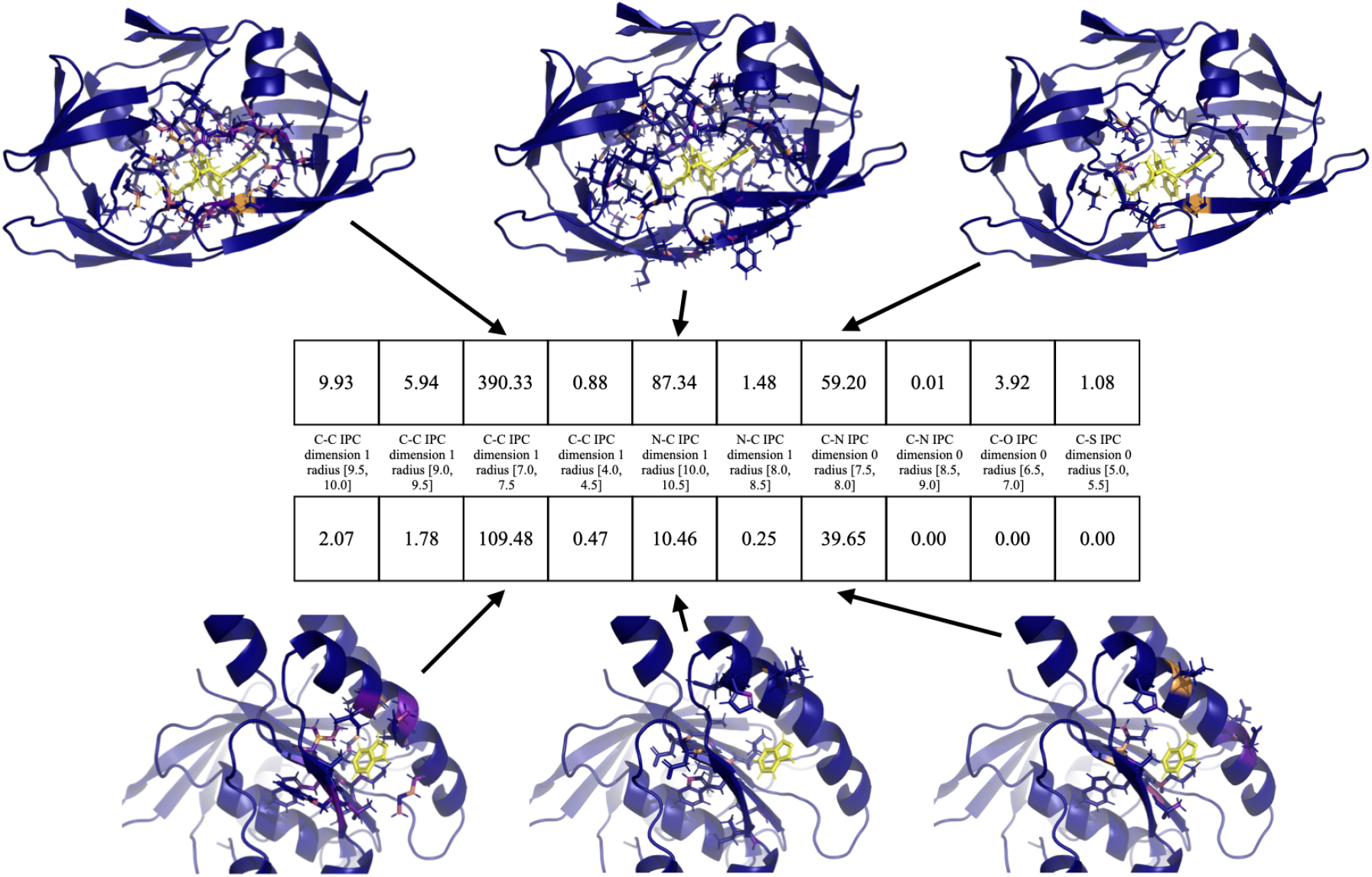
Two protein-ligand complexes shown with their persistence fingerprint. ^Top^HIV-1 protease in complex with VX-478 (PDB ID: 1hpv [48]). ^Bottom^Humanised monomeric RadA in complex with indazole (PDB ID: 4b2i [95]). Contributions to three persistence fingerprint components are shown. The ligand atoms are shown in yellow. Each protein atom is colored according to their contribution to the persistence fingerprint, just like in Figure 5. Each persistence fingerprint component is labeled by the IPC and bin where the IPC is integrated over to yield this component.

**Fig. 19.**
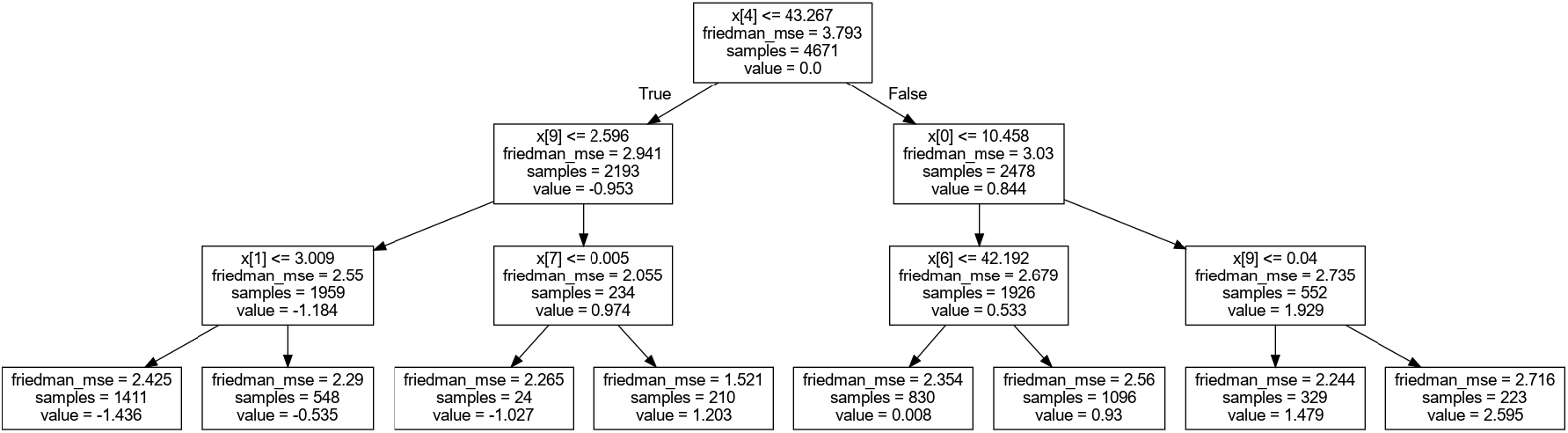
An example decision tree in PATH^+^: Decision tree 1 of 13 in PATH^+^. The “x” at each node is the persistence fingerprint vector, and the indexing which follows is 0-based. For example, the topmost node reads “x[4]”. We can refer to Table 2 to find this corresponds to the nitrogen-carbon bipartite matching at a distance of 10 Å. Given a protein-ligand complex that we wish to predict the binding affinity of, the decision first travels down one of two branches depending on the value of x[4] of the persistence representation. The “value” property of each of the two nodes downstream indicates the average binding affinity of the protein-ligand complexes in the training dataset which reached that node, subtracted by the average binding affinity of the entire dataset.

**Fig. 20.**
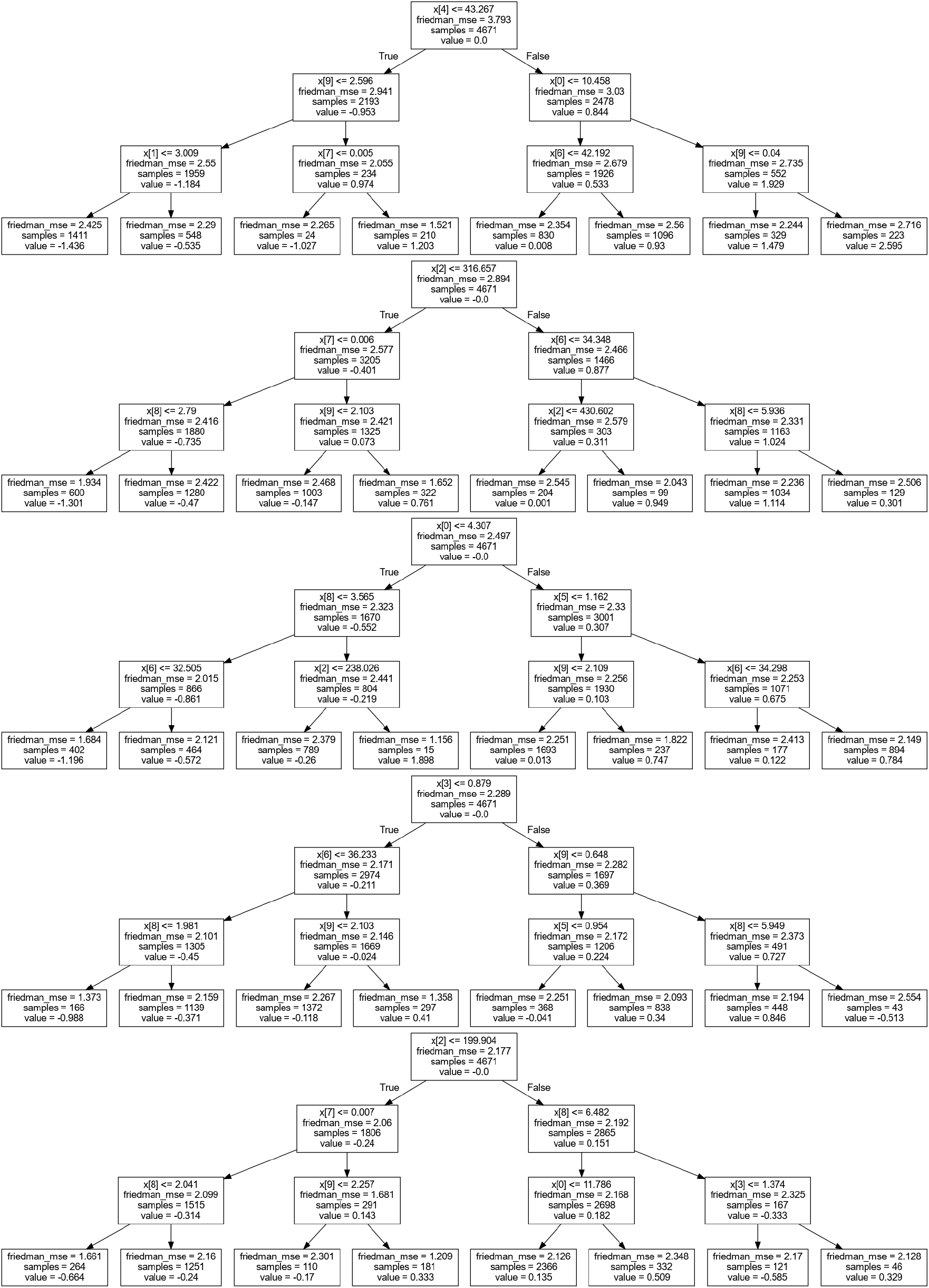
Decision trees 1-5 in PATH^+^

**Fig. 21.**
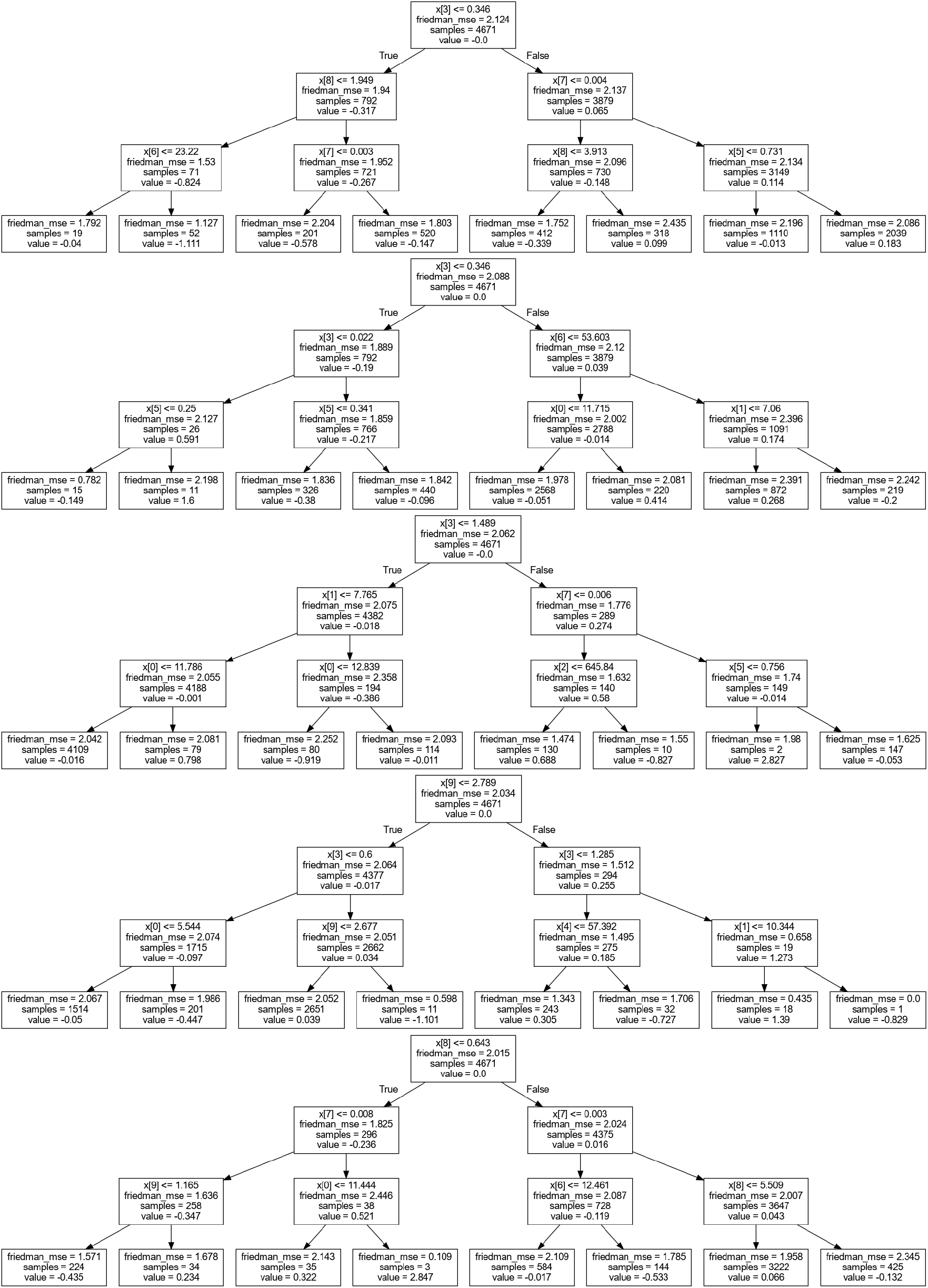
Decision 3tr3ees 6-10 in PATH^+^

**Fig. 22.**
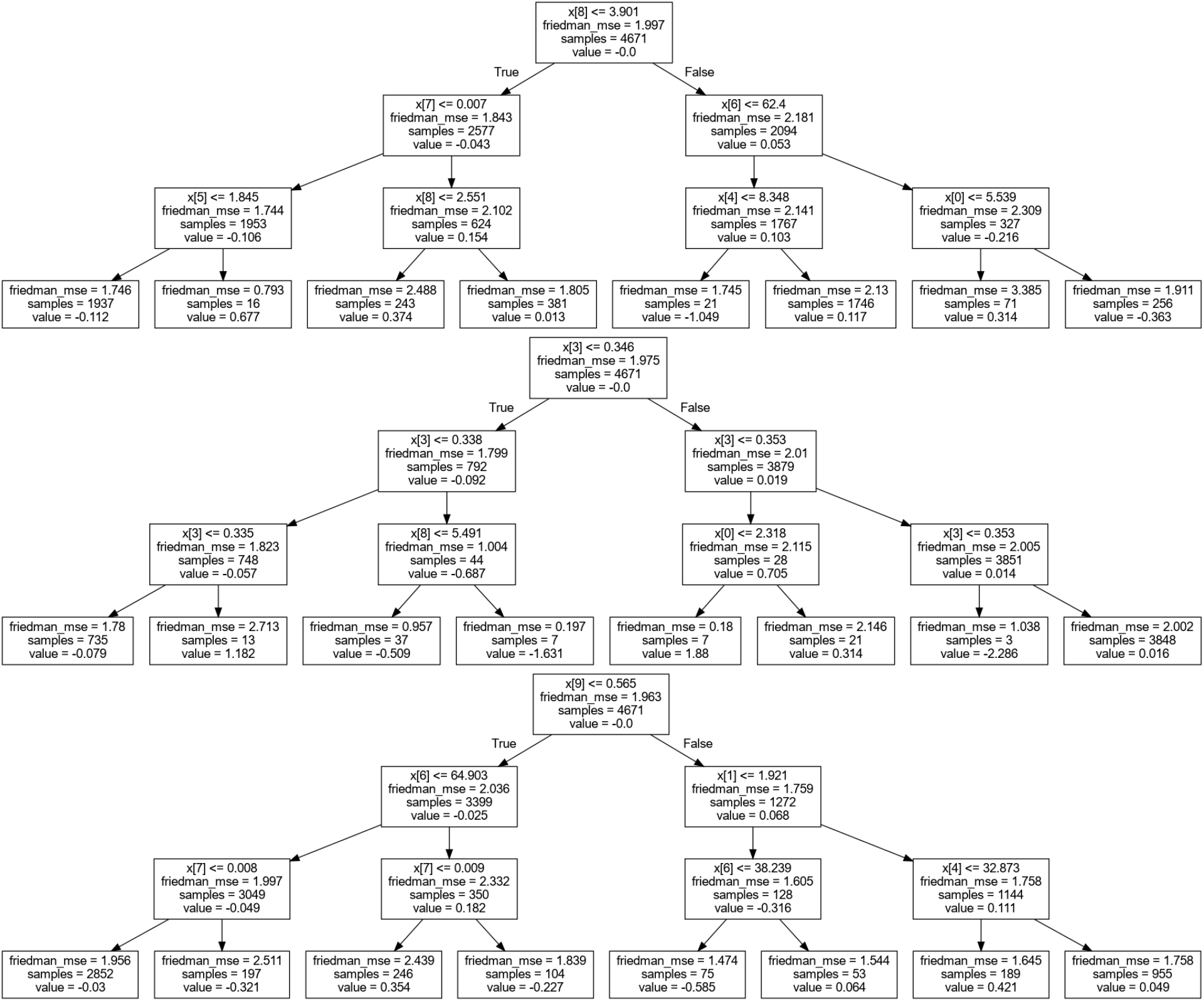
Decision trees 11-13 in PATH^+^

**Table 7.** Δ*G* RMSE (in kcal/mol) of benchmarked binding affinity prediction algorithms on PDBBind, Binding MOAD, and BindingDB.

| Algorithm | PDBBind | BindingDB | Binding MOAD |
| --- | --- | --- | --- |
| AA-Score | 4.41 | 5.15 | 5.23 |
| ADFR | 2.68 | 3.04 | 3.05 |
| GNINA | 1.9 | 2.21 | 2.18 |
| OnionNet | 1.53 | 2.63 | 2.53 |
| PATH<sup>+</sup> | 2.0 | 2.06 | 2.21 |
| PLANET | 1.26 | 2.79 | 2.63 |
| SMoG2016 | 2.27 | 2.41 | 2.46 |
| TNet-BP | 1.68 | 2.24 | 2.39 |
| vinardo | 2.59 | 4.15 | 3.93 |

**Table 8.**
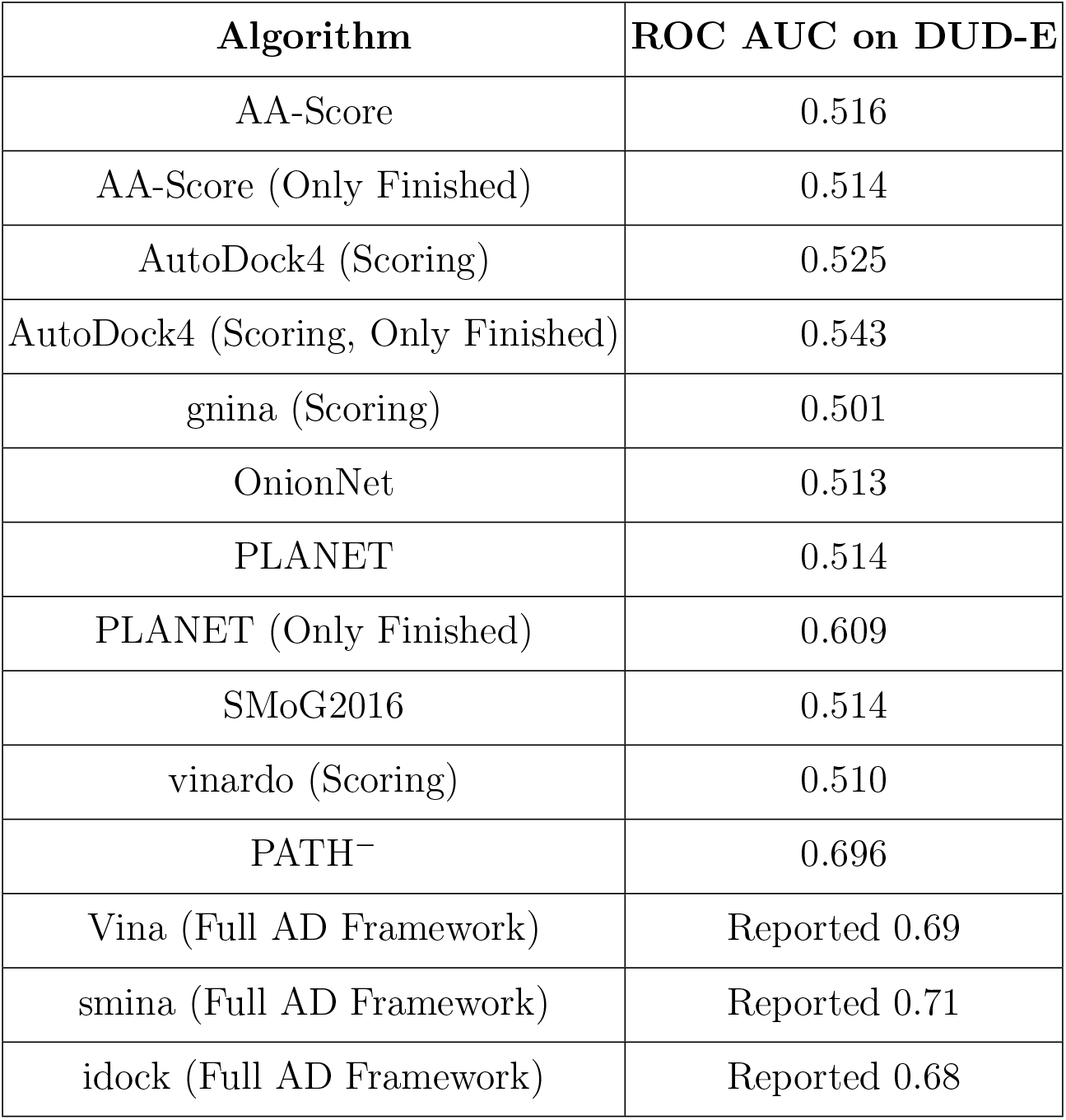
ROC AUCs of binding affinity algorithms on the subset of DUD-E dataset as described in Section 4 and Appendix E. The ROC AUC for Vina, smina, and idock on DUD-E by running the full AutoDock (AD) framework are reported from [71]. “Only finished” indicates that the ROC AUC reported in that row is computed using the protein-ligand complexes for which the row’s software successfully returned a prediction, under the experimental conditions described in Section E.

## Footnotes

3 Edelsbrunner and Harer [31] define the Vietoris-Rips complex with filtration parameter *r* as **VR**_*r*_ (*S*) = {*σ* ⊂ *S* : diam *σ* ≤ 2*r*}. We use the definition from Zomorodian and Carlsson [122] as it is consistent with our implementation. These two alternative definitions for the Vietoris-Rips complex can be reconciled simply by a rescaling of the filtration parameter.

4 Fixing the persistence fingerprint component also fixes a bin *I* and a dimension (0 or 1) such that this persistence fingerprint component is the integral over *I* of the IPC in that dimension (0 or 1) constructed with the atoms *P* and *L*. This information is not needed for computing IPCs, but will be useful to us when calculating the persistence fingerprint.

5 We also learn that for this component, *I* = [10.0, 10.5], but this information is only needed for construction of persistence fingerprint and will not be needed for construction of IPC.

6 We use the hard sphere model for simplicity of calculation. The result will be identical if we used, say, a space-filling model.

## References

1. Adams, H., Emerson, T., Kirby, M., Neville, R., Peterson, C., Shipman, P., Chepushtanova, S., Hanson, E., Motta, F., Ziegelmeier, L.: Persistence images: A stable vector representation of persistent homology. Journal of Machine Learning Research 18 (2017)

2. Adams, H., Segert, J.: Simplicial complex filtration demonstrations in Mathematica, https://www.math.colostate.edu/adams/research/

3. Ahmed, A., Smith, R.D., Clark, J.J., Dunbar Jr, J.B., Carlson, H.A.: Recent improvements to binding moad: a resource for protein–ligand binding affinities and structures. Nucleic acids research 43(D1), D465–D469 (2015)

4. Aldeghi, M., Gapsys, V., de Groot, B.L.: Accurate estimation of ligand binding affinity changes upon protein mutation. ACS central science 4(12), 1708–1718 (2018)

5. Anand, D.V., Meng, Z., Xia, K., Mu, Y.: Weighted persistent homology for osmolyte molecular aggregation and hydrogen-bonding network analysis. Scientific reports 10(1), 9685 (2020)

6. Anderson, A.C.: The process of structure-based drug design. Chemistry & biology 10(9), 787–797 (2003)

7. Bash, P.A., Singh, U.C., Brown, F.K., Langridge, R., Kollman, P.A.: Calculation of the relative change in binding free energy of a protein-inhibitor complex. Science 235(4788), 574–576 (1987)

8. Batool, M., Ahmad, B., Choi, S.: A structure-based drug discovery paradigm. International journal of molecular sciences 20(11), 2783 (2019)

9. Bauer, U.: Ripser: efficient computation of Vietoris-Rips persistence barcodes. J. Appl. Comput. Topol. 5(3), 391–423 (2021). 10.1007/s41468-021-00071-5, https://doi.org/10.1007/s41468-021-00071-5

10. Boissonnat, J.D., Pritam, S.: Computing persistent homology of flag complexes via strong collapses (2018)

11. Buel, G.R., Walters, K.J.: Can alphafold2 predict the impact of missense mutations on structure? Nature Structural & Molecular Biology 29(1), 1–2 (2022)

12. Cang, Z., Mu, L., Wei, G.W.: Representability of algebraic topology for biomolecules in machine learning based scoring and virtual screening. PLoS computational biology 14(1), e1005929 (2018)

13. Cang, Z., Wei, G.W.: Topologynet: Topology based deep convolutional and multi-task neural networks for biomolecular property predictions. PLoS computational biology 13(7), e1005690 (2017)

14. Cang, Z., Wei, G.W.: Integration of element specific persistent homology and machine learning for protein-ligand binding affinity prediction. International journal for numerical methods in biomedical engineering 34(2), e2914 (2018)

15. Chazal, F., Cohen-Steiner, D., Glisse, M., Guibas, L.J., Oudot, S.Y.: Proximity of persistence modules and their diagrams. In: Proceedings of the twenty-fifth annual symposium on Computational geometry. pp. 237–246 (2009)

16. Chazal, F., De Silva, V., Oudot, S.: Persistence stability for geometric complexes. Geometriae Dedicata 173(1), 193–214 (2014)

17. Chen, C.Y., Georgiev, I., Anderson, A.C., Donald, B.R.: Computational structure-based redesign of enzyme activity. Proceedings of the National Academy of Sciences 106(10), 3764–3769 (2009)

18. Choudhary, A., Kerber, M., Raghvendra, S.: Improved approximate rips filtrations with shifted integer lattices and cubical complexes. Journal of Applied and Computational Topology 5(3), 425–458 (2021)

19. Cohen-Steiner, D., Edelsbrunner, H., Harer, J.: Stability of persistence diagrams. In: Proceedings of the twenty-first annual symposium on Computational geometry. pp. 263–271 (2005)

20. wwPDB consortium: Protein Data Bank: the single global archive for 3D macromolecular structure data. Nucleic Acids Research 47(D1), D520–D528 (10 2018). 10.1093/nar/gky949, https://doi.org/10.1093/nar/gky949

21. Cufar, M., Virk, Ž.: Fast computation of persistent homology representatives with involuted persistent homology. arXiv preprint arXiv:2105.03629 (2021)

22. David, V., Grinberg, N., Moldoveanu, S.C., Grinberg, N., Moldoveanu, S.: Long-range molecular interactions involved in the retention mechanisms of liquid chromatography. Advances in chromatography pp. 73–110 (2017)

23. Debroise, T., Shakhnovich, E.I., Chéron, N.: A hybrid knowledge-based and empirical scoring function for protein–ligand interaction: Smog2016. Journal of chemical information and modeling 57(3), 584–593 (2017)

24. Deng, L., Sui, Y., Zhang, J.: Xgbprh: prediction of binding hot spots at protein–rna interfaces utilizing extreme gradient boosting. Genes 10(3), 242 (2019)

25. DePristo, M.A., de Bakker, P.I., Blundell, T.L.: Heterogeneity and inaccuracy in protein structures solved by x-ray crystallography. Structure 12(5), 831–838 (2004)

26. Dlotko, P.: Persistence representations. In: GUDHI User and Reference Manual. GUDHI Editorial Board (2017)

27. Do Kwon, Y., Pancera, M., Acharya, P., Georgiev, I.S., Crooks, E.T., Gorman, J., Joyce, M.G., Guttman, M., Ma, X., Narpala, S., et al.: Crystal structure, conformational fixation and entry-related interactions of mature ligand-free hiv-1 env. Nature structural & molecular biology 22(7), 522–531 (2015)

28. Donald, B.R.: Algorithms in structural molecular biology. MIT Press (2023)

29. Eberhardt, J., Santos-Martins, D., Tillack, A.F., Forli, S.: Autodock vina 1.2. 0: New docking methods, expanded force field, and python bindings. Journal of chemical information and modeling 61(8), 3891–3898 (2021)

30. Edelsbrunner, H., Harer, J., et al.: Persistent homology-a survey. Contemporary mathematics 453(26), 257–282 (2008)

31. Edelsbrunner, H., Harer, J.L.: Computational Topology: An Introduction. American Mathematical Society, hardcover edn. (2009)

32. Fan, C., Liu, D., Huang, R., Chen, Z., Deng, L.: Predrsa: a gradient boosted regression trees approach for predicting protein solvent accessibility. In: Bmc Bioinformatics. vol. 17, pp. 85–95. BioMed Central (2016)

33. Fasy, B.T., Patel, A.: Persistent homology transform cosheaf. arXiv preprint arXiv:2208.05243 (2022)

34. Friedman, J.H.: Greedy function approximation: a gradient boosting machine. Annals of statistics pp. 1189–1232 (2001)

35. Fugacci, U., Scaramuccia, S., Iuricich, F., De Floriani, L., et al.: Persistent homology: a step-by-step introduction for newcomers. In: STAG. pp. 1–10 (2016)

36. Gaieb, Z., Parks, C.D., Chiu, M., Yang, H., Shao, C., Walters, W.P., Lambert, M.H., Nevins, N., Bembenek, S.D., Ameriks, M.K., et al.: D3r grand challenge 3: blind prediction of protein–ligand poses and affinity rankings. Journal of computer-aided molecular design 33, 1–18 (2019)

37. Gilson, M.K., Liu, T., Baitaluk, M., Nicola, G., Hwang, L., Chong, J.: Bindingdb in 2015: a public database for medicinal chemistry, computational chemistry and systems pharmacology. Nucleic acids research 44(D1), D1045–D1053 (2016)

38. Gorczynski, M.J., Grembecka, J., Zhou, Y., Kong, Y., Roudaia, L., Douvas, M.G., Newman, M., Bielnicka, I., Baber, G., Corpora, T., et al.: Allosteric inhibition of the protein-protein interaction between the leukemia-associated proteins runx1 and cbfβ. Chemistry & biology 14(10), 1186–1197 (2007)

39. Hales, T.C.: A proof of the kepler conjecture. Annals of mathematics pp. 1065–1185 (2005)

40. Hallen, M.A., Martin, J.W., Ojewole, A., Jou, J.D., Lowegard, A.U., Frenkel, M.S., Gainza, P., Nisonoff, H.M., Mukund, A., Wang, S., et al.: Osprey 3.0: open-source protein redesign for you, with powerful new features. Journal of computational chemistry 39(30), 2494–2507 (2018)

41. Hatcher, A.: Algebraic Topology. Cambridge University Press, Cambridge, England (Dec 2001)

42. Holt, G.T., Gorman, J., Wang, S., Lowegard, A.U., Zhang, B., Liu, T., Lin, B.C., Louder, M.K., Frenkel, M.S., McKee, K., et al.: Improved hiv-1 neutralization breadth and potency of v2-apex antibodies by in silico design. Cell reports 42(7) (2023)

43. Hu, L., Benson, M.L., Smith, R.D., Lerner, M.G., Carlson, H.A.: Binding moad (mother of all databases). Proteins: Structure, Function, and Bioinformatics 60(3), 333–340 (2005)

44. Jin, Z., Wu, T., Chen, T., Pan, D., Wang, X., Xie, J., Quan, L., Lyu, Q.: Capla: improved prediction of protein– ligand binding affinity by a deep learning approach based on a cross-attention mechanism. Bioinformatics 39(2), btad049 (2023)

45. Jones, D., Kim, H., Zhang, X., Zemla, A., Stevenson, G., Bennett, W.D., Kirshner, D., Wong, S.E., Lightstone, F.C., Allen, J.E.: Improved protein–ligand binding affinity prediction with structure-based deep fusion inference. Journal of chemical information and modeling 61(4), 1583–1592 (2021)

46. Jumper, J., Evans, R., Pritzel, A., Green, T., Figurnov, M., Ronneberger, O., Tunyasuvunakool, K., Bates, R., Žídek, A., Potapenko, A., et al.: Highly accurate protein structure prediction with alphafold. Nature 596(7873), 583–589 (2021)

47. Kanari, L., Dłotko, P., Scolamiero, M., Levi, R., Shillcock, J., Hess, K., Markram, H.: A topological representation of branching neuronal morphologies. Neuroinformatics 16, 3–13 (2018)

48. Kim, E., Baker, C., Dwyer, M., Murcko, M., Rao, B., Tung, R., Navia, M.: Crystal structure of hiv-1 protease in complex with vx-478, a potent and orally bioavailable inhibitor of the enzyme. Journal of the American Chemical Society 117(3), 1181–1182 (1995)

49. Koes, D.R., Baumgartner, M.P., Camacho, C.J.: Lessons learned in empirical scoring with smina from the csar 2011 benchmarking exercise. Journal of chemical information and modeling 53(8), 1893–1904 (2013)

50. Kontoyianni, M.: Docking and virtual screening in drug discovery. Proteomics for drug discovery: Methods and protocols pp. 255–266 (2017)

51. Kovalevsky, A.Y., Tie, Y., Liu, F., Boross, P.I., Wang, Y.F., Leshchenko, S., Ghosh, A.K., Harrison, R.W., Weber, I.T.: Effectiveness of nonpeptide clinical inhibitor tmc-114 on hiv-1 protease with highly drug resistant mutations d30n, i50v, and l90m. Journal of medicinal chemistry 49(4), 1379–1387 (2006)

52. Kryshtafovych, A., Schwede, T., Topf, M., Fidelis, K., Moult, J.: Critical assessment of methods of protein structure prediction (casp)—round xiv. Proteins: Structure, Function, and Bioinformatics 89(12), 1607–1617 (2021)

53. Le Gall, F.: Powers of tensors and fast matrix multiplication. In: Proceedings of the 39th international symposium on symbolic and algebraic computation. pp. 296–303 (2014)

54. Li, H., Leung, K.S., Wong, M.H.: idock: A multithreaded virtual screening tool for flexible ligand docking. In: 2012 IEEE Symposium on Computational Intelligence in Bioinformatics and Computational Biology (CIBCB). pp. 77–84. IEEE (2012)

55. Li, S., Zhou, J., Xu, T., Huang, L., Wang, F., Xiong, H., Huang, W., Dou, D., Xiong, H.: Structure-aware interactive graph neural networks for the prediction of protein-ligand binding affinity. In: Proceedings of the 27th ACM SIGKDD Conference on Knowledge Discovery & Data Mining. pp. 975–985 (2021)

56. Li, S., Xi, L., Wang, C., Li, J., Lei, B., Liu, H., Yao, X.: A novel method for protein-ligand binding affinity prediction and the related descriptors exploration. Journal of computational chemistry 30(6), 900–909 (2009)

57. Li, Y., Han, L., Liu, Z., Wang, R.: Comparative assessment of scoring functions on an updated benchmark: 2. evaluation methods and general results. Journal of chemical information and modeling 54(6), 1717–1736 (2014)

58. Liang, J., Edelsbrunner, H., Fu, P., Sudhakar, P.V., Subramaniam, S.: Analytical shape computation of macro-molecules: Ii. inaccessible cavities in proteins. Proteins: Structure, Function, and Bioinformatics 33(1), 18–29 (1998)

59. Liu, F., Kovalevsky, A.Y., Tie, Y., Ghosh, A.K., Harrison, R.W., Weber, I.T.: Effect of flap mutations on structure of hiv-1 protease and inhibition by saquinavir and darunavir. Journal of molecular biology 381(1), 102–115 (2008)

60. Liu, T., Lin, Y., Wen, X., Jorissen, R.N., Gilson, M.K.: Bindingdb: a web-accessible database of experimentally determined protein–ligand binding affinities. Nucleic acids research 35(suppl_1), D198–D201 (2007)

61. Liu, X., Feng, H., Lü, Z., Xia, K.: Persistent tor-algebra for protein–protein interaction analysis. Briefings in Bioinformatics 24(2), bbad046 (2023)

62. Liu, X., Feng, H., Wu, J., Xia, K.: Dowker complex based machine learning (dcml) models for protein-ligand binding affinity prediction. PLoS Computational Biology 18(4), e1009943 (2022)

63. Liu, X., Feng, H., Wu, J., Xia, K.: Hom-complex-based machine learning (hcml) for the prediction of protein– protein binding affinity changes upon mutation. Journal of chemical information and modeling 62(17), 3961– 3969 (2022)

64. Liu, X., Wang, X., Wu, J., Xia, K.: Hypergraph-based persistent cohomology (hpc) for molecular representations in drug design. Briefings in Bioinformatics 22(5), bbaa411 (2021)

65. Liu, Z., Li, Y., Han, L., Li, J., Liu, J., Zhao, Z., Nie, W., Liu, Y., Wang, R.: Pdb-wide collection of binding data: current status of the pdbbind database. Bioinformatics 31(3), 405–412 (2015)

66. Liu, Z., Su, M., Han, L., Liu, J., Yang, Q., Li, Y., Wang, R.: Forging the basis for developing protein–ligand interaction scoring functions. Accounts of chemical research 50(2), 302–309 (2017)

67. Louppe, G.: Understanding random forests: From theory to practice. arXiv preprint arXiv:1407.7502 (2014)

68. Van der Maaten, L., Hinton, G.: Visualizing data using t-sne. Journal of machine learning research 9(11) (2008)

69. Maia, E.H.B., Assis, L.C., De Oliveira, T.A., Da Silva, A.M., Taranto, A.G.: Structure-based virtual screening: from classical to artificial intelligence. Frontiers in chemistry 8, 343 (2020)

70. Maria, C., Boissonnat, J.D., Glisse, M., Yvinec, M.: The gudhi library: Simplicial complexes and persistent homology. In: Mathematical Software–ICMS 2014: 4th International Congress, Seoul, South Korea, August 5-9, 2014. Proceedings 4. pp. 167–174. Springer (2014)

71. Masters, L., Eagon, S., Heying, M.: Evaluation of consensus scoring methods for autodock vina, smina and idock. Journal of Molecular Graphics and Modelling 96, 107532 (2020)

72. McInnes, L., Healy, J., Melville, J.: Umap: Uniform manifold approximation and projection for dimension reduction. arXiv preprint arXiv:1802.03426 (2018)

73. McNutt, A.T., Francoeur, P., Aggarwal, R., Masuda, T., Meli, R., Ragoza, M., Sunseri, J., Koes, D.R.: Gnina 1.0: molecular docking with deep learning. Journal of cheminformatics 13(1), 43 (2021)

74. Meli, R., Morris, G.M., Biggin, P.C.: Scoring functions for protein-ligand binding affinity prediction using structure-based deep learning: A review. Frontiers in bioinformatics 2, 57 (2022)

75. Menze, B.H., Kelm, B.M., Masuch, R., Himmelreich, U., Bachert, P., Petrich, W., Hamprecht, F.A.: A comparison of random forest and its gini importance with standard chemometric methods for the feature selection and classification of spectral data. BMC bioinformatics 10, 1–16 (2009)

76. Merrick, L.: Randomized ablation feature importance. arXiv preprint arXiv:1910.00174 (2019)

77. Mey, A.S., Allen, B.K., Macdonald, H.E.B., Chodera, J.D., Hahn, D.F., Kuhn, M., Michel, J., Mobley, D.L., Naden, L.N., Prasad, S., et al.: Best practices for alchemical free energy calculations [article v1. 0]. Living journal of computational molecular science 2(1) (2020)

78. Milosavljevic, N., Morozov, D., Skraba, P.: Zigzag persistent homology in matrix multiplication time. In: Proceedings of the twenty-seventh Annual Symposium on Computational Geometry. pp. 216–225 (2011)

79. Morris, G.M., Huey, R., Lindstrom, W., Sanner, M.F., Belew, R.K., Goodsell, D.S., Olson, A.J.: Autodock4 and autodocktools4: Automated docking with selective receptor flexibility. Journal of computational chemistry 30(16), 2785–2791 (2009)

80. Murdoch, W.J., Singh, C., Kumbier, K., Abbasi-Asl, R., Yu, B.: Definitions, methods, and applications in interpretable machine learning. Proceedings of the National Academy of Sciences 116(44), 22071–22080 (2019)

81. Mysinger, M.M., Carchia, M., Irwin, J.J., Shoichet, B.K.: Directory of useful decoys, enhanced (dud-e): better ligands and decoys for better benchmarking. Journal of medicinal chemistry 55(14), 6582–6594 (2012)

82. Nguyen, D.D., Cang, Z., Wu, K., Wang, M., Cao, Y., Wei, G.W.: Mathematical deep learning for pose and binding affinity prediction and ranking in d3r grand challenges. Journal of computer-aided molecular design 33, 71–82 (2019)

83. Pak, M.A., Markhieva, K.A., Novikova, M.S., Petrov, D.S., Vorobyev, I.S., Maksimova, E.S., Kondrashov, F.A., Ivankov, D.N.: Using alphafold to predict the impact of single mutations on protein stability and function. Plos one 18(3), e0282689 (2023)

84. Pan, X., Wang, H., Zhang, Y., Wang, X., Li, C., Ji, C., Zhang, J.Z.: Aa-score: a new scoring function based on amino acid-specific interaction for molecular docking. Journal of Chemical Information and Modeling 62(10), 2499–2509 (2022)

85. Pandala, S.R.: Lazypredict. https://github.com/shankarpandala/lazypredict (2022)

86. Pedregosa, F., Varoquaux, G., Gramfort, A., Michel, V., Thirion, B., Grisel, O., Blondel, M., Prettenhofer, P., Weiss, R., Dubourg, V., Vanderplas, J., Passos, A., Cournapeau, D., Brucher, M., Perrot, M., Duchesnay, E.: Scikit-learn: Machine learning in Python. Journal of Machine Learning Research 12, 2825–2830 (2011)

87. Pérez, J.B., Hauke, S., Lupo, U., Caorsi, M., Dassatti, A.: giotto-ph: A python library for high-performance computation of persistent homology of vietoris–rips filtrations (2021)

88. Qi, Y., Martin, J.W., Barb, A.W., Thélot, F., Yan, A.K., Donald, B.R., Oas, T.G.: Continuous interdomain orientation distributions reveal components of binding thermodynamics. Journal of molecular biology 430(18), 3412–3426 (2018)

89. Quinlan, J.R.: Induction of decision trees. Machine learning 1, 81–106 (1986)

90. Quiroga, R., Villarreal, M.A.: Vinardo: A scoring function based on autodock vina improves scoring, docking, and virtual screening. PloS one 11(5), e0155183 (2016)

91. Ravindranath, P.A., Forli, S., Goodsell, D.S., Olson, A.J., Sanner, M.F.: Autodockfr: advances in protein-ligand docking with explicitly specified binding site flexibility. PLoS computational biology 11(12), e1004586 (2015)

92. Rudicell, R.S., Kwon, Y.D., Ko, S.Y., Pegu, A., Louder, M.K., Georgiev, I.S., Wu, X., Zhu, J., Boyington, J.C., Chen, X., et al.: Enhanced potency of a broadly neutralizing hiv-1 antibody in vitro improves protection against lentiviral infection in vivo. Journal of virology 88(21), 12669–12682 (2014)

93. Rudin, C.: Stop explaining black box machine learning models for high stakes decisions and use interpretable models instead. Nature machine intelligence 1(5), 206–215 (2019)

94. Rudin, C., Chen, C., Chen, Z., Huang, H., Semenova, L., Zhong, C.: Interpretable machine learning: Fundamental principles and 10 grand challenges. Statistic Surveys 16, 1–85 (2022)

95. Scott, D.E., Ehebauer, M.T., Pukala, T., Marsh, M., Blundell, T.L., Venkitaraman, A.R., Abell, C., Hyvönen, M.: Using a fragment-based approach to target protein–protein interactions. ChemBioChem 14(3), 332–342 (2013)

96. Seo, S., Choi, J., Park, S., Ahn, J.: Binding affinity prediction for protein–ligand complex using deep attention mechanism based on intermolecular interactions. BMC bioinformatics 22, 1–15 (2021)

97. Sheehy, D.R.: Linear-size approximations to the vietoris-rips filtration. In: Proceedings of the twenty-eighth annual symposium on Computational geometry. pp. 239–248 (2012)

98. Shoichet, B.K.: Virtual screening of chemical libraries. Nature 432(7019), 862–865 (2004)

99. Smith, R.D., Clark, J.J., Ahmed, A., Orban, Z.J., Dunbar Jr, J.B., Carlson, H.A.: Updates to binding moad (mother of all databases): polypharmacology tools and their utility in drug repurposing. Journal of molecular biology 431(13), 2423–2433 (2019)

100. Stepniewska-Dziubinska, M.M., Zielenkiewicz, P., Siedlecki, P.: Development and evaluation of a deep learning model for protein–ligand binding affinity prediction. Bioinformatics 34(21), 3666–3674 (2018)

101. Tauzin, G., Lupo, U., Tunstall, L., Pérez, J.B., Caorsi, M., Medina-Mardones, A.M., Dassatti, A., Hess, K.: giotto-tda: A topological data analysis toolkit for machine learning and data exploration. The Journal of Machine Learning Research 22(1), 1834–1839 (2021)

102. Vega, S., Kang, L.W., Velazquez-Campoy, A., Kiso, Y., Amzel, L.M., Freire, E.: A structural and thermodynamic escape mechanism from a drug resistant mutation of the hiv-1 protease. Proteins: Structure, Function, and Bioinformatics 55(3), 594–602 (2004)

103. Wagle, S., Smith, R.D., Dominic III, A.J., DasGupta, D., Tripathi, S.K., Carlson, H.A.: Sunsetting binding moad with its last data update and the addition of 3d-ligand polypharmacology tools. Scientific Reports 13(1), 3008 (2023)

104. Wang, D.D., Chan, M.T.: Protein-ligand binding affinity prediction based on profiles of intermolecular contacts. Computational and Structural Biotechnology Journal 20, 1088–1096 (2022)

105. Wang, H., Liu, H., Ning, S., Zeng, C., Zhao, Y.: Dlssaffinity: protein–ligand binding affinity prediction via a deep learning model. Physical Chemistry Chemical Physics 24(17), 10124–10133 (2022)

106. Wang, M., Cang, Z., Wei, G.W.: A topology-based network tree for the prediction of protein–protein binding affinity changes following mutation. Nature Machine Intelligence 2(2), 116–123 (2020)

107. Wang, R., Fang, X., Lu, Y., Wang, S.: The pdbbind database: Collection of binding affinities for protein-ligand complexes with known three-dimensional structures. Journal of medicinal chemistry 47(12), 2977–2980 (2004)

108. Wang, S., Reeve, S.M., Holt, G.T., Ojewole, A.A., Frenkel, M.S., Gainza, P., Keshipeddy, S., Fowler, V.G., Wright, D.L., Donald, B.R.: Chiral evasion and stereospecific antifolate resistance in staphylococcus aureus. PLoS Computational Biology 18(2), e1009855 (2022)

109. Wang, Y., Huang, H., Rudin, C., Shaposhnik, Y.: Understanding how dimension reduction tools work: An empirical approach to deciphering t-sne, umap, trimap, and pacmap for data visualization. Journal of Machine Learning Research 22(201), 1–73 (2021), http://jmlr.org/papers/v22/20-1061.html

110. Wee, J., Xia, K.: Persistent spectral based ensemble learning (perspect-el) for protein–protein binding affinity prediction. Briefings in Bioinformatics 23(2), bbac024 (2022)

111. Wójcikowski, M., Kukiełka, M., Stepniewska-Dziubinska, M.M., Siedlecki, P.: Development of a protein–ligand extended connectivity (plec) fingerprint and its application for binding affinity predictions. Bioinformatics 35(8), 1334–1341 (2019)

112. Wu, K., Zhao, Z., Wang, R., Wei, G.W.: Topp–s: Persistent homology-based multi-task deep neural networks for simultaneous predictions of partition coefficient and aqueous solubility. Journal of computational chemistry 39(20), 1444–1454 (2018)

113. Xin, R., Zhong, C., Chen, Z., Takagi, T., Seltzer, M., Rudin, C.: Exploring the whole rashomon set of sparse decision trees. Advances in Neural Information Processing Systems 35, 14071–14084 (2022)

114. Yang, J., Roy, A., Zhang, Y.: Biolip: a semi-manually curated database for biologically relevant ligand–protein interactions. Nucleic acids research 41(D1), D1096–D1103 (2012)

115. Yi, Y., Wan, X., Zhao, K., Ou-Yang, L., Zhao, P.: Predicting protein-ligand binding affinity with equivariant line graph network. arXiv preprint arXiv:2210.16098 (2022)

116. Zhang, C., Zhang, X., Freddolino, P.L., Zhang, Y.: Biolip2: an updated structure database for biologically relevant ligand–protein interactions. Nucleic Acids Research p. gkad630 (2023)

117. Zhang, R., Xin, R., Seltzer, M., Rudin, C.: Optimal sparse regression trees. In: Proceedings of the AAAI Conference on Artificial Intelligence. vol. 37, pp. 11270–11279 (2023)

118. Zhang, X., Gao, H., Wang, H., Chen, Z., Zhang, Z., Chen, X., Li, Y., Qi, Y., Wang, R.: Planet: a multi-objective graph neural network model for protein–ligand binding affinity prediction. Journal of Chemical Information and Modeling 64(7), 2205–2220 (2023)

119. Zheng, L., Fan, J., Mu, Y.: Onionnet: a multiple-layer intermolecular-contact-based convolutional neural network for protein–ligand binding affinity prediction. ACS omega 4(14), 15956–15965 (2019)

120. Zhou, C., Yu, H., Ding, Y., Guo, F., Gong, X.J.: Multi-scale encoding of amino acid sequences for predicting protein interactions using gradient boosting decision tree. PLoS One 12(8), e0181426 (2017)

121. Zhou, M., Li, Q., Wang, R.: Current experimental methods for characterizing protein–protein interactions. ChemMedChem 11(8), 738–756 (2016)

122. Zomorodian, A., Carlsson, G.: Computing persistent homology. In: Proceedings of the twentieth annual symposium on Computational geometry. pp. 347–356 (2004)

